# Bioinformatics analysis of potential key genes and mechanisms in type 2 diabetes mellitus

**DOI:** 10.1101/2021.03.28.437386

**Authors:** Basavaraj Vastrad, Chanabasayya Vastrad

## Abstract

Type 2 diabetes mellitus (T2DM) is etiologically related to metabolic disorder. The aim of our study was to screen out candidate genes of T2DM and to elucidate the underlying molecular mechanisms by bioinformatics methods. Expression profiling by high throughput sequencing data of GSE154126 was downloaded from Gene Expression Omnibus (GEO) database. The differentially expressed genes (DEGs) between T2DM and normal control were identified. And then, functional enrichment analyses of gene ontology (GO) and REACTOME pathway analysis was performed. Protein–protein interaction (PPI) network and module analyses were performed based on the DEGs. Additionally, potential miRNAs of hub genes were predicted by miRNet database. Transcription factors (TFs) of hub genes were detected by NetworkAnalyst database. Further, validations were performed by receiver operating characteristic curve (ROC) analysis and real-time polymerase chain reaction (RT-PCR). In total, 925 DEGs were identified in T2DM, including 447 up regulated genes and 478 down-regulated genes. Functional enrichment analysis results showed that up regulated DEGs were significantly enriched in defense response, neutrophil degranulation, cell adhesion and extracellular matrix organization. The top 10 hub genes, JUN, VCAM1, RELA, U2AF2, ADRB2, FN1, CDK1, TK1, A2M and ACTA2 were identified from the PPI network, modules, miRNA-hub gene regulatory network and TF-hub gene regulatory network. Furthermore, ROC analysis and RT-PCR revealed that JUN, VCAM1, RELA, U2AF2, ADRB2, FN1, CDK1, TK1, A2M and ACTA2 might serve as biomarkers in T2DM. Bioinformatics analysis is a useful tool to explore the molecular mechanism and pathogenesis of T2DM. The identified hub genes may participate in the onset and advancement of T2DM and serve as therapeutic targets.

## Introduction

Type 2 diabetes mellitus (T2DM) is the most common chronic metabolic disorder in adults and its affects more than 90% people worldwide [1]. The incidence rate of T2DM is high, which vigorously affects the patient’s health. T2DM can affect the brain [2], kidney [3], eyes [4] and heart [5]. Numerous risk factors might affect T2DM progression, including genetic [6], environmental [7] metabolic [8] risk factors, obesity [9] and hypertension [10]; however, how these factors affect the progression of T2DM requires further investigation and no effective treatment has been advanced for the relief of T2DM. In addition, molecular biology investigation have identified numerous biomarkers and signaling pathways that contribute to T2DM, including the OCT1 [11], TNF-α [12], TCF7L2 [13], KCNJ11 [14], SDF-1β [14], EDNRB [15], NOS3 [15], PI3K/AKT pathway [16], IL-6 signalling pathways [17], Wnt signaling pathway [18], insulin signaling [19] and NF-κB signaling pathway [20]. Further investigation into the molecular events linked with T2DM progression is required.

With the advancement of molecular biological and pathological techniques, a range of markers are considered powerful tools for searching diagnostic and predictive biomarkers, especially in the therapies targeting T2DM. Expression profiling by high throughput sequencing, efficient and large-scale techniques for gaining genetic data, have been widely performed to collect and to study Expression profiling by high throughput sequencing data in T2DM. These high throughput sequencing studies develop a novel approach for identifying T2DM related genes and implement encouraging expectation for molecular prediction and therapeutic targets [21].

In the current investigation, the expression profiling by high throughput sequencing data of GSE154126 [22] was downloaded from Gene Expression Omnibus (GEO, http://www.ncbi.nlm.nih.gov/geo/) [23] to screen the differentially expressed genes (DEGs) between T2DM and normal control. Next, the DEGs were analyzed using a bioinformatics approach. To identify the enriched biological functions and pathways of DEGs, the Gene Ontology (GO) and REACTOME pathway enrichment analysis of DEGs were performed. Subsequently, functional enrichment, protein-protein interaction (PPI) networks construction, module analyses and miRNA-hub regulatory gene network construction and TF-hub regulatory gene network construction were performed to determine the significant pathogenic genes and their key pathways involved in the mechanism of occurrence and progression of T2DM. Hub genes were validated using the receiver operating characteristic curve (ROC) analysis and RT-PCR. These results may provide information for subsequent experimental studies of T2DM, and contribute to the understanding of the molecular mechanisms underlying the advancement of T2DM.

## Material and Methods

### RNA sequencing data

The expression profiling by high throughput sequencing of GSE154126 was downloaded from the GEO database. The data were produced based on the GPL11154 platform Illumina HiSeq 2000 (Homo sapiens). The GSE154126 dataset contained 1276 samples, including 508 samples of T2DM samples and 768 normal control samples.

### Identification of DEGs

The limma package [24] in R Bioconductor was used to identify DEGs between T2DM samples and normal control samples. The P-values of DEGs were calculated using a t-test in R with the limma package. P value <0.05, |logFC| > 0.63 for up regulated genes and |logFC| < −1.99 for down regulated genes were set as the cutoff criterion for statistically significant DEGs. In this investigation, ggplot2 and the gplots in R package were used to plot the volcano diagrams and heat map of the DEGs, respectively.

### Gene Ontology (GO) and REACTOME pathway enrichment analysis

Gene Ontology (GO) (http://geneontology.org/) [25] and REACTOME (https://reactome.org/) [26] pathway enrichment analyses of T2DM DEGs were performed using ToppGene (ToppFun) (https://toppgene.cchmc.org/enrichment.jsp) [27]. GO terms of biological processes (BP), cellular components (CC), and molecular functions (MF) linked with a p-value <0.05 were considered to be significantly enriched.

### PPI network construction and module analysis

The online IntAct interactome (https://www.ebi.ac.uk/intact/) [28] database was used to identify potential interaction among the common DEGs, and a confidence score of ≥0.4 was set as the threshold. Cytoscape software, version 3.8.2 (www.cytape.org) [29] was used to visualize the PPI network of common DEGs. The Network Analyzer plug-in was used to calculate node degree [30], betweenness centrality [31], stress centrality [32] and closeness centrality [33], that were to filter hub genes of PPI. The PEWCC1 (http://apps.cytoscape.org/apps/PEWCC1) [34] was used to search modules of the PPI network and the default parameters (Degree cutoff ≥10, node score cutoff ≥0.4, K-core ≥4, and max depth=100.) were set in the functional interface of Cytoscape software.

### MiRNA-hub gene regulatory network construction

miRNet database (https://www.mirnet.ca/) [35] for miRNA-hub gene regulations, through which regulatory associations between hub genes and miRNAs can be identified. This study inputted the overlapped hub genes into the database to examine the regulatory association pairs between miRNAs and hub genes. Based on the data this study obtained, miRNA-hub gene regulatory network was constructed and visualized by Cytoscape 3.8.2 software to show the overlapped miRNAs and hub genes. Therefore, these miRNAs and hub genes might play a potential role in the pathogenesis and treatment of T2DM.

### TF-hub gene regulatory network construction

NetworkAnalyst database (https://www.networkanalyst.ca/) [36] for TF-hub gene regulations, through which regulatory associations between hub genes and TFs can be identified. This study inputted the overlapped hub genes into the database to examine the regulatory association pairs between TFs and hub genes. Based on the data this study obtained, TF -hub gene regulatory network was constructed and visualized by Cytoscape 3.8.2 software to show the overlapped TFs and hub genes. Therefore, these TFs and hub genes might play a potential role in the pathogenesis and treatment of T2DM.

### Validation of hub genes by receiver operating characteristic curve (ROC) analysis

Genes in the PPI network identified by Network Analyzer plug-in were selected as candidate hub genes. To evaluate the role of candidate genes in the diagnosis of T2DM, receiver operating characteristic (ROC) curve analysis was organize in RStudio with pROC package [37]. The genes with area under curve (AUC) >0.8 was considered as hub genes of T2DM.

### Detection of the mRNA expression of the hub genes by RT-PCR

Pancreatic beta cell line MIN6 culture treated with streptozotocin for T2DM and pancreatic beta cell line MIN6 culture for normal control were used to examine the mRNA expression of the hub genes in T2DM and normal control. Pancreatic beta cell line MIN6 were maintained in 15 % fetal calf serum, 50 mg/l streptomycin and 75 mg/l penicillin sulphate in Dulbecco’s modified Eagle’s medium (DMEM). Pancreatic beta cell line MIN6 were cultured at 37 °C and CO_2_ concentration is 5%. Total RNA was isolated from pancreatic beta cell line MIN6 culture treated with streptozotocin and pancreatic beta cell line MIN6 culture using TRI Reagent (Sigma, USA) according to the instructions. RNA samples were reverse transcribed into cDNA by reverse transcription kit (Thermo Fisher Scientific, Waltham, MA, USA). RT-PCR analysis was performed QuantStudio 7 Flex real-time PCR system (Thermo Fisher Scientific, Waltham, MA, USA). The set of primers used in the investigation is listed in Table 1.The experiment was repeated in triplicate. PCR profiles used in this investigation were: denaturing for 10 min at 95°C, annealing for 15 sec at 95°C, and extension for 10 sec at 72°C. PCR was carried out for 30 cycles. The relative value of the hub gene was normalized to those of β-actin through the 2^-ΔΔCt^ calculation [38].

**Table 1.**
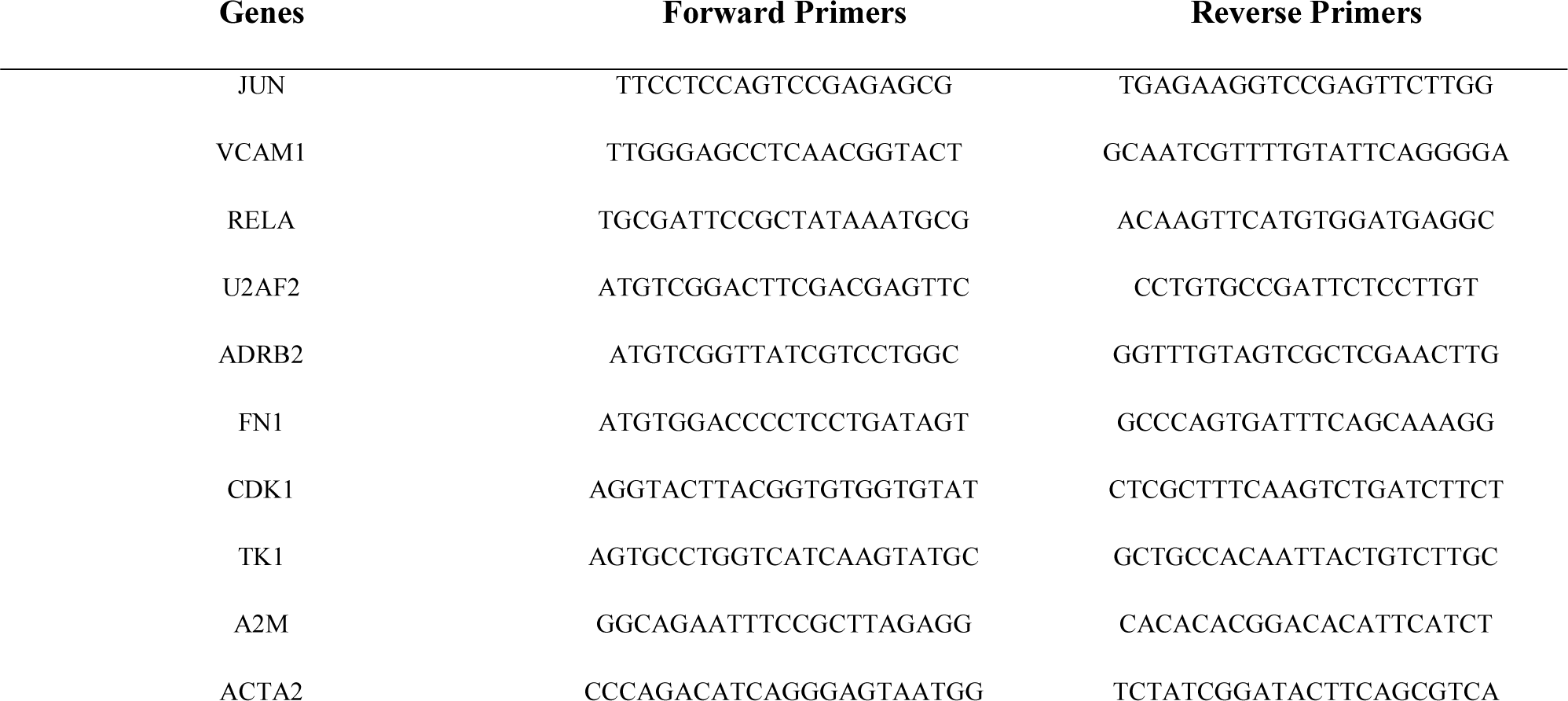
The sequences of primers for quantitative RT-PCR

## Results

### Identification of DEGs

Standardization of the expression profiling by high throughput sequencing results from GSE154126, DEGs were identified. A total of 925 differentially expressed genes including 447 up regulated genes and 478 down regulated genes were detected and are listed in Table 2. The results were validated with a volcano plot of all up regulated genes and down regulated genes (Fig.1). Fig. 2 shows the DEG expression heat map.

**Fig. 1.**
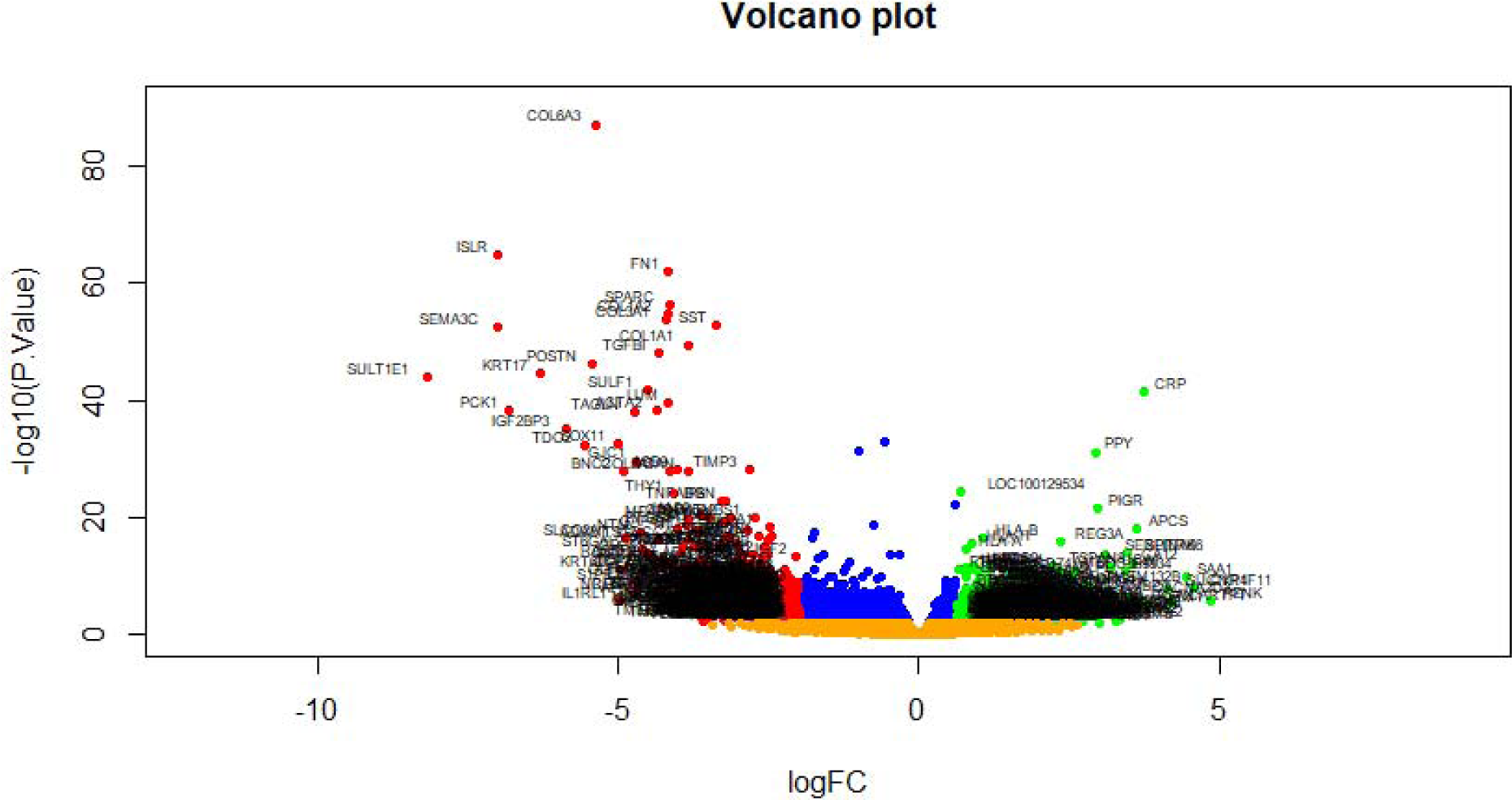
Volcano plot of differentially expressed genes. Genes with a significant change of more than two-fold were selected. Green dot represented up regulated significant genes and red dot represented down regulated significant genes.

**Fig. 2.**
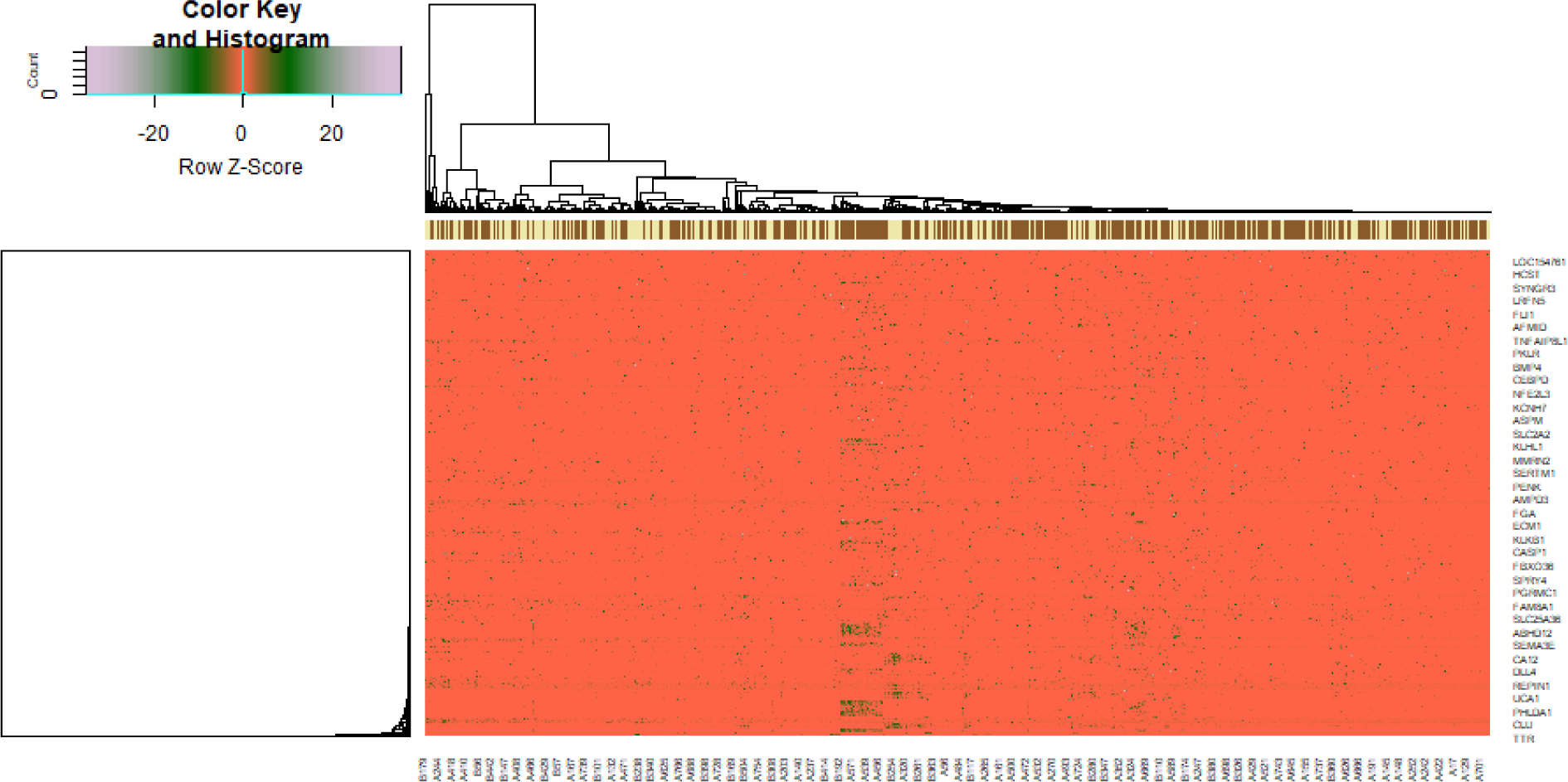
Heat map of differentially expressed genes. Legend on the top left indicate log fold change of genes. (A1 – A508= normal control; B1 – B508 = T2DM)

**Table 2.**
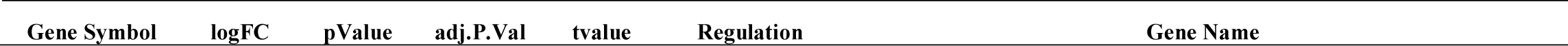
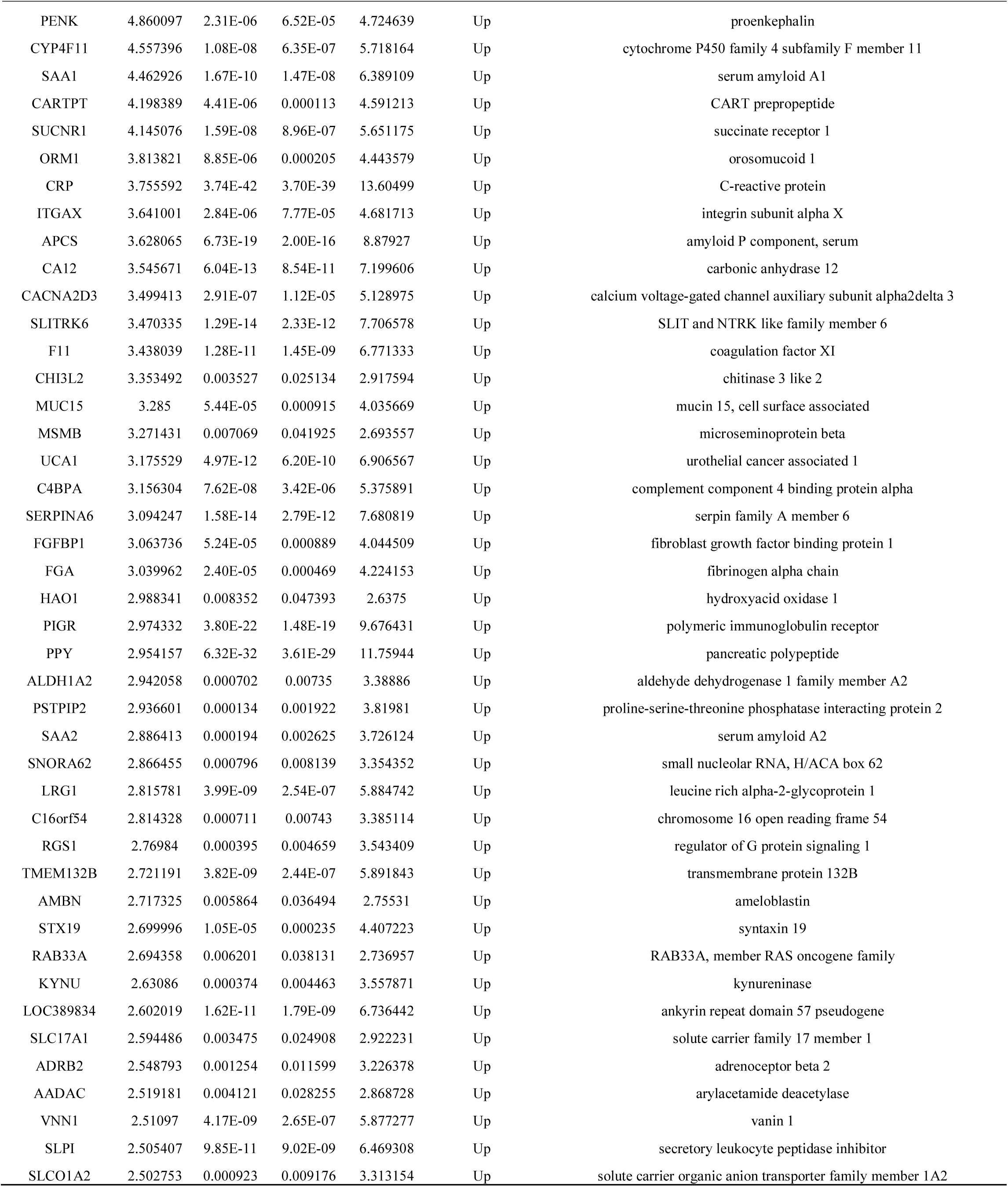
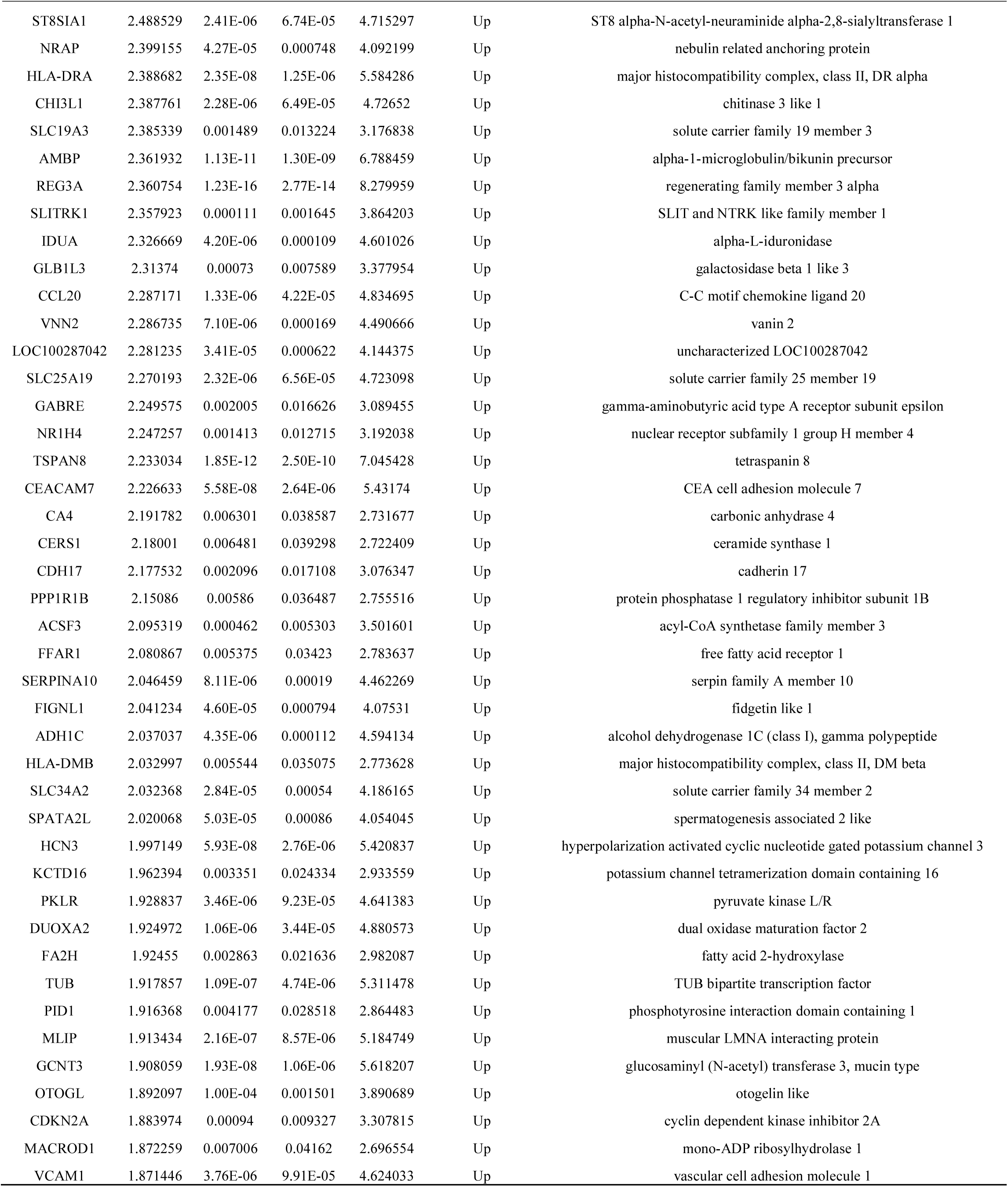
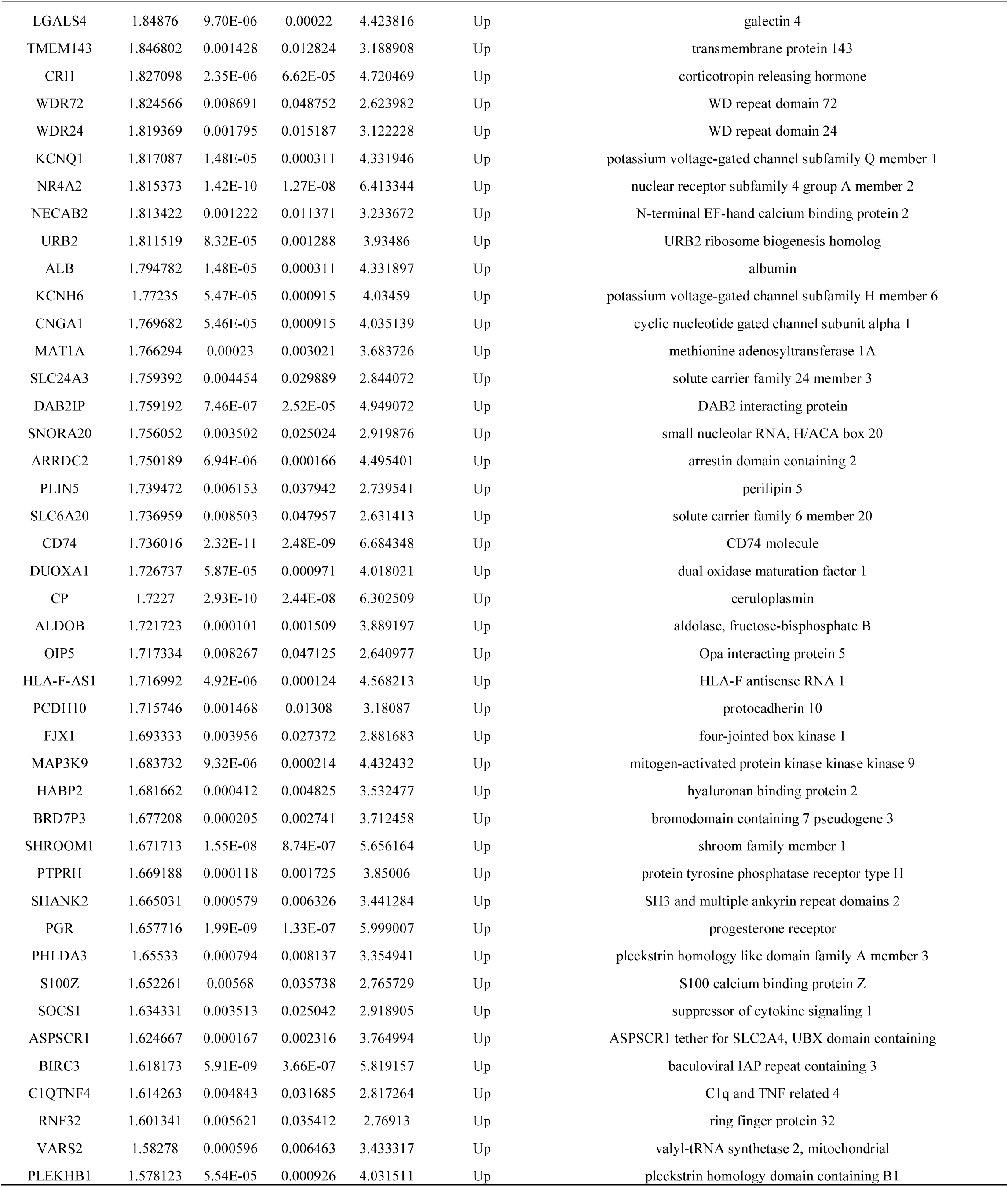
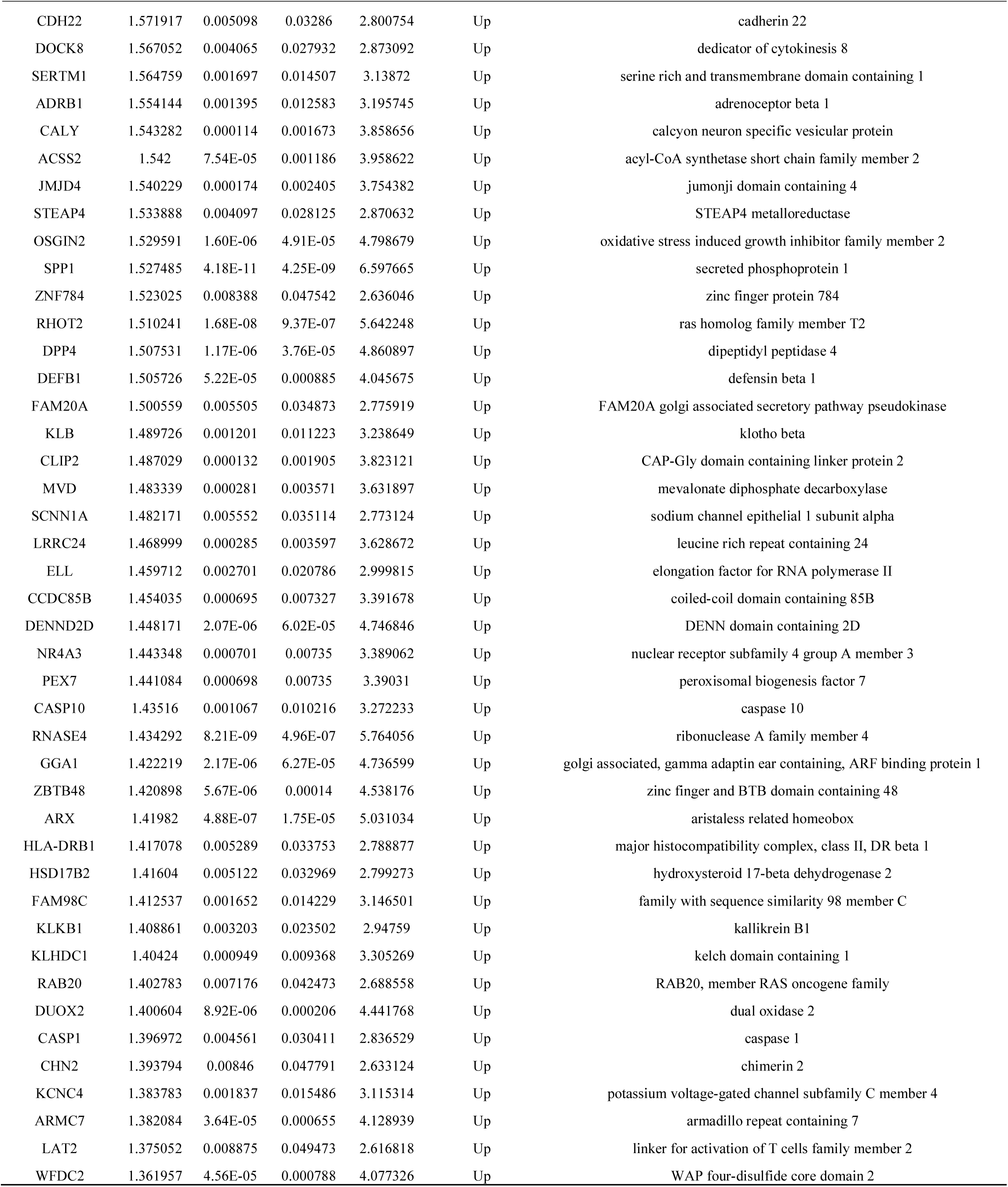
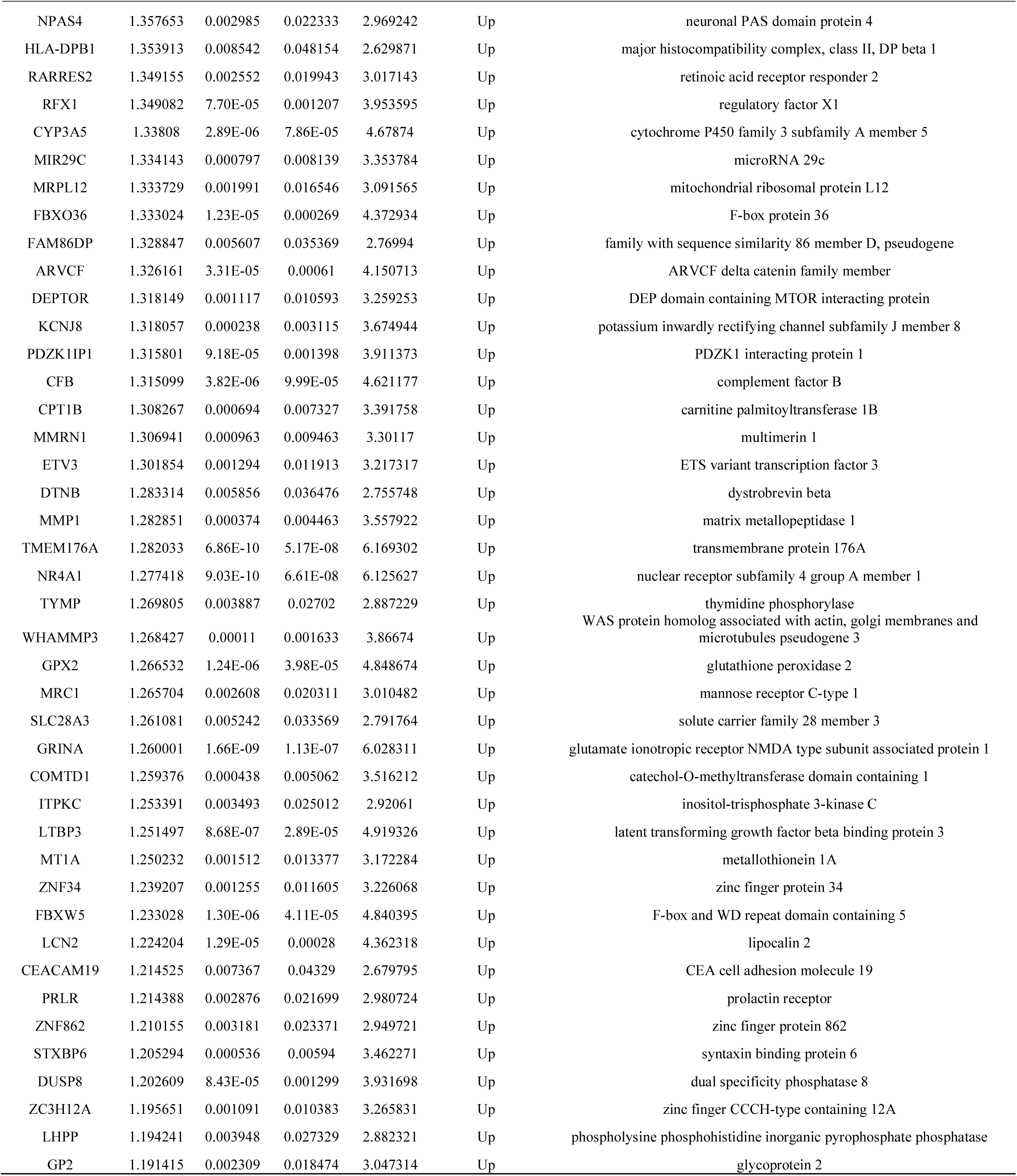
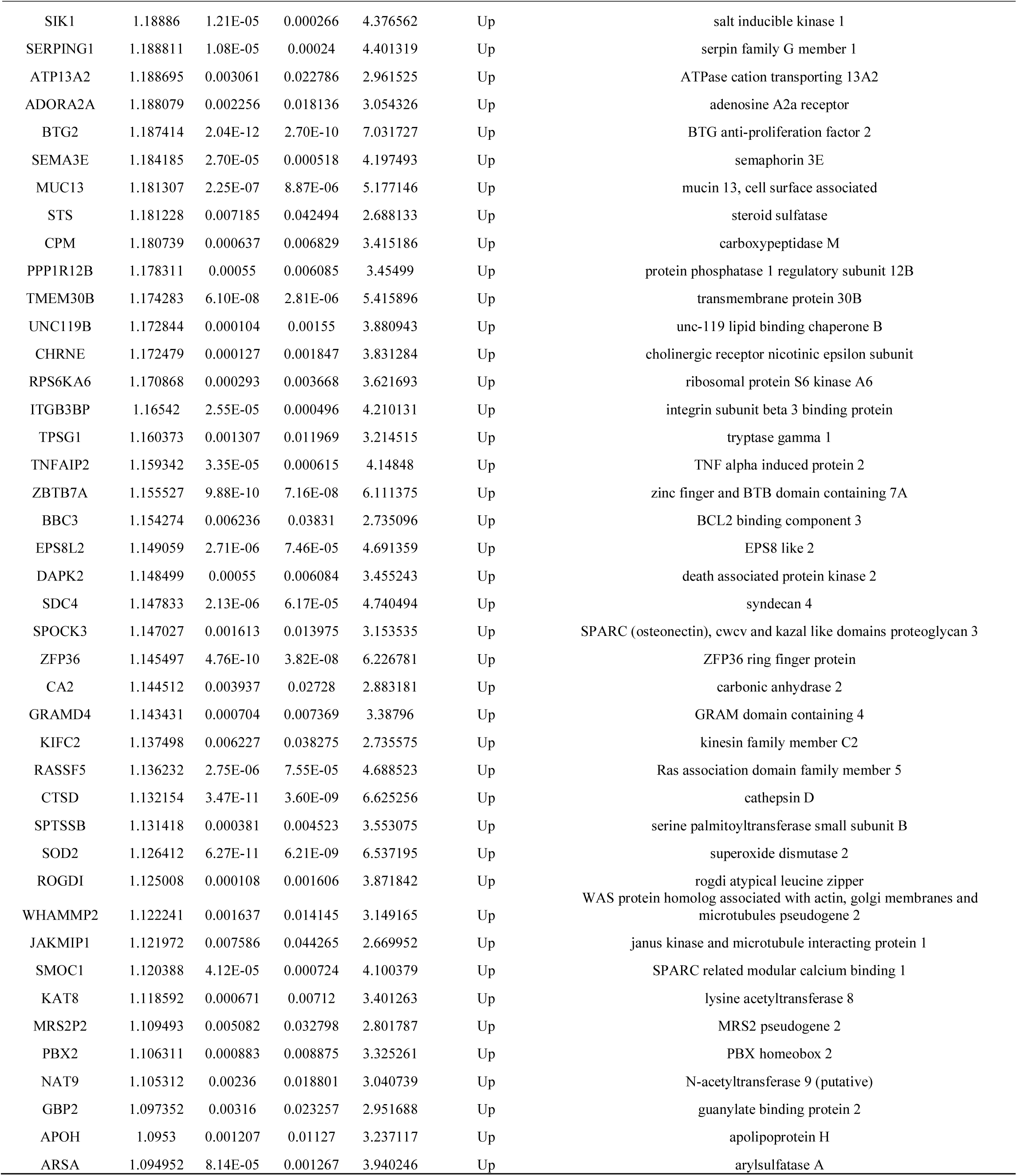
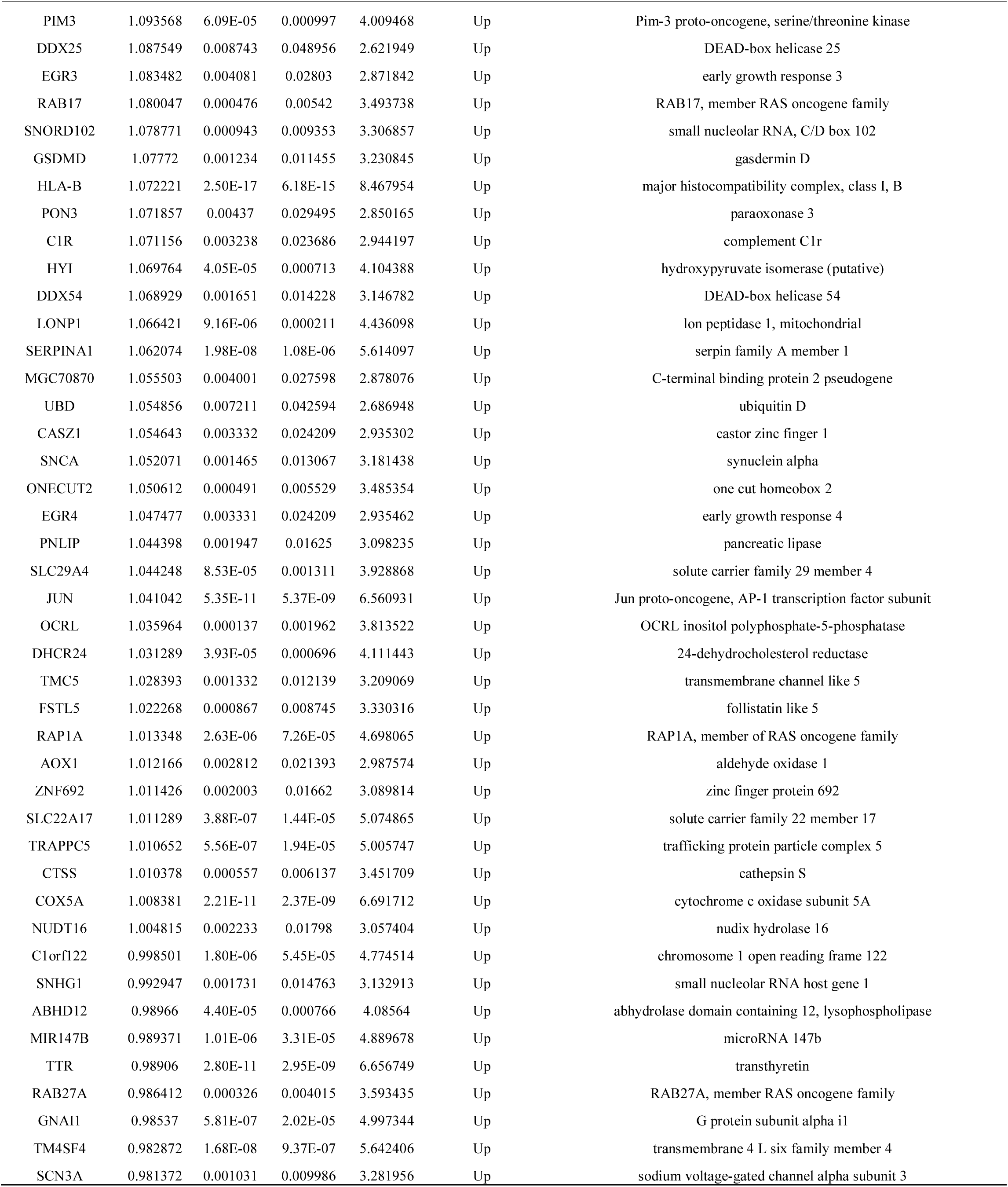
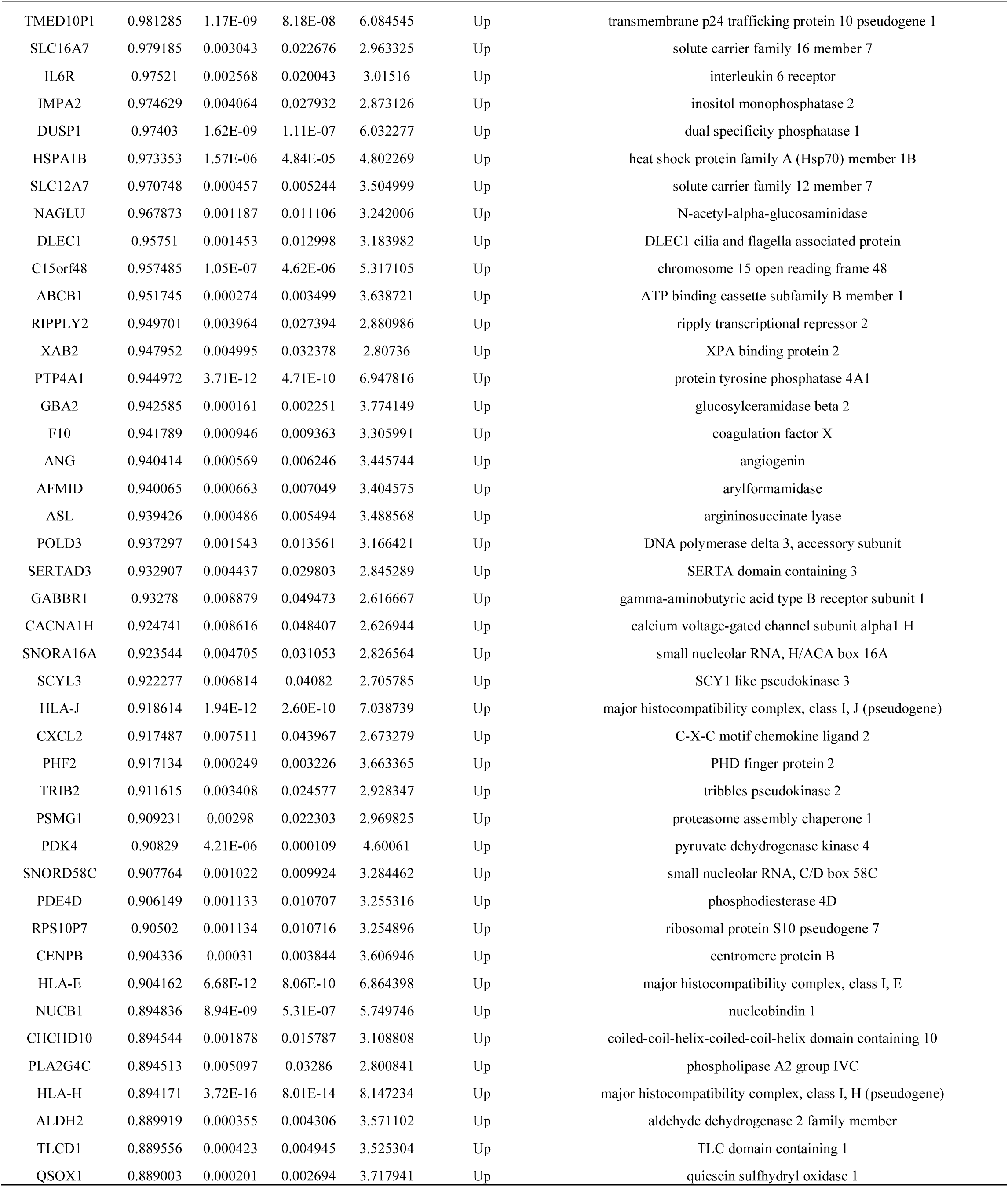
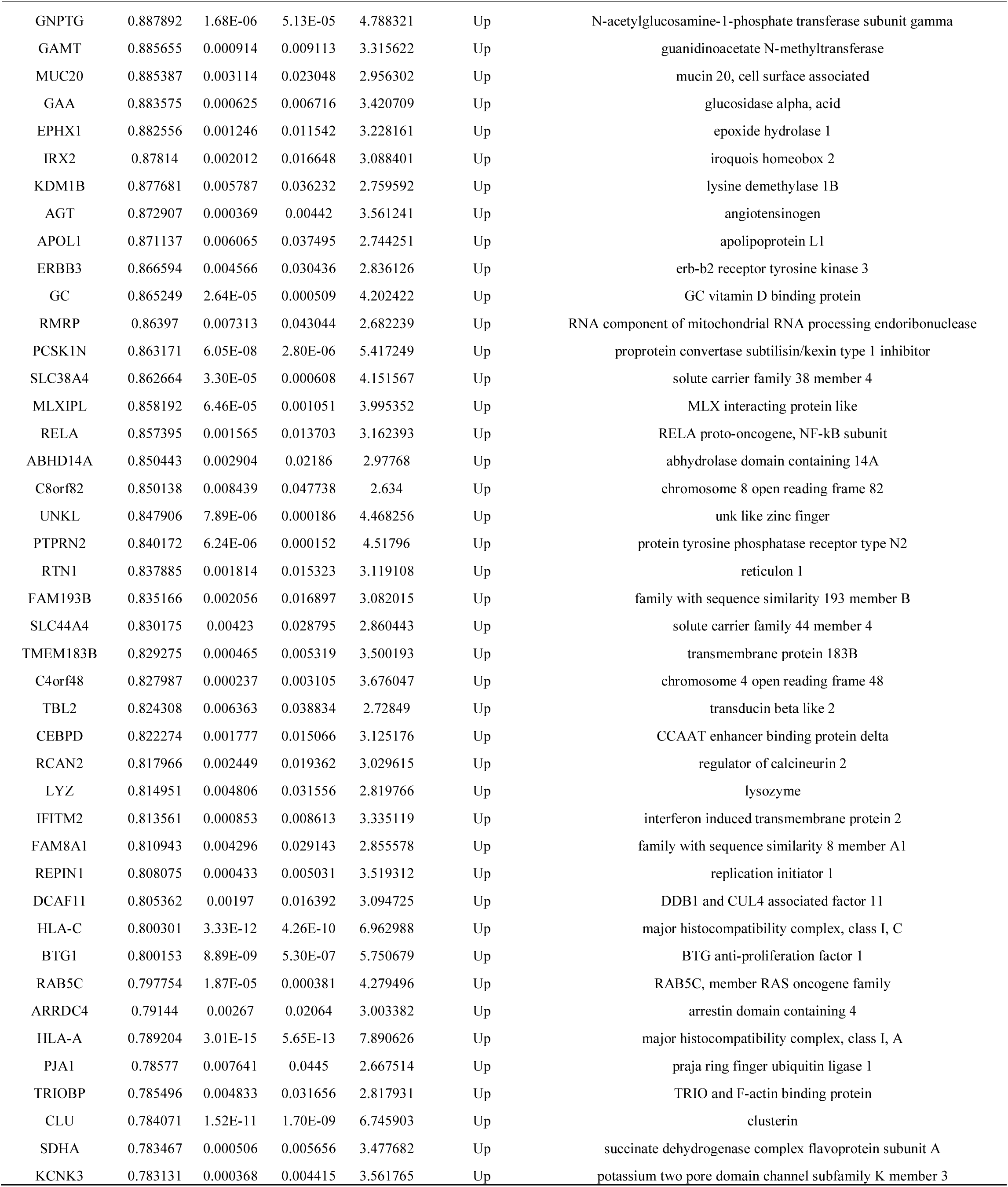
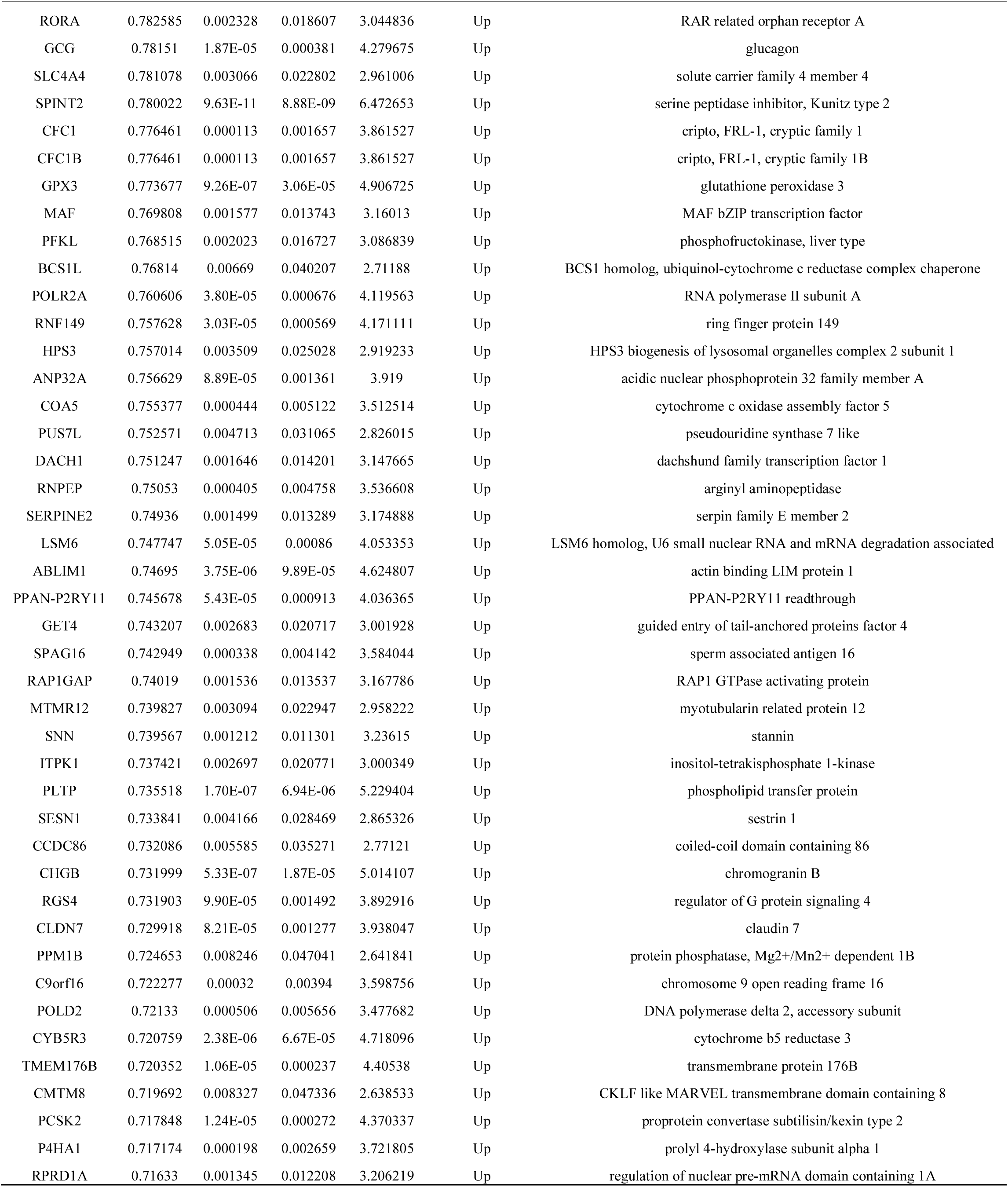
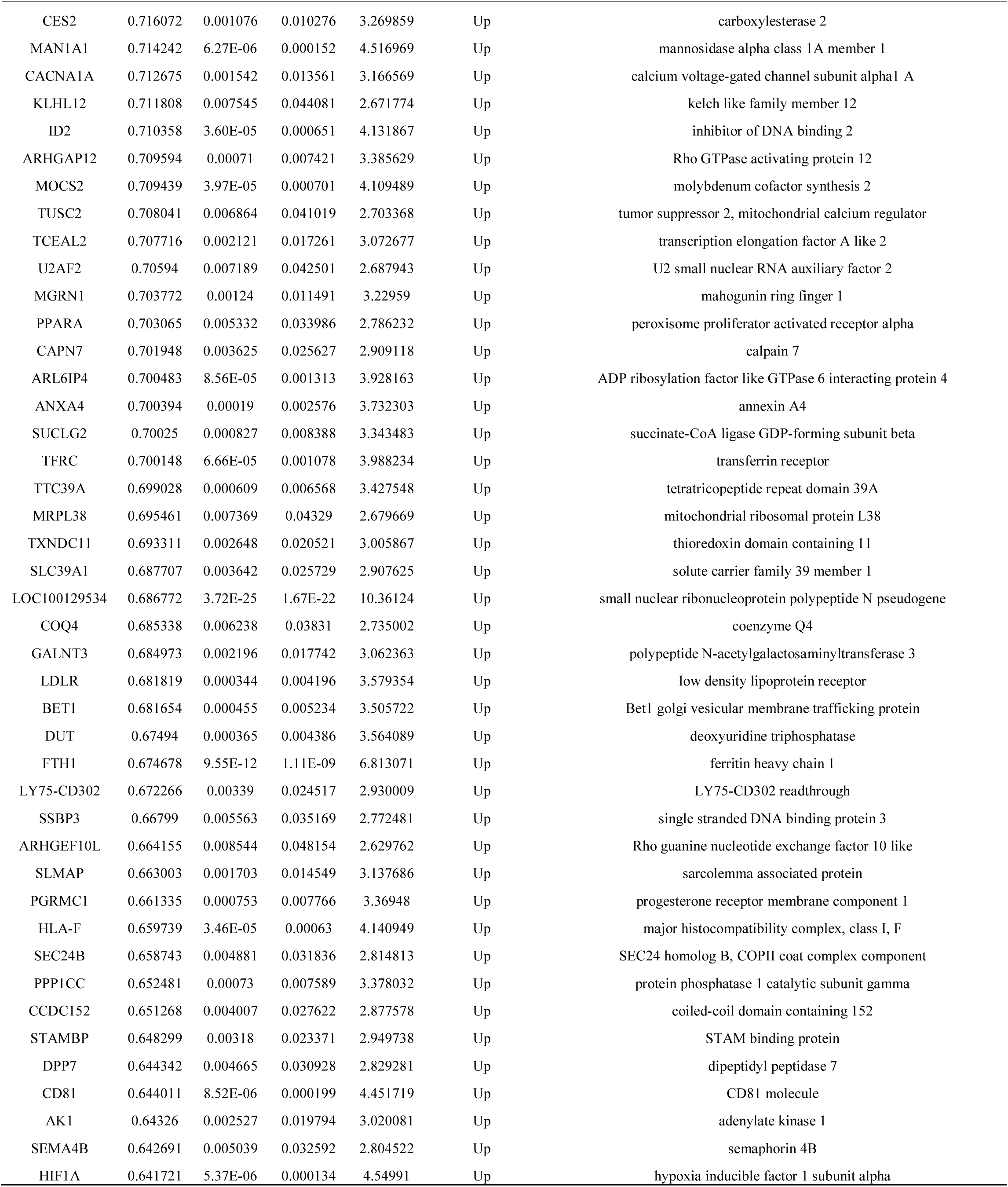
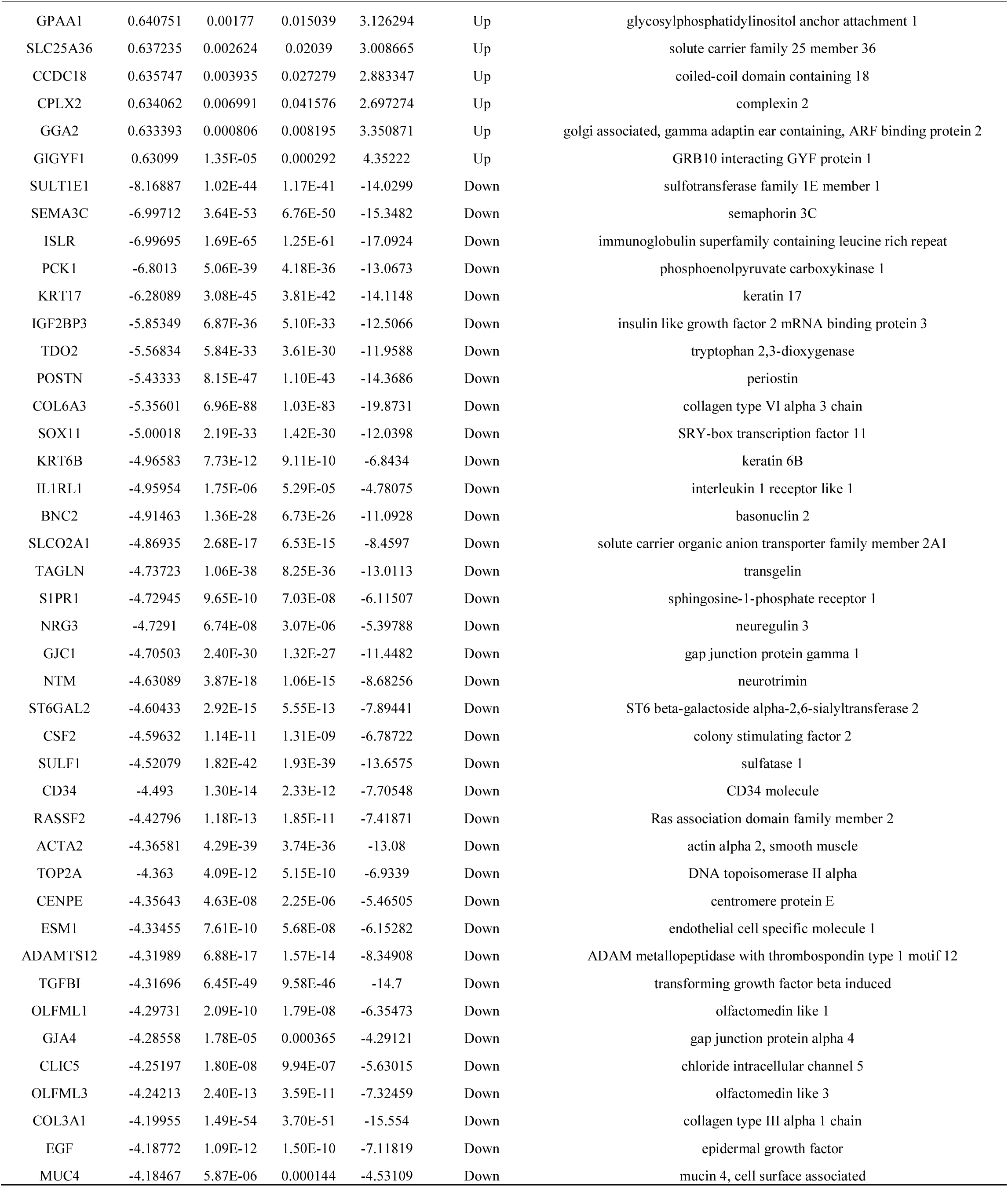
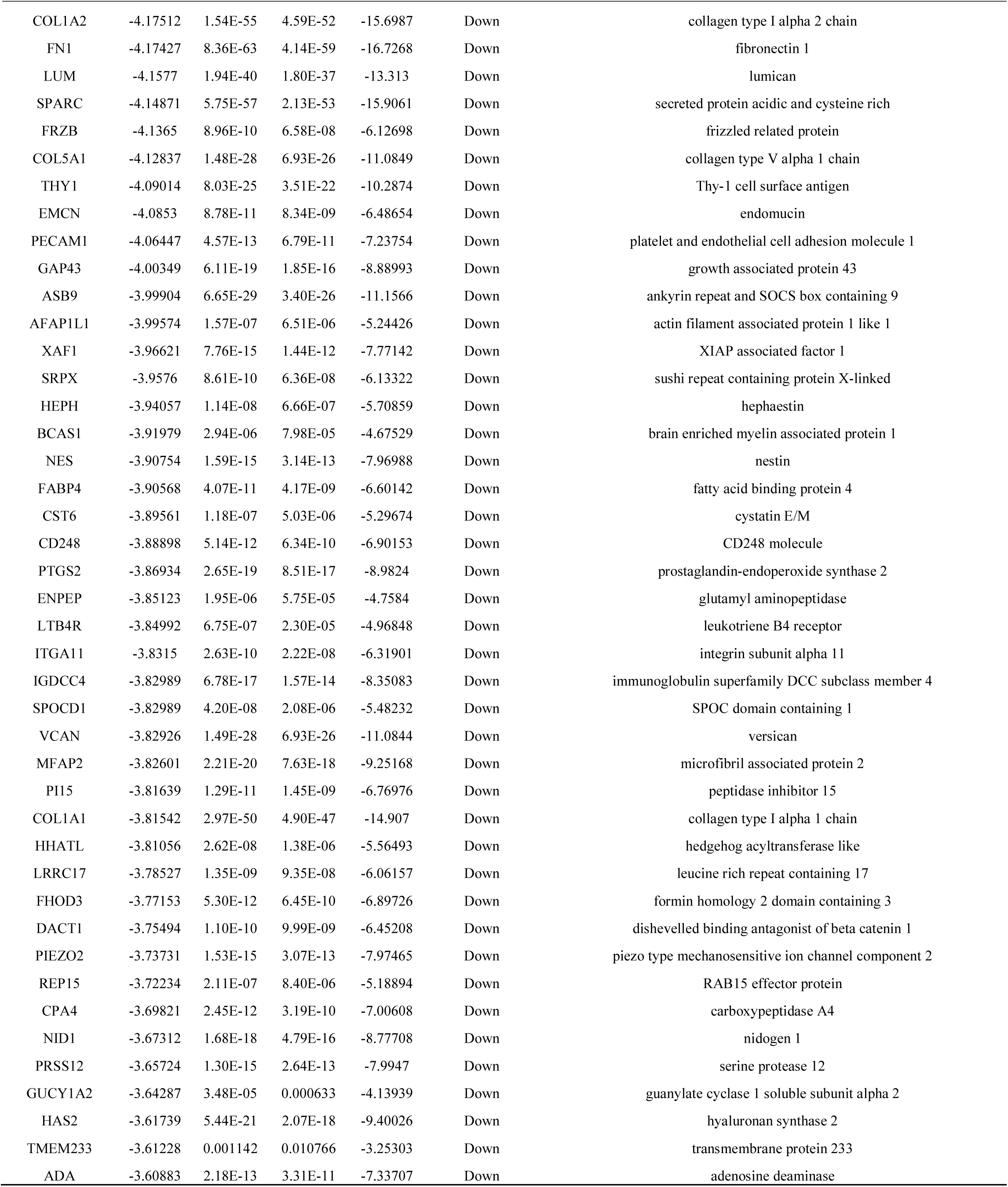
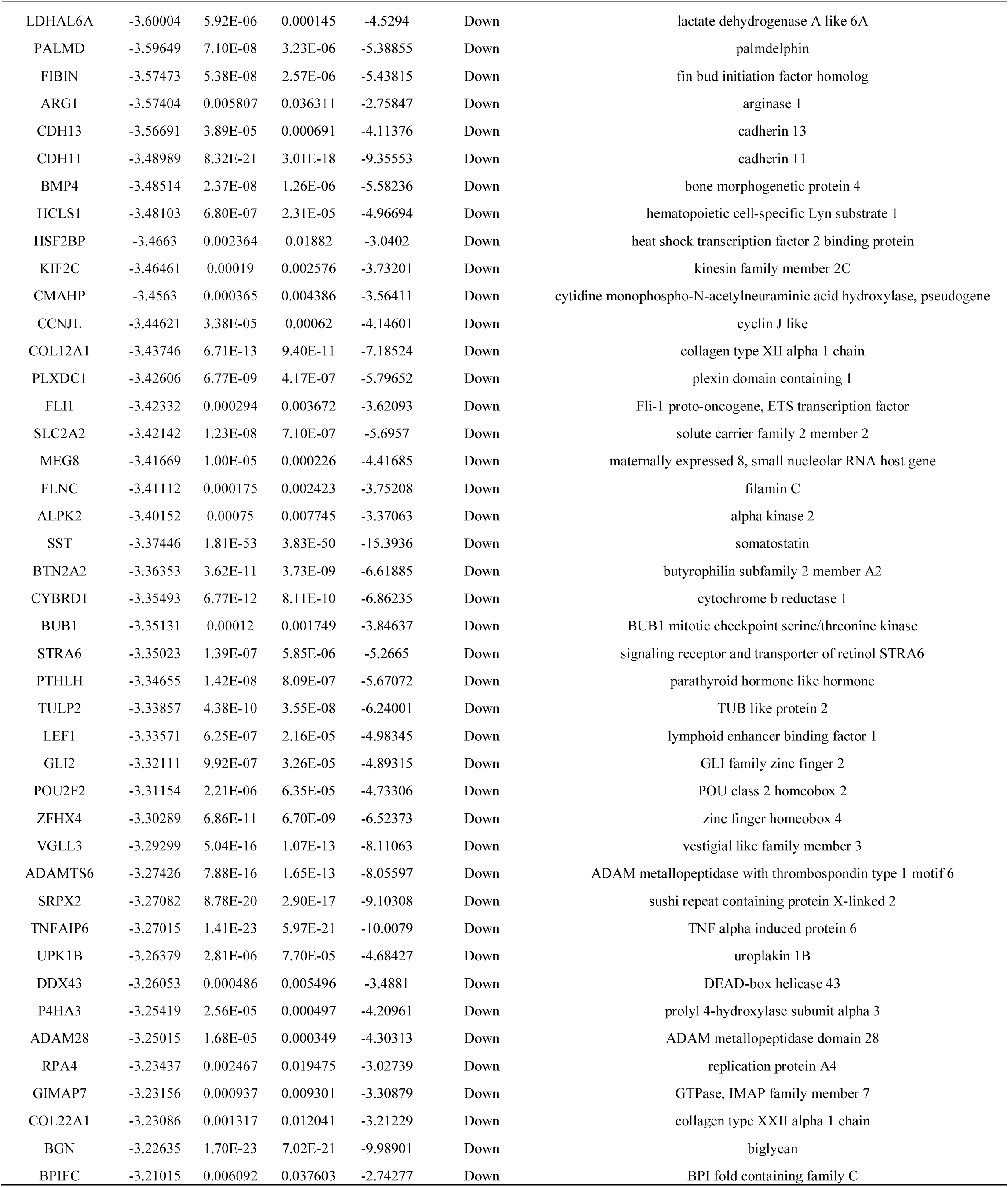
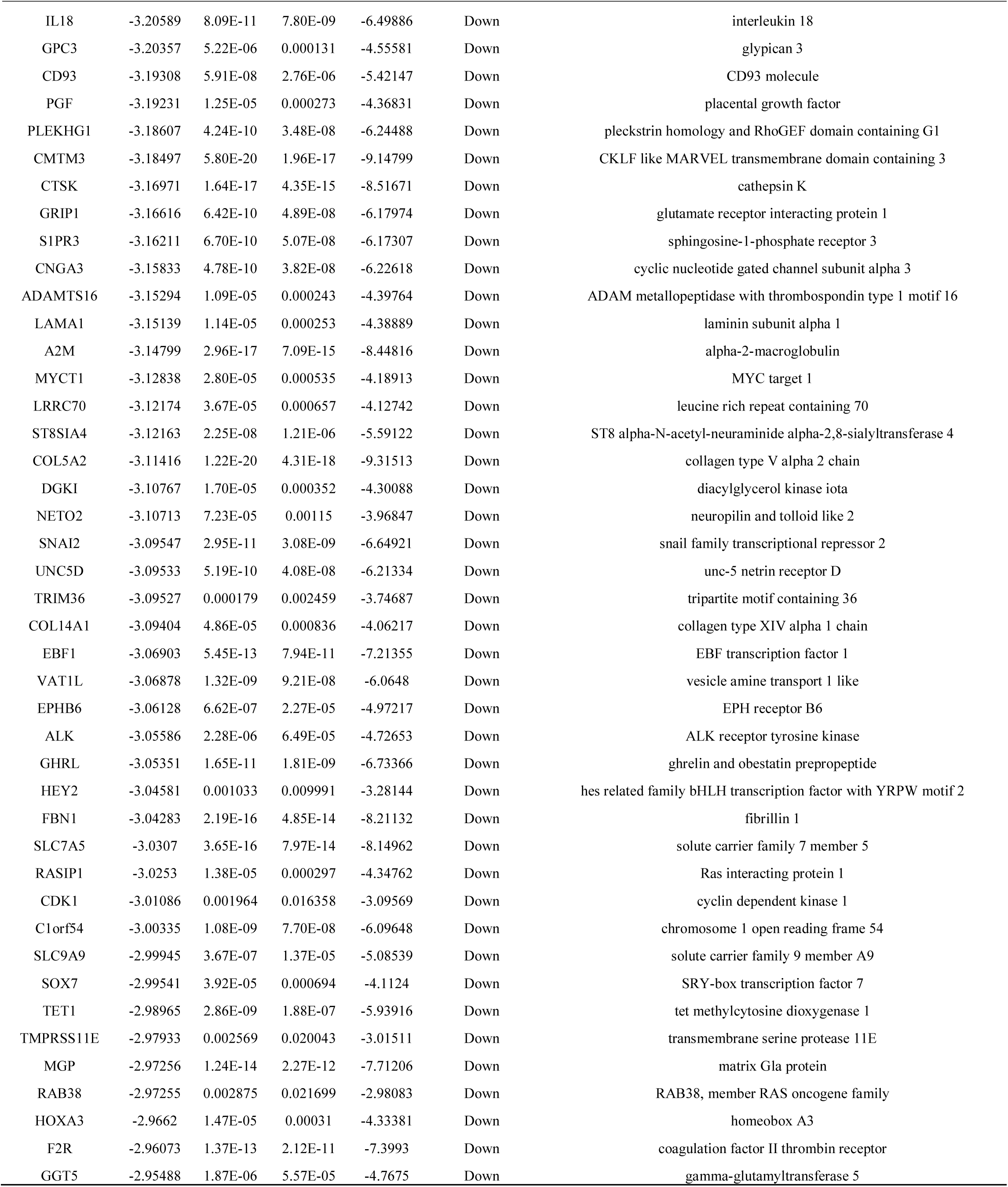
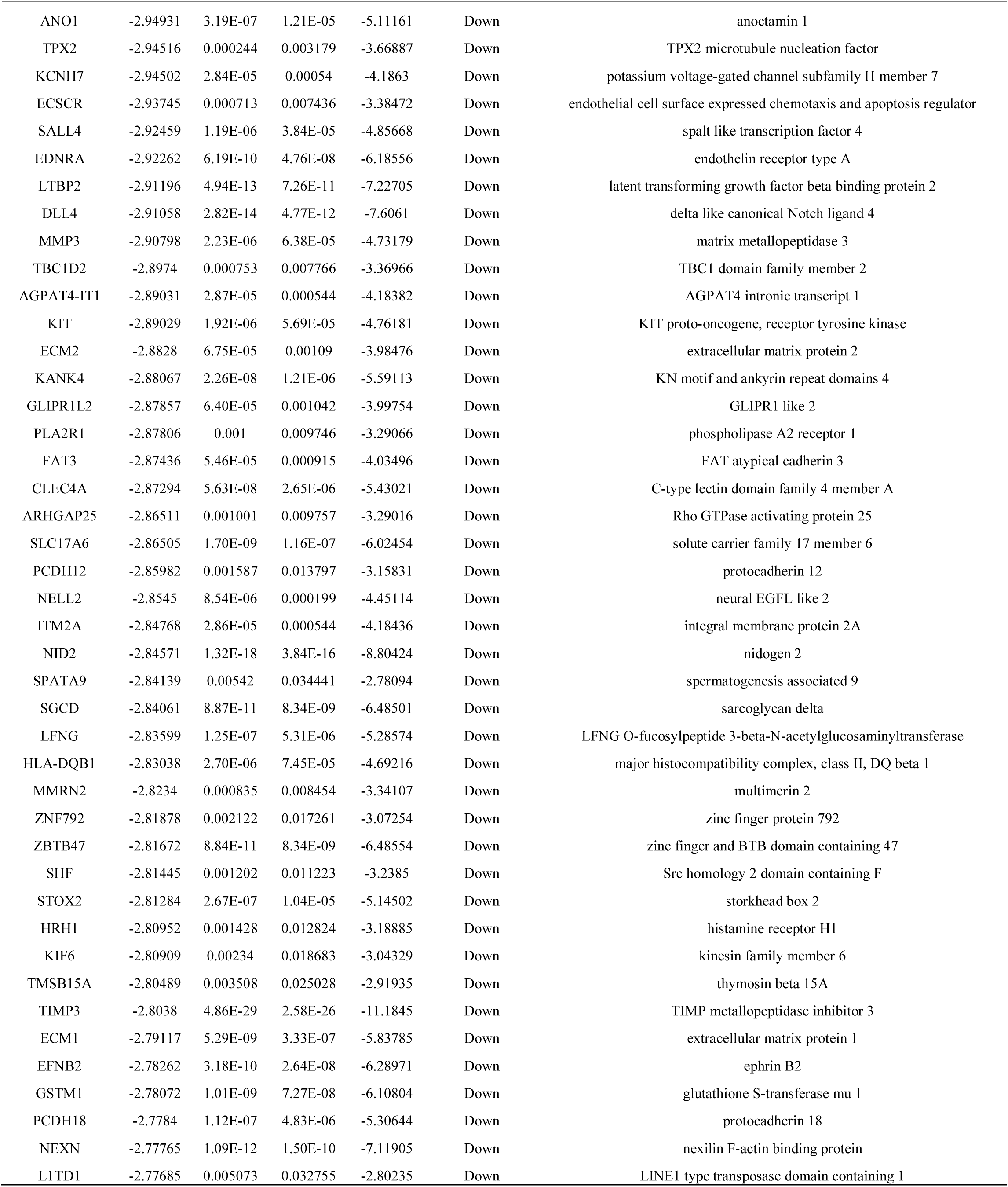
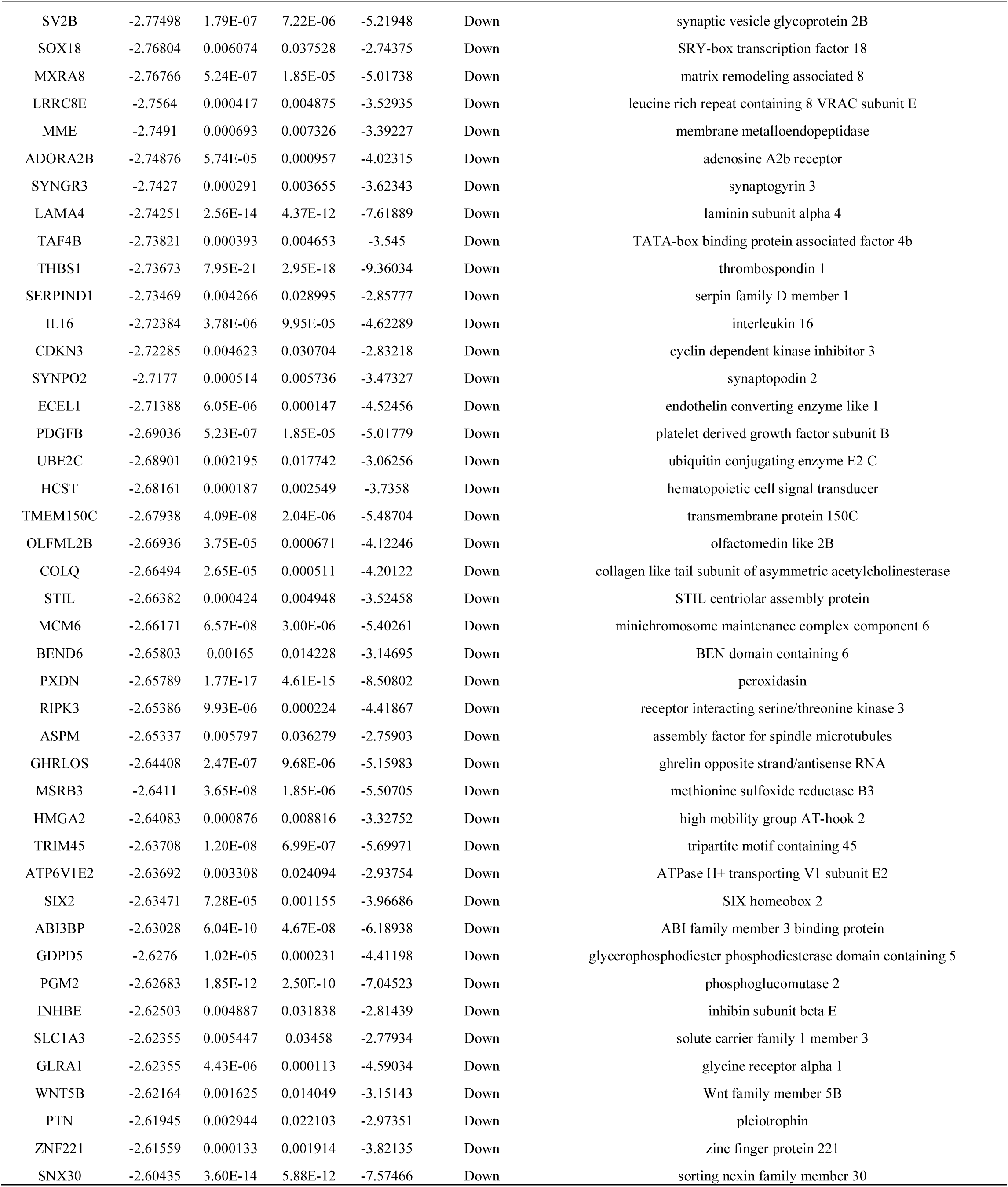
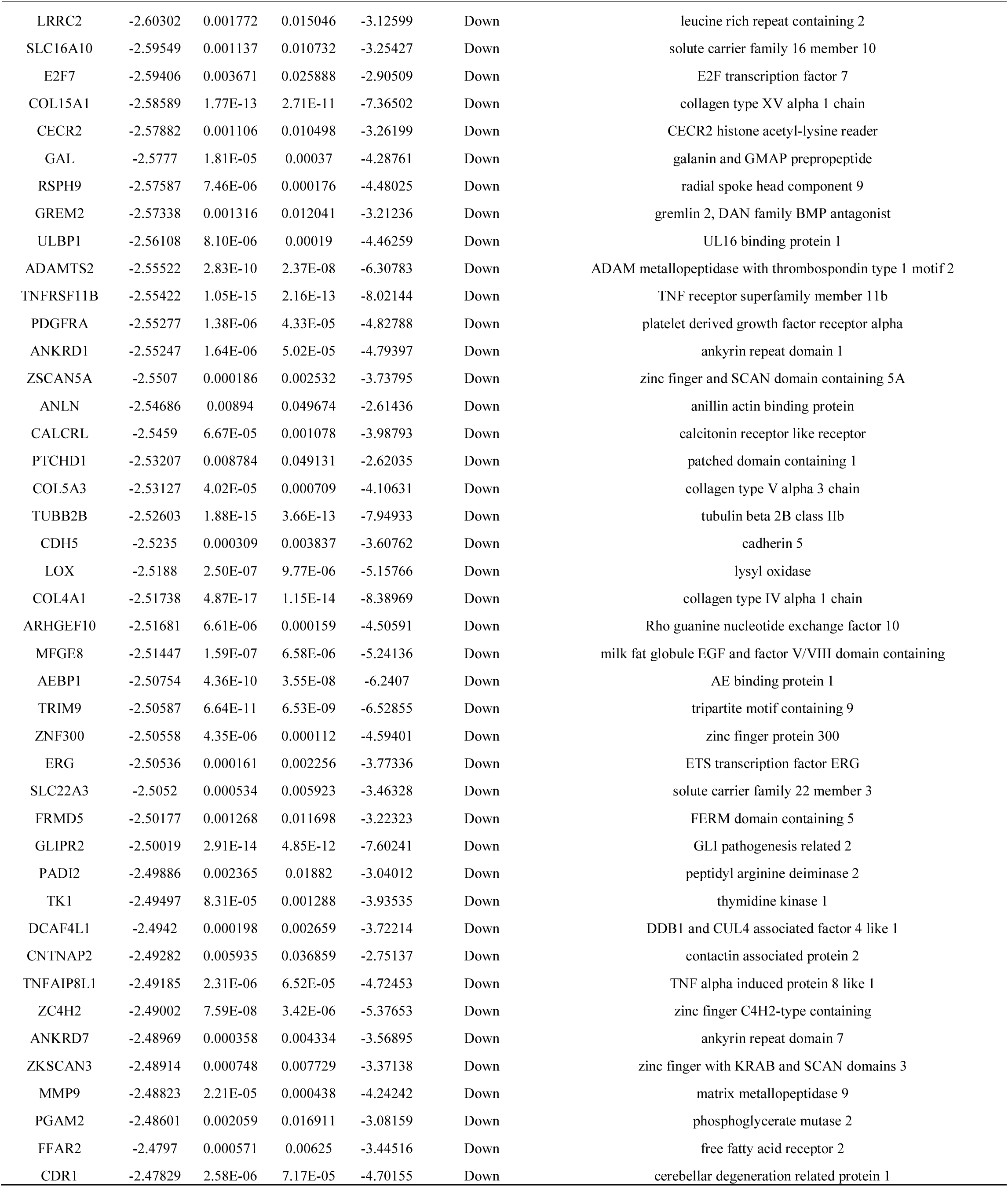
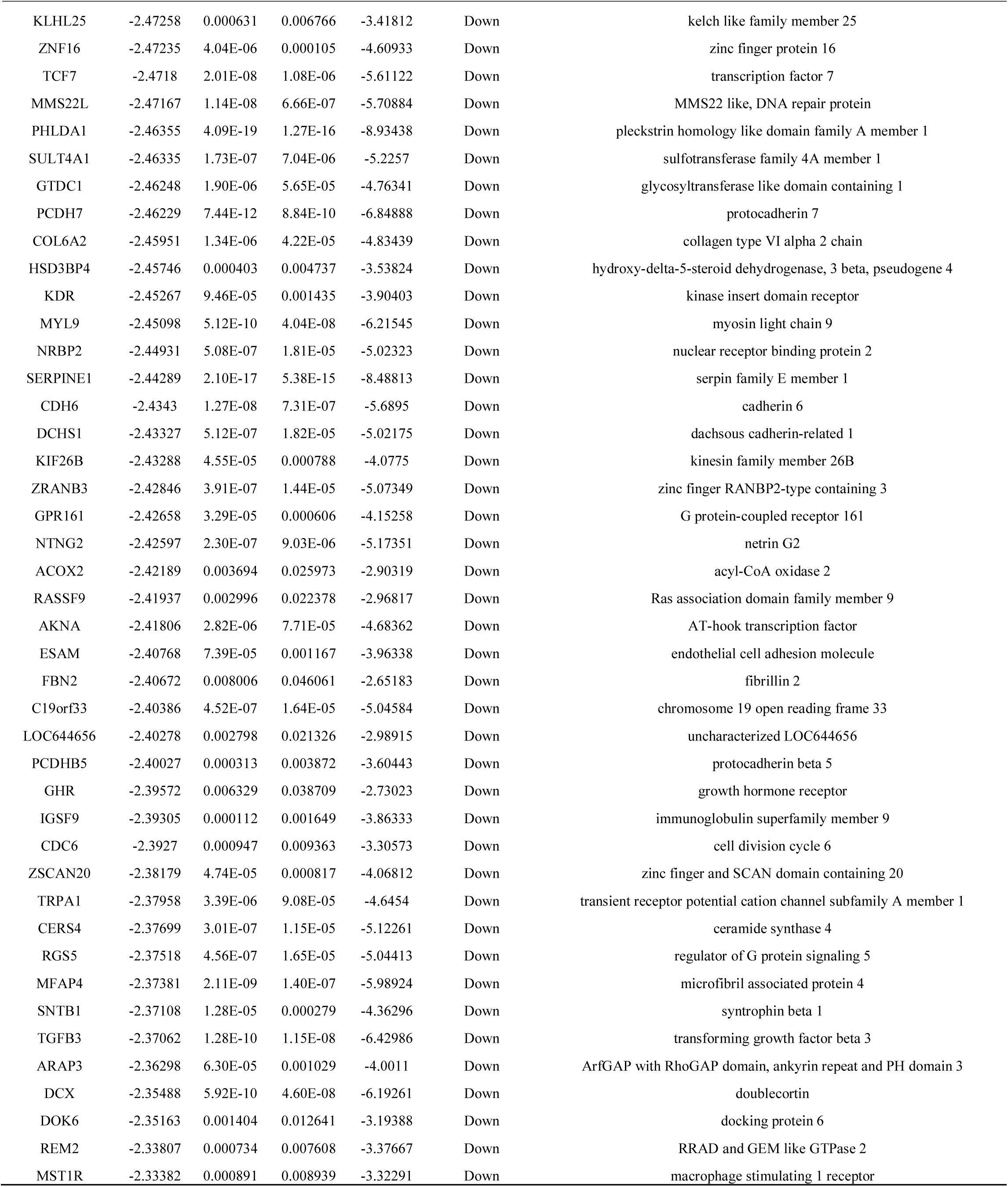
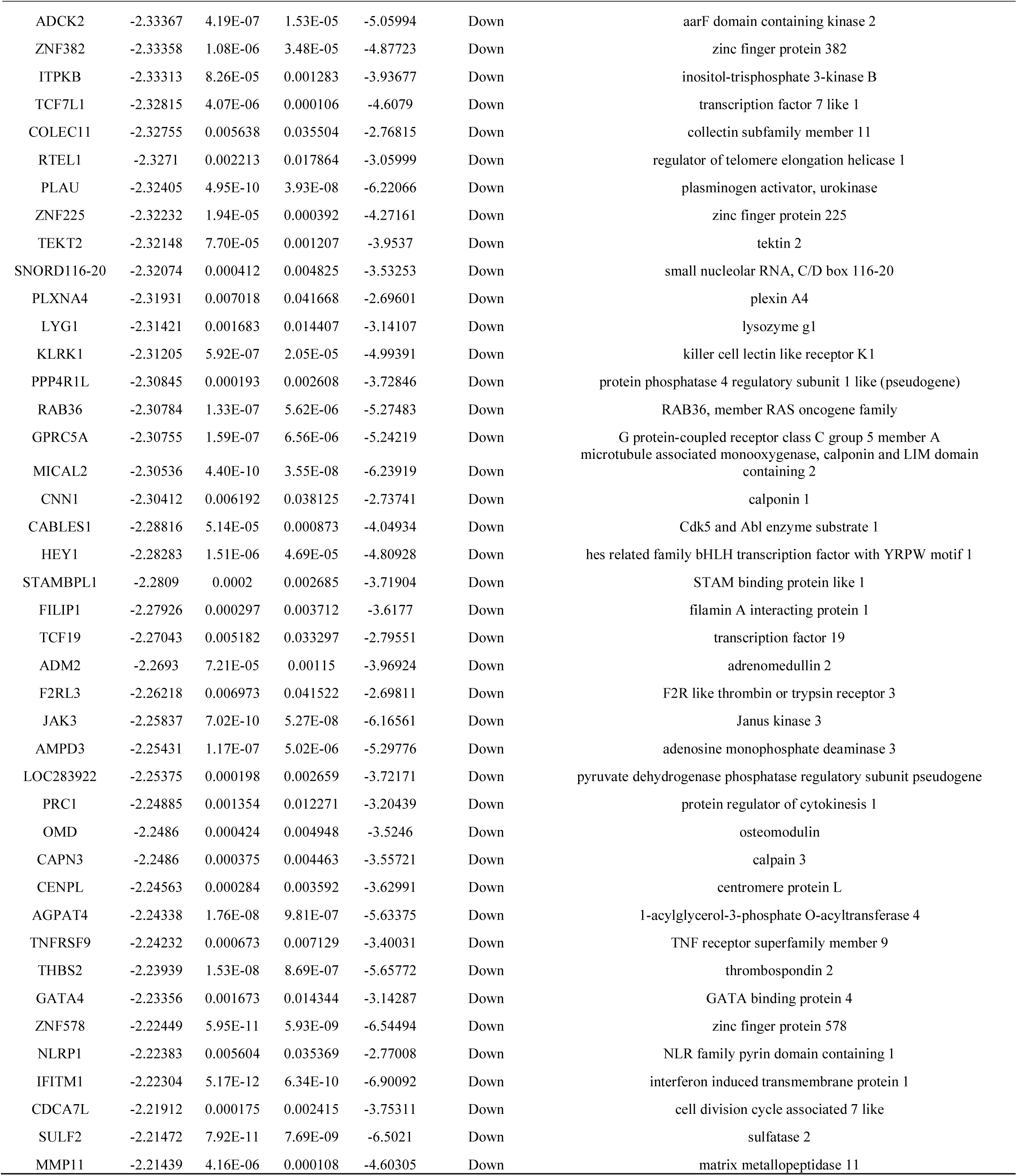
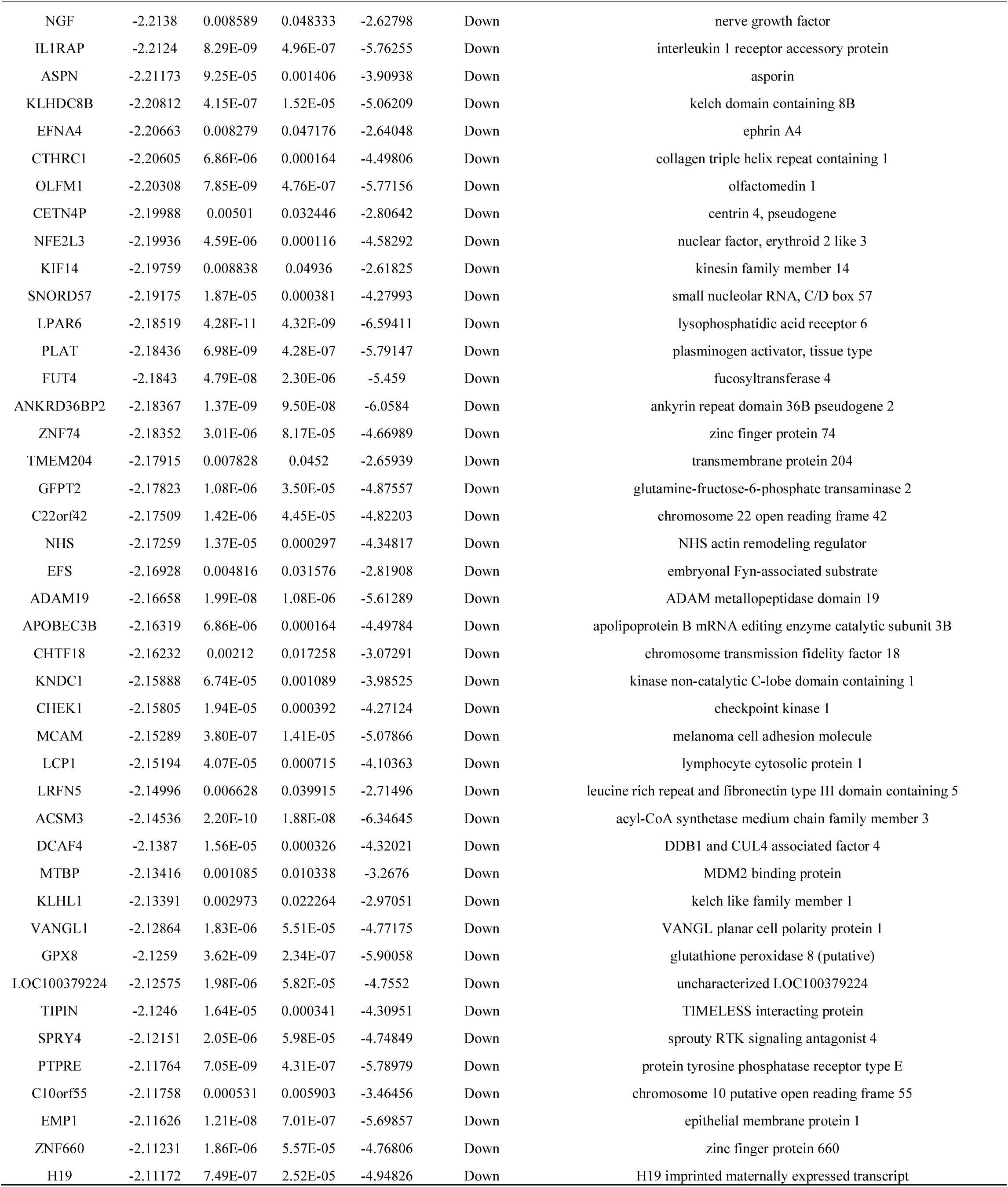
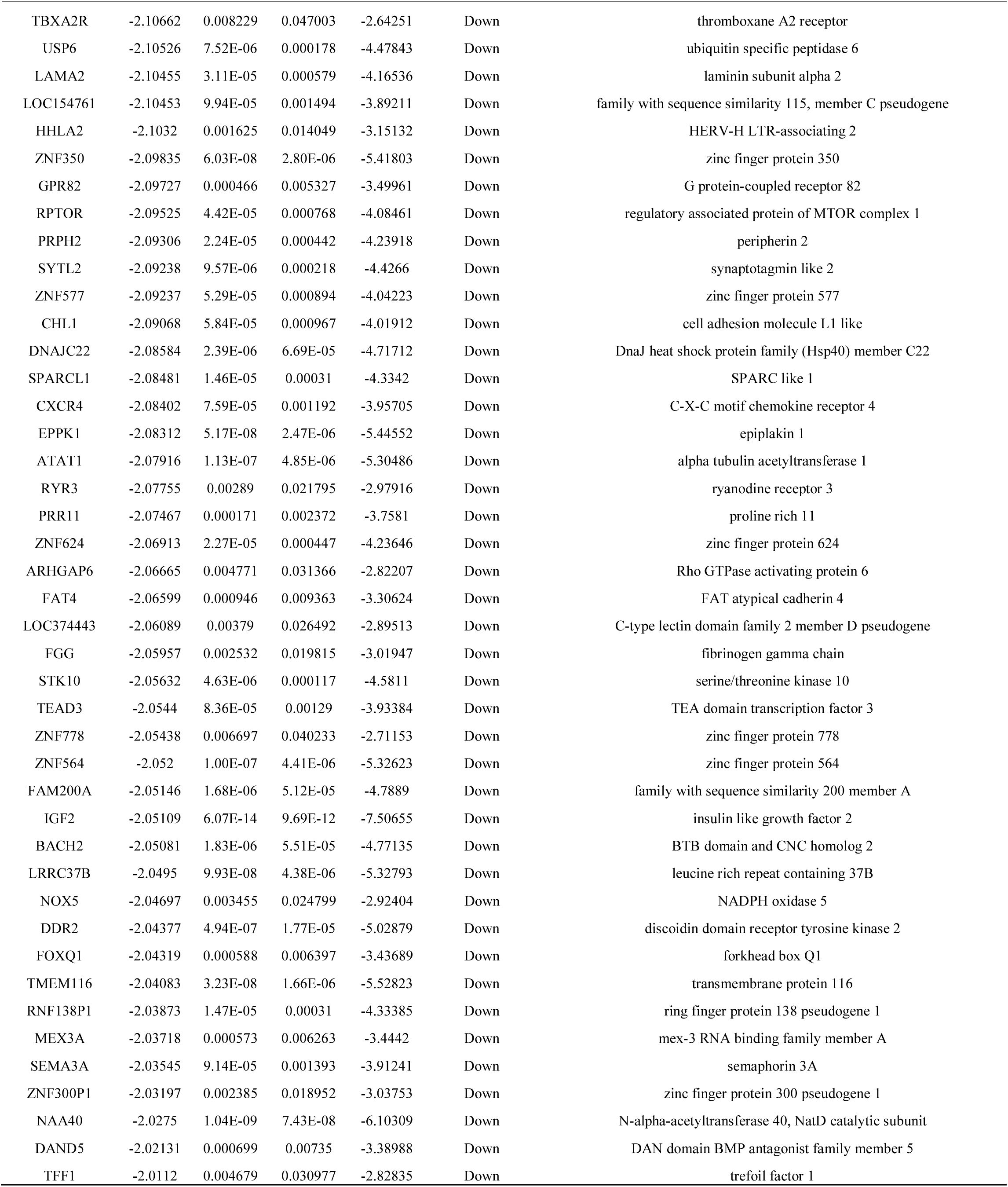
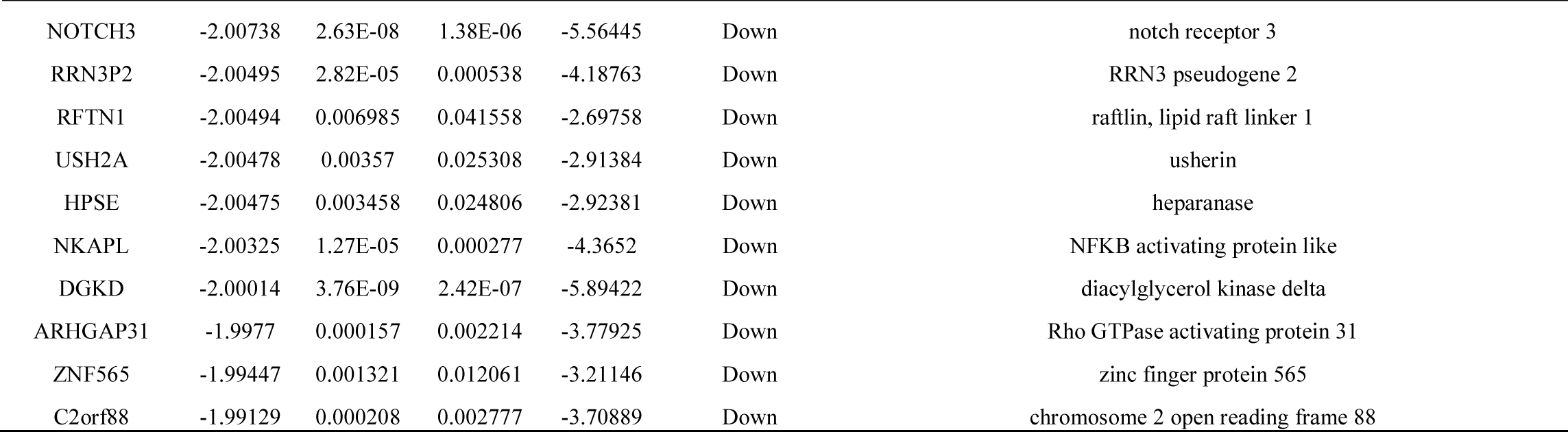
The statistical metrics for key differentially expressed genes (DEGs)

### Gene Ontology (GO) and REACTOME pathway enrichment analysis

GO enrichment analysis showed that the DEGs were mainly involved in defense response, secretion, secretory vesicle, golgi apparatus, identical protein binding, transmembrane transporter activity, cell adhesion, localization of cell, extracellular matrix, intrinsic component of plasma membrane, structural molecule activity and signaling receptor binding, and are listed in Table 3. REACTOME pathway enrichment analysis revealed that the DEGs were mainly enriched in the neutrophil degranulation, innate immune system, extracellular matrix organization and diseases of glycosylation, and are listed in Table 4.

**Table 3.**
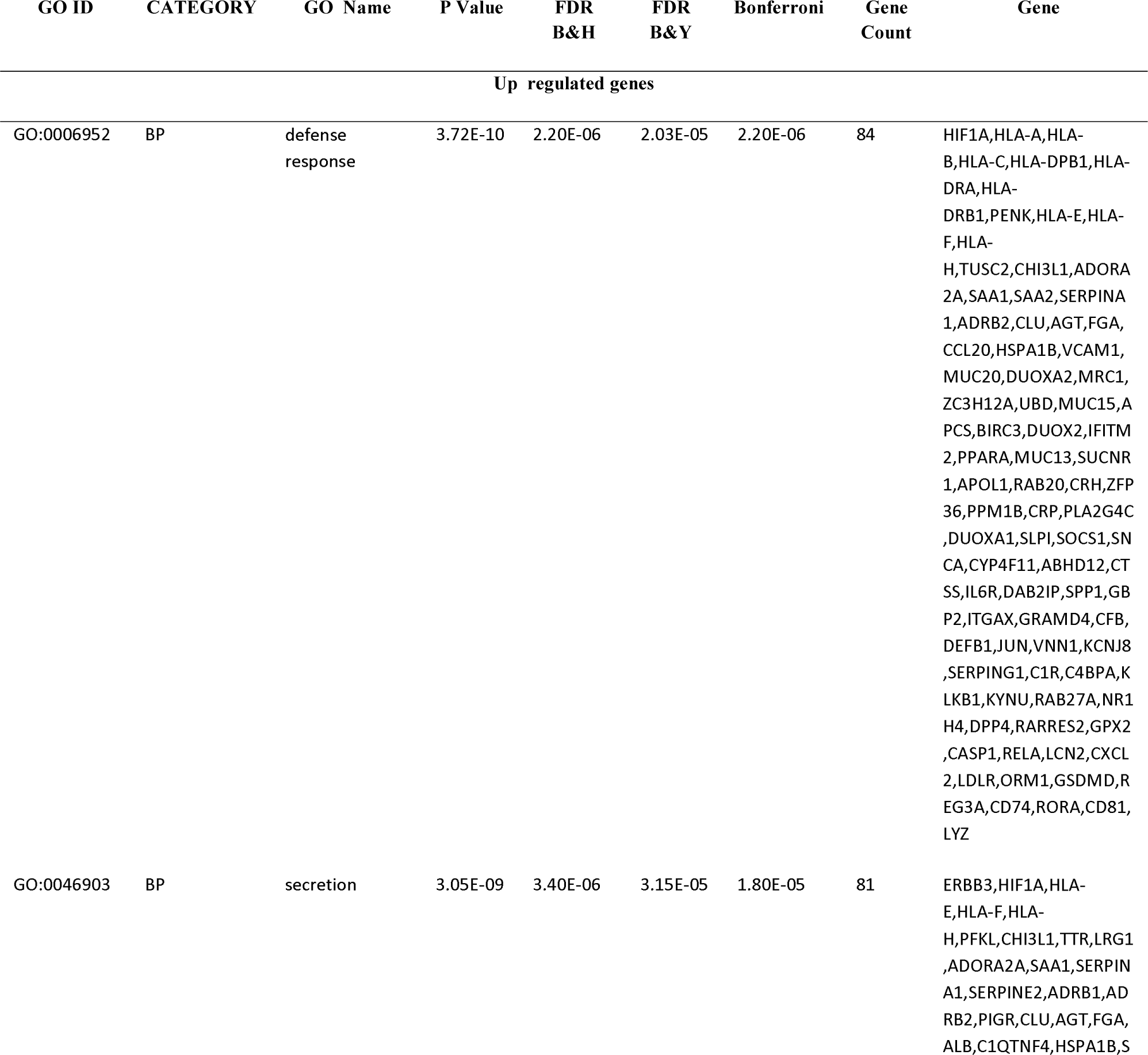
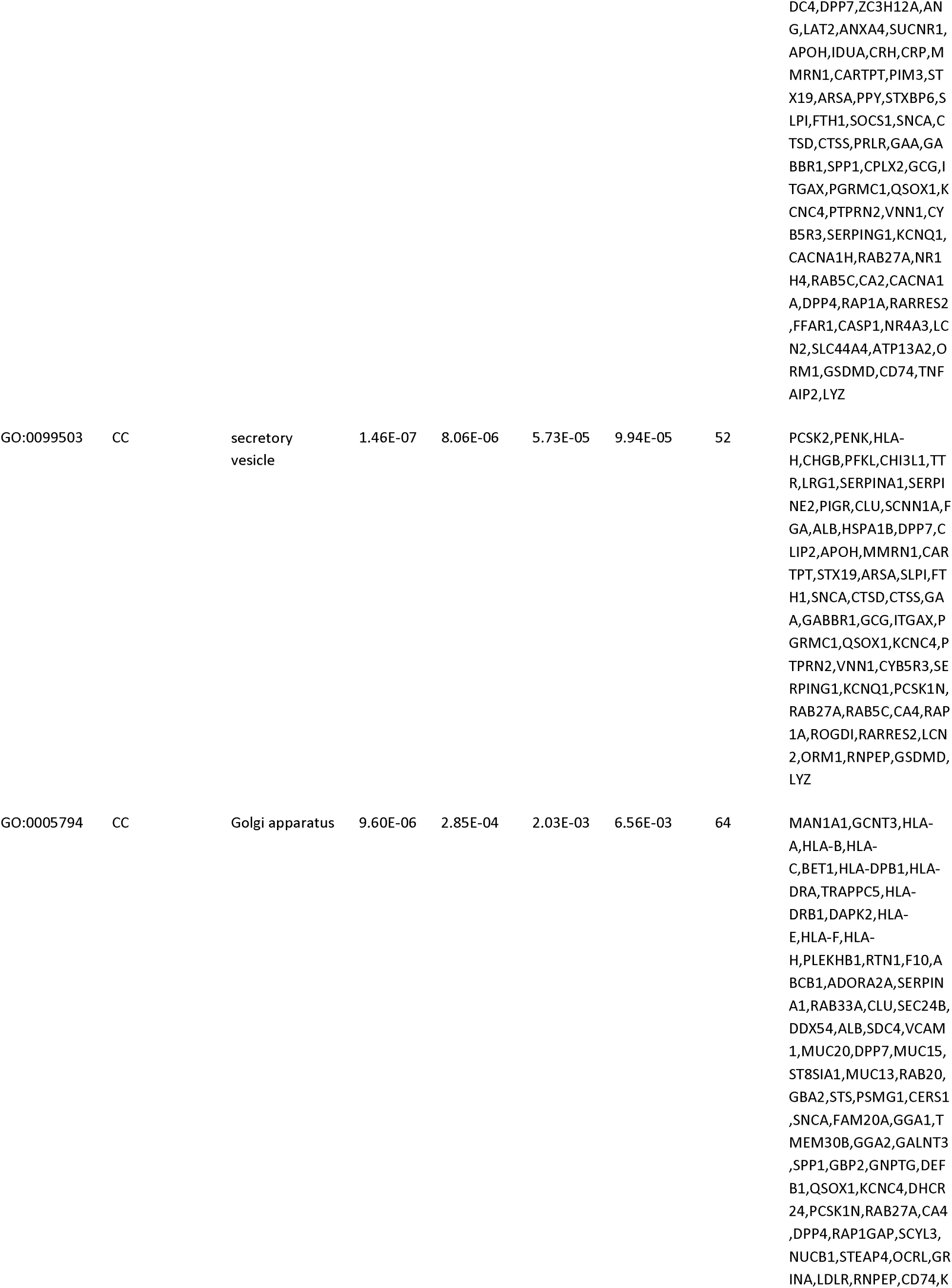
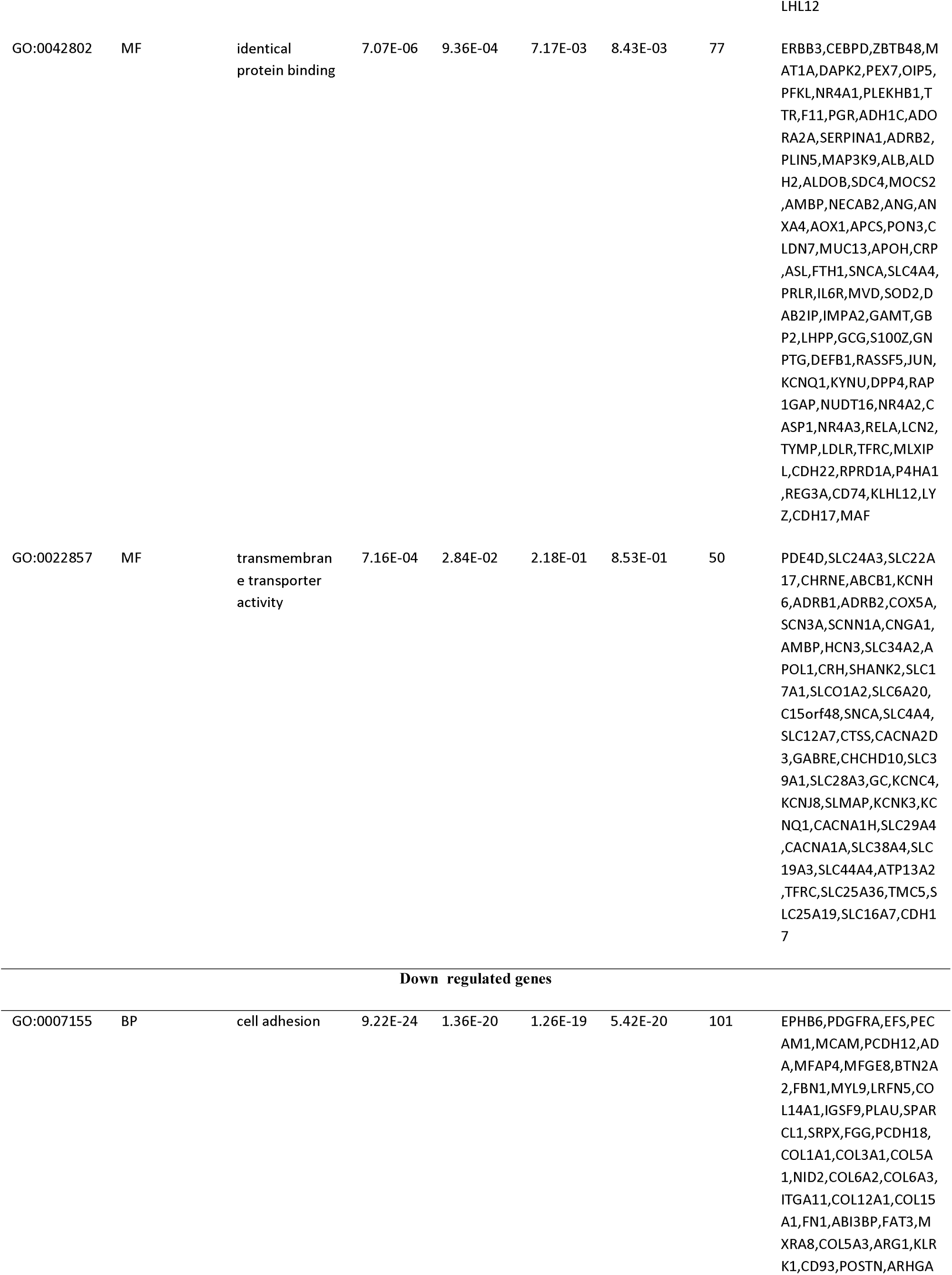
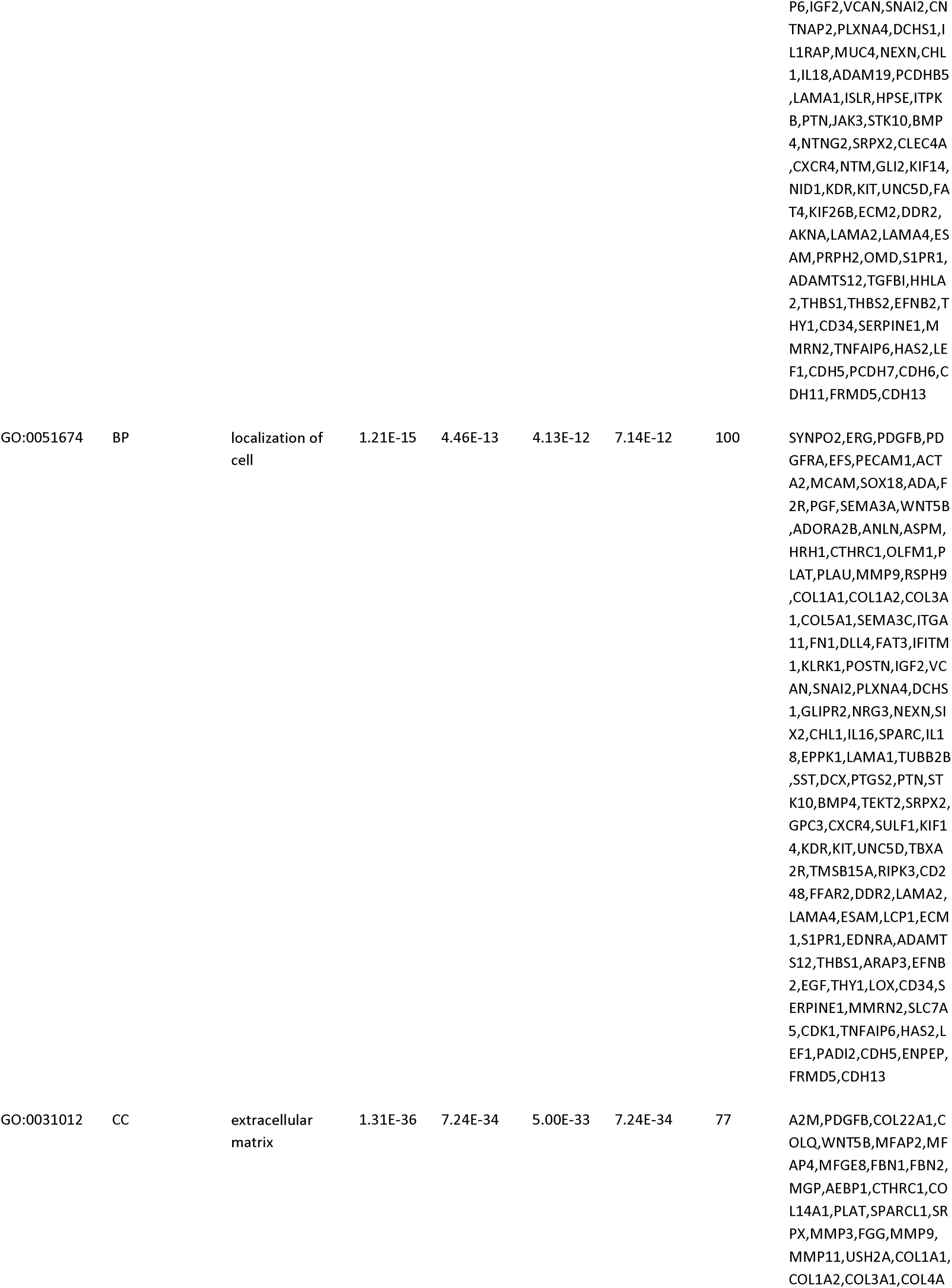
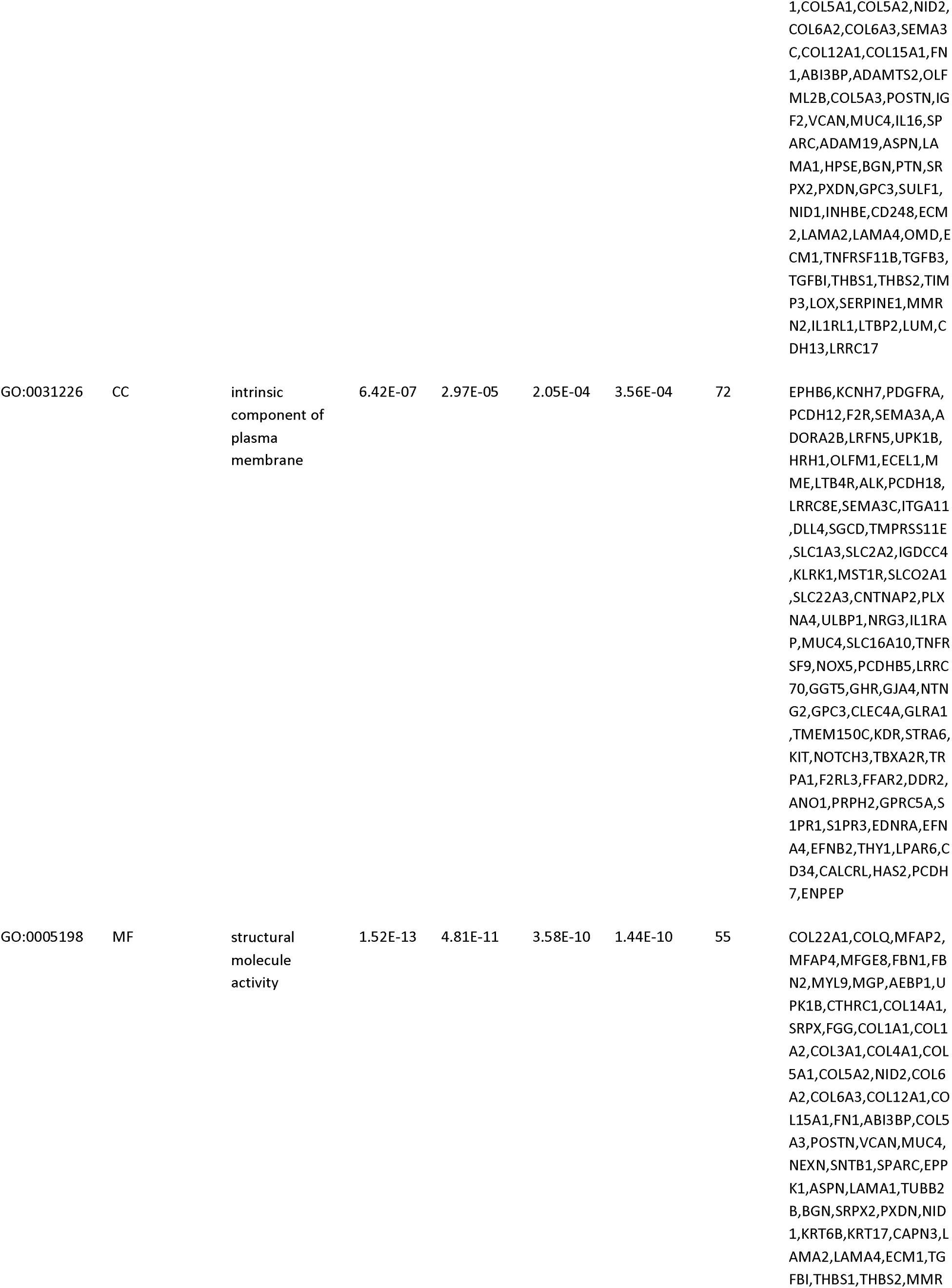
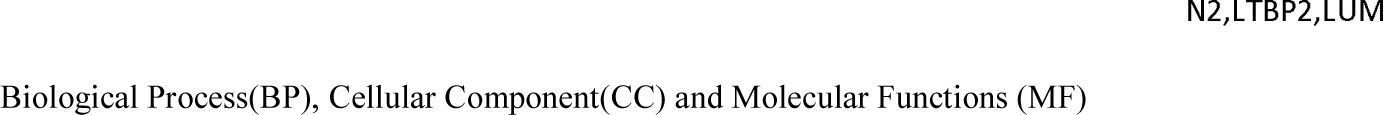
The enriched GO terms of the up and down regulated differentially expressed genes

**Table 4.**
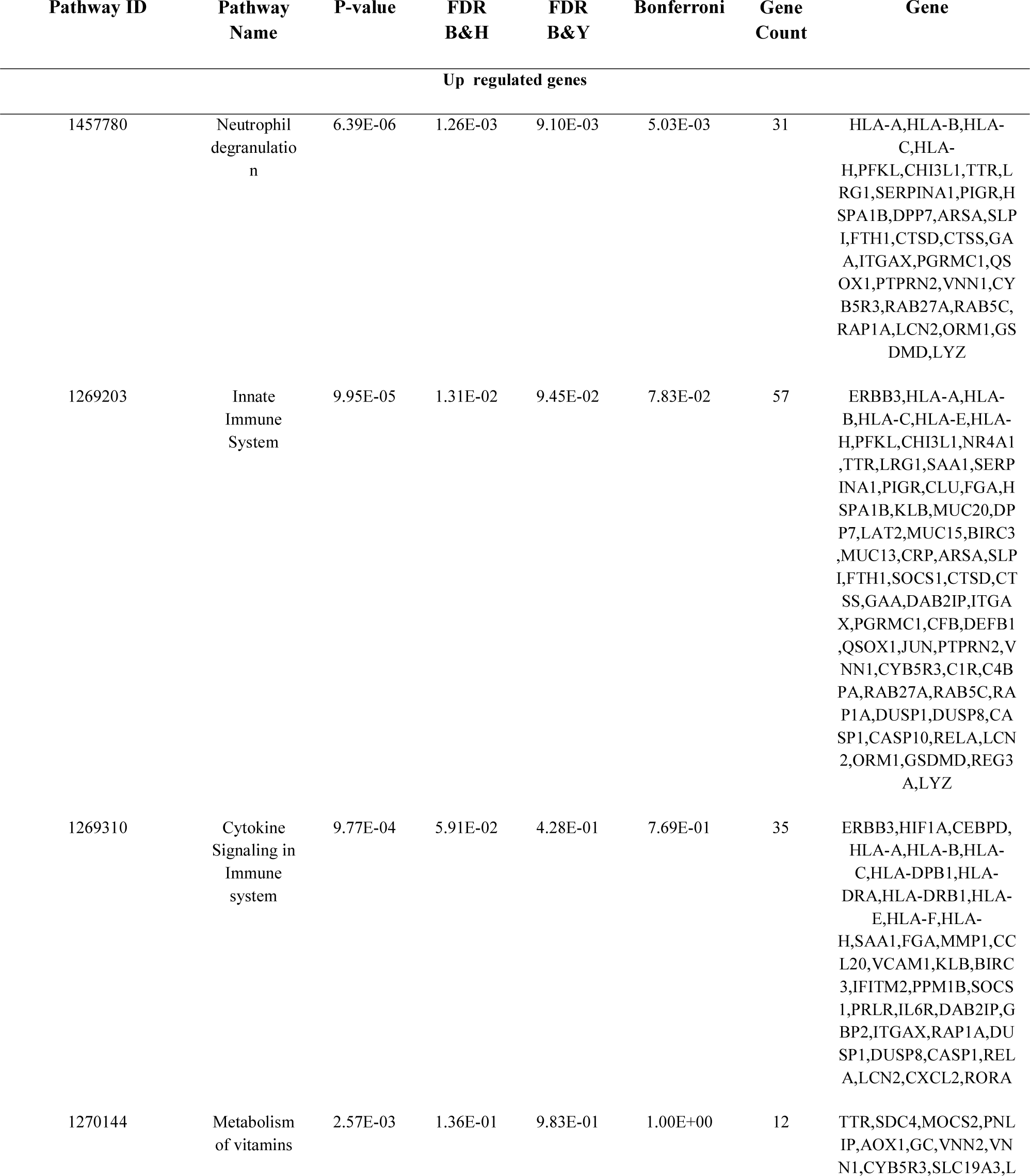
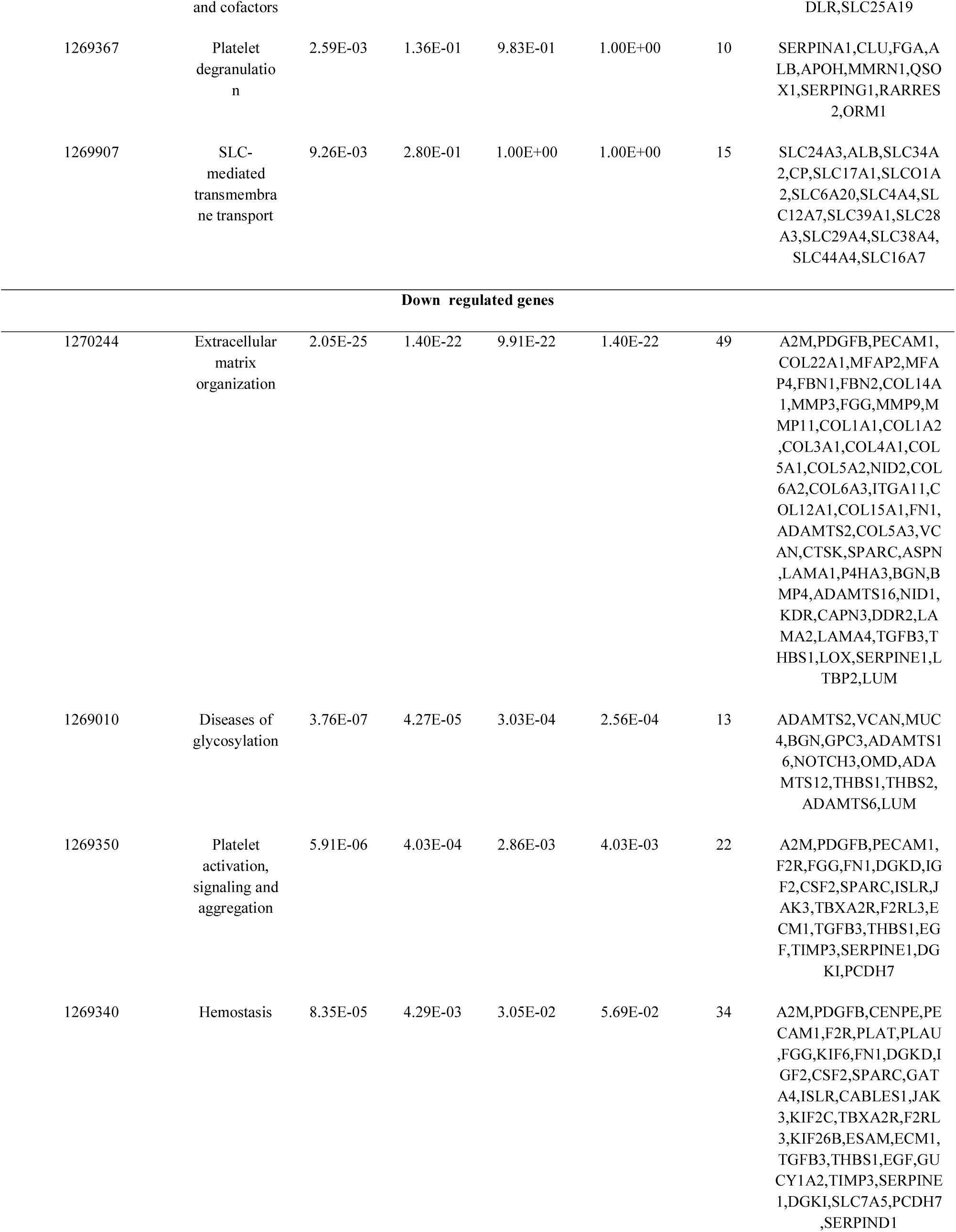
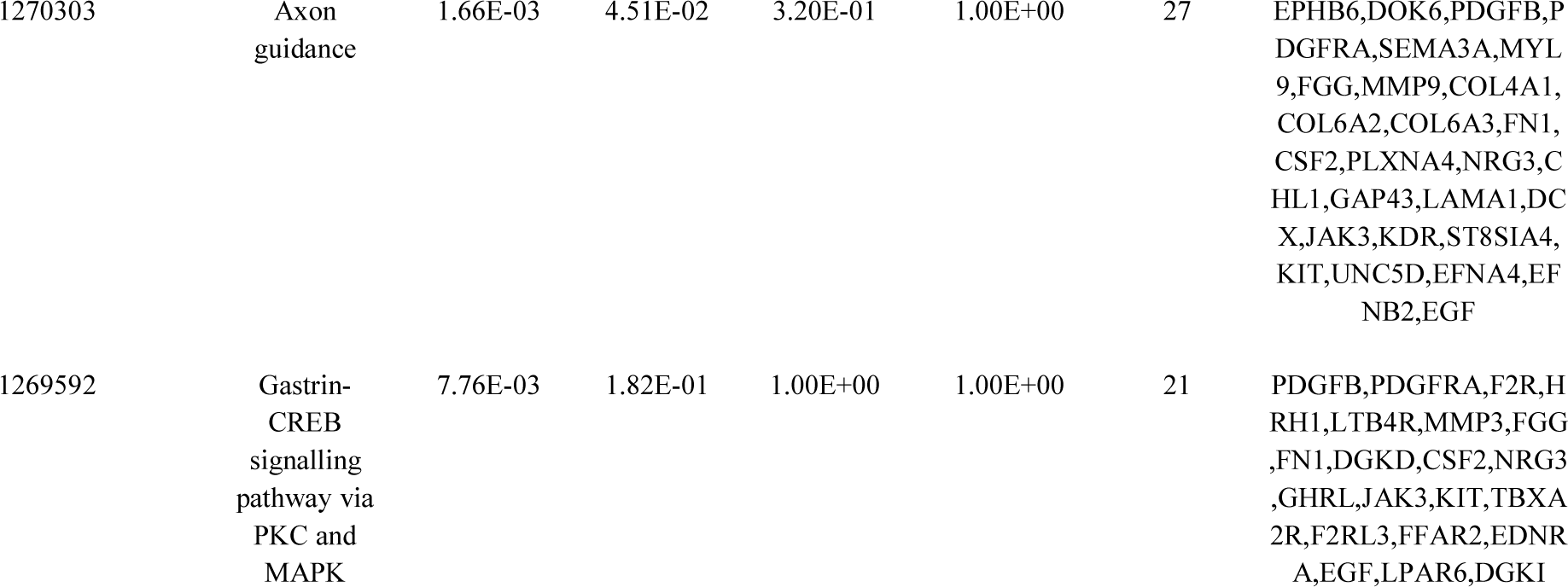
The enriched pathway terms of the up and down regulated differentially expressed genes

### PPI network construction and module analysis

The PPI network of DEGs was constructed with 7195 nodes and 16541 edges (Fig.3). The node degree, betweenness centrality, stress centrality and closeness centrality methods were performed to calculate the top ranking hub genes. The results revealed 10 genes identified as hub genes, including JUN, VCAM1, RELA, U2AF2, ADRB2, FN1, CDK1, TK1, A2M and ACTA2, and are listed Table 5. PEWCC1 was used to identify the two significant cluster modules in the PPI network and the top modules were selected (Fig.4A and Fig.4B). Gene Ontology (GO) and REACTOME pathway enrichment analysis revealed that the module 1 (48 nodes and 104 edges) and module 2 (30 nodes and 92 edges) were mainly enriched in cytokine signaling in immune system, identical protein binding, neutrophil degranulation, golgi apparatus, innate immune system, extracellular matrix organization, axon guidance, cell adhesion, platelet activation, signaling and aggregation, diseases of glycosylation, hemostasis, extracellular matrix, localization of cell and structural molecule activity.

**Fig. 3.**
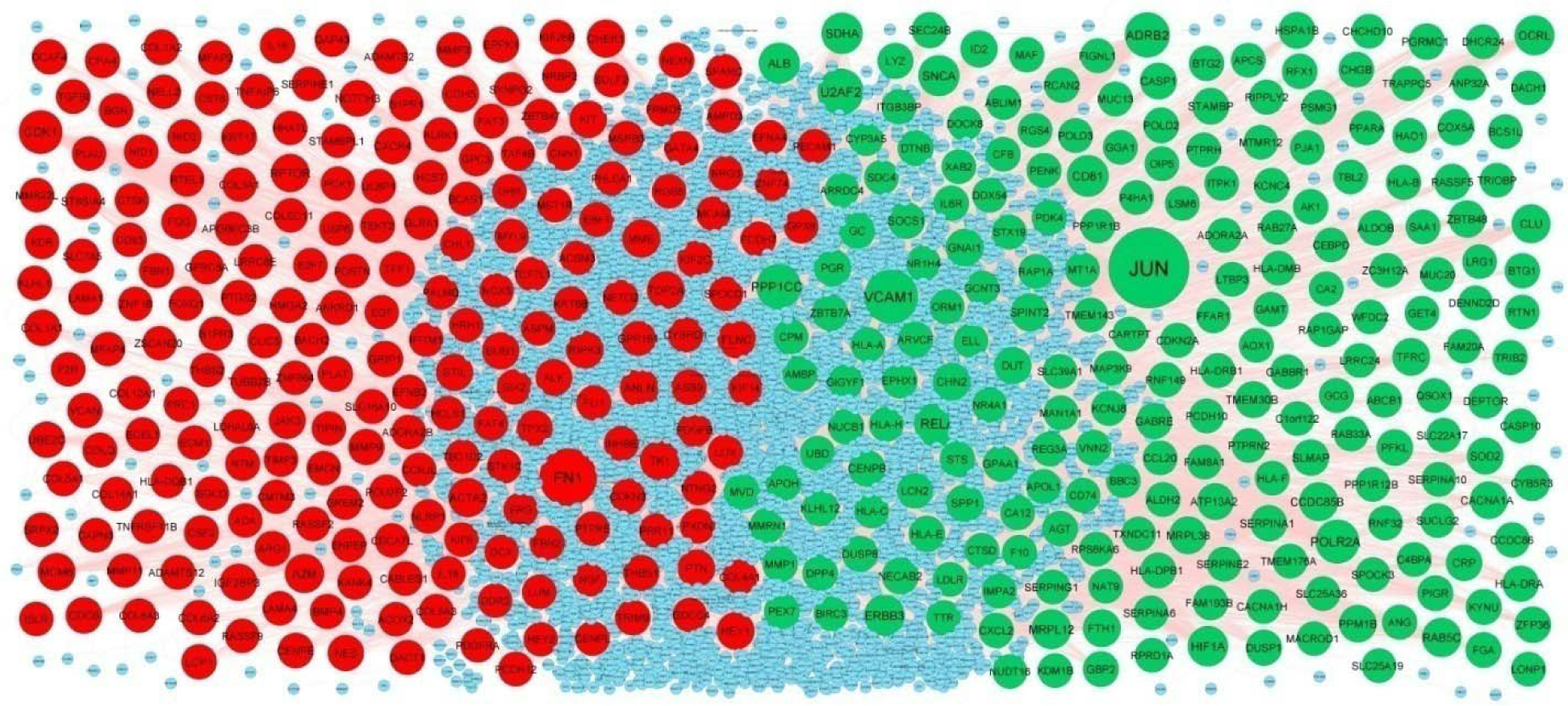
PPI network of DEGs. The PPI network of DEGs was constructed using Cytoscap. Up regulated genes are marked in green; down regulated genes are marked in red

**Fig. 4.**
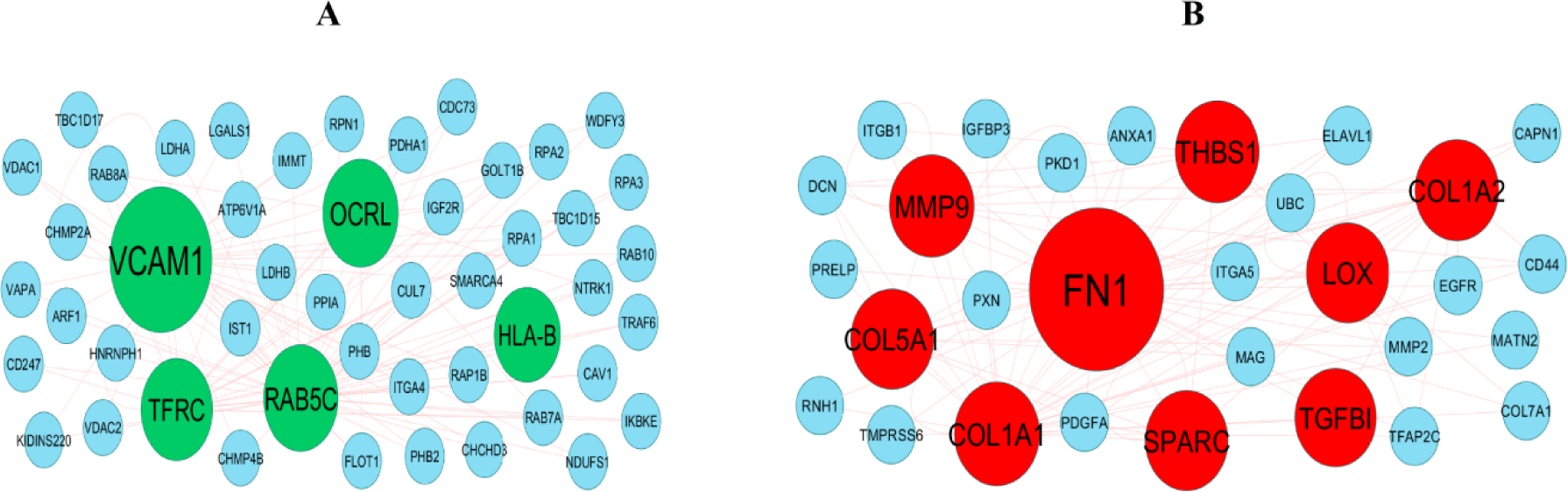
Modules of isolated form PPI of DEGs. (A) The most significant module was obtained from PPI network with 48 nodes and 104 edges for up regulated genes (B) The most significant module was obtained from PPI network with 30 nodes and 92 edges for down regulated genes. Up regulated genes are marked in green; down regulated genes are marked in red.

**Table 5.** Topology table for up and down regulated genes

| Regulation | Node | Degree | Betweenness | Stress | Closeness |
| --- | --- | --- | --- | --- | --- |
| Up | JUN | 1596 | 0.297948 | 5.04E+08 | 0.437877 |
| Up | VCAM1 | 661 | 0.055649 | 1.83E+08 | 0.379038 |
| Up | RELA | 440 | 0.066286 | 65857496 | 0.392096 |
| Up | U2AF2 | 401 | 0.032981 | 50615222 | 0.375241 |
| Up | ADRB2 | 360 | 0.051656 | 1.09E+08 | 0.357594 |
| Up | PPP1CC | 350 | 0.055551 | 48521756 | 0.382627 |
| Up | POLR2A | 320 | 0.042098 | 73677082 | 0.357008 |
| Up | CD81 | 244 | 0.02104 | 53426396 | 0.344262 |
| Up | SDHA | 228 | 0.023854 | 17009994 | 0.380039 |
| Up | SNCA | 224 | 0.025545 | 56250412 | 0.341272 |
| Up | ALB | 217 | 0.032126 | 27109520 | 0.350929 |
| Up | ERBB3 | 207 | 0.023464 | 46497836 | 0.338957 |
| Up | HIF1A | 201 | 0.022327 | 16543606 | 0.372868 |
| Up | OCRL | 183 | 0.021433 | 38355000 | 0.33816 |
| Up | MRPL12 | 182 | 0.021656 | 24757490 | 0.335917 |
| Up | RAB5C | 164 | 0.019293 | 18077018 | 0.349243 |
| Up | CCDC85B | 148 | 0.026064 | 17688698 | 0.320629 |
| Up | CLU | 148 | 0.018422 | 34351976 | 0.331337 |
| Up | FTH1 | 112 | 0.015754 | 15114778 | 0.329531 |
| Up | TFRC | 110 | 0.016321 | 10628564 | 0.378798 |
| Up | SPINT2 | 108 | 0.018333 | 13787598 | 0.314599 |
| Up | MRPL38 | 108 | 0.014795 | 7135306 | 0.347488 |
| Up | CACNA1A | 101 | 0.013778 | 14792904 | 0.318909 |
| Up | SPP1 | 100 | 0.011753 | 13635028 | 0.324872 |
| Up | GNAI1 | 100 | 0.016452 | 7052876 | 0.346084 |
| Up | PPM1B | 99 | 0.010628 | 13845334 | 0.333148 |
| Up | CHN2 | 94 | 0.003043 | 7480904 | 0.324097 |
| Up | ZBTB48 | 91 | 0.007794 | 14008360 | 0.306699 |
| Up | ID2 | 88 | 0.008382 | 20399246 | 0.31338 |
| Up | DUT | 85 | 0.006791 | 9371824 | 0.325667 |
| Up | PGR | 83 | 0.005038 | 6050756 | 0.33088 |
| Up | TMEM30B | 80 | 0.012048 | 8252762 | 0.301821 |
| Up | HSPA1B | 78 | 0.00697 | 6310746 | 0.350331 |
| Up | SOD2 | 77 | 0.006883 | 13105960 | 0.314833 |
| Up | CASP1 | 75 | 0.008457 | 14148852 | 0.309164 |
| Up | STAMBP | 73 | 0.008256 | 8962166 | 0.315013 |
| Up | NECAB2 | 72 | 0.00898 | 20828314 | 0.294 |
| Up | SOCS1 | 71 | 0.00732 | 4459136 | 0.332855 |
| Up | ATP13A2 | 70 | 0.01148 | 8100584 | 0.311952 |
| Up | CENPB | 69 | 0.00962 | 5877652 | 0.328913 |
| Up | COX5A | 68 | 0.004787 | 9014728 | 0.310793 |
| Up | PGRMC1 | 67 | 0.005965 | 5851738 | 0.324374 |
| Up | PFKL | 67 | 0.005301 | 4220404 | 0.339501 |
| Up | ZFP36 | 65 | 0.005869 | 5840334 | 0.325814 |
| Up | SLMAP | 64 | 0.008847 | 14906016 | 0.300925 |
| Up | KLHL12 | 62 | 0.006002 | 9477918 | 0.31082 |
| Up | SERPINA1 | 61 | 0.004959 | 3442108 | 0.32758 |
| Up | CRP | 61 | 0.00901 | 3976752 | 0.320944 |
| Up | HLA-DRA | 58 | 0.006376 | 6984242 | 0.303374 |
| Up | SEC24B | 57 | 0.005122 | 7921100 | 0.311736 |
| Up | HLA-E | 57 | 0.008193 | 5417646 | 0.301114 |
| Up | RASSF5 | 54 | 0.005973 | 5581726 | 0.309603 |
| Up | CYB5R3 | 53 | 0.005535 | 4457008 | 0.322007 |
| Up | DUSP1 | 50 | 0.0047 | 10162126 | 0.297576 |
| Up | LYZ | 48 | 0.004406 | 2822858 | 0.352771 |
| Up | DTNB | 46 | 0.006085 | 5998870 | 0.297047 |
| Up | GBP2 | 46 | 0.002674 | 2254472 | 0.315954 |
| Up | KYNU | 45 | 0.003541 | 8318140 | 0.294385 |
| Up | ZBTB7A | 44 | 0.003479 | 2378948 | 0.318641 |
| Up | NUCB1 | 44 | 0.003535 | 3000438 | 0.311156 |
| Up | GABRE | 42 | 0.006362 | 6963592 | 0.273467 |
| Up | DUSP8 | 42 | 0.003924 | 3730912 | 0.305695 |
| Up | FGA | 41 | 0.002177 | 1408648 | 0.322498 |
| Up | VNN2 | 39 | 0.004648 | 4918216 | 0.275309 |
| Up | PIGR | 38 | 0.003219 | 4218970 | 0.283356 |
| Up | CHGB | 37 | 0.002356 | 1883312 | 0.310458 |
| Up | HLA-DPB1 | 37 | 0.005437 | 4024006 | 0.28368 |
| Up | MVD | 37 | 0.00277 | 1785902 | 0.307209 |
| Up | GPAA1 | 36 | 0.006228 | 5917528 | 0.291675 |
| Up | POLD2 | 36 | 0.004619 | 5556090 | 0.301063 |
| Up | DPP4 | 34 | 0.004172 | 1947850 | 0.298874 |
| Up | RFX1 | 32 | 0.003758 | 7609514 | 0.295413 |
| Up | EPHX1 | 31 | 0.00324 | 3148816 | 0.29602 |
| Up | ALDOB | 31 | 0.002792 | 2018518 | 0.30784 |
| Up | AGT | 31 | 0.002966 | 1865644 | 0.306294 |
| Up | PEX7 | 28 | 0.002627 | 7979132 | 0.279546 |
| Up | MMP1 | 28 | 0.001636 | 962486 | 0.318768 |
| Up | IL6R | 28 | 0.004718 | 1522490 | 0.295316 |
| Up | AK1 | 27 | 0.002177 | 3763326 | 0.290274 |
| Up | KCNC4 | 27 | 0.002966 | 4065280 | 0.287685 |
| Up | MAF | 26 | 0.001371 | 928624 | 0.315427 |
| Up | LCN2 | 26 | 0.002077 | 4747150 | 0.281902 |
| Up | TRIB2 | 26 | 0.002374 | 1372656 | 0.307157 |
| Up | RNF149 | 26 | 0.003073 | 2741500 | 0.27934 |
| Up | GC | 25 | 0.001091 | 1093748 | 0.300309 |
| Up | BTG1 | 25 | 0.0019 | 963086 | 0.321863 |
| Up | KCNJ8 | 24 | 0.002693 | 1543874 | 0.254755 |
| Up | STS | 23 | 0.002783 | 1476396 | 0.279026 |
| Up | CCDC86 | 23 | 7.45E-04 | 1261094 | 0.295801 |
| Up | UBD | 22 | 0.001864 | 3627404 | 0.290415 |
| Up | APOH | 22 | 9.68E-04 | 1240028 | 0.29797 |
| Up | ABCB1 | 22 | 0.001386 | 2114132 | 0.29802 |
| Up | RCAN2 | 22 | 0.002346 | 3980546 | 0.273 |
| Up | DEPTOR | 22 | 0.001221 | 937788 | 0.307538 |
| Up | BTG2 | 20 | 6.52E-04 | 1046918 | 0.283703 |
| Up | ARVCF | 19 | 0.002101 | 909702 | 0.281471 |
| Up | AMBP | 17 | 7.00E-04 | 525586 | 0.31932 |
| Up | CXCL2 | 17 | 9.45E-04 | 498648 | 0.305228 |
| Up | MMRN1 | 17 | 4.60E-04 | 617814 | 0.267011 |
| Up | FAM193B | 17 | 0.001148 | 1447216 | 0.291699 |
| Up | HLA-B | 16 | 0.001753 | 1709356 | 0.316705 |
| Up | GCG | 16 | 0.001734 | 1201492 | 0.270831 |
| Up | ANG | 15 | 0.001061 | 1182188 | 0.286699 |
| Up | RIPPLY2 | 13 | 6.72E-04 | 1051108 | 0.249022 |
| Up | CDKN2A | 11 | 7.57E-04 | 1062712 | 0.32113 |
| Up | PDK4 | 11 | 5.73E-04 | 839962 | 0.257952 |
| Up | PENK | 10 | 0.001694 | 1717524 | 0.24922 |
| Up | MAN1A1 | 10 | 3.87E-04 | 238680 | 0.259994 |
| Up | GAMT | 9 | 3.76E-04 | 408938 | 0.279872 |
| Up | CYP3A5 | 8 | 0.001126 | 1514978 | 0.241538 |
| Up | AOX1 | 8 | 2.93E-04 | 399780 | 0.258536 |
| Up | HLA-DRB1 | 7 | 1.24E-04 | 74760 | 0.250776 |
| Up | HLA-A | 7 | 7.84E-04 | 861244 | 0.28934 |
| Up | RAB33A | 7 | 5.82E-04 | 1629132 | 0.221958 |
| Up | LONP1 | 6 | 5.55E-04 | 2034278 | 0.328373 |
| Up | HLA-H | 6 | 1.96E-04 | 63810 | 0.271086 |
| Up | MAP3K9 | 6 | 2.80E-04 | 523732 | 0.268857 |
| Up | ALDH2 | 5 | 2.38E-04 | 448744 | 0.310928 |
| Up | HLA-C | 5 | 4.48E-04 | 299492 | 0.271342 |
| Up | NR4A1 | 5 | 1.74E-04 | 265578 | 0.298378 |
| Up | CARTPT | 5 | 3.33E-04 | 343864 | 0.26055 |
| Up | APOL1 | 4 | 1.94E-05 | 34026 | 0.25131 |
| Up | TBL2 | 4 | 7.27E-04 | 1174624 | 0.311925 |
| Up | TTR | 4 | 5.60E-04 | 453308 | 0.270913 |
| Up | CD74 | 4 | 2.05E-04 | 127222 | 0.259189 |
| Up | CPM | 4 | 5.33E-04 | 590954 | 0.28459 |
| Up | TMEM176A | 4 | 2.82E-04 | 303148 | 0.244104 |
| Up | HAO1 | 4 | 3.12E-04 | 1039252 | 0.250113 |
| Up | CHCHD10 | 3 | 4.87E-05 | 78354 | 0.27908 |
| Up | SUCLG2 | 3 | 8.47E-05 | 93604 | 0.283569 |
| Up | CTSD | 3 | 4.02E-05 | 270406 | 0.307314 |
| Up | DDX54 | 3 | 6.87E-05 | 270908 | 0.307341 |
| Up | ANP32A | 3 | 2.05E-05 | 65462 | 0.310753 |
| Up | BIRC3 | 3 | 5.17E-05 | 155646 | 0.286608 |
| Up | GIGYF1 | 2 | 5.44E-06 | 14190 | 0.272638 |
| Up | RTN1 | 2 | 1.68E-05 | 23170 | 0.266921 |
| Up | SERPING1 | 2 | 2.26E-05 | 9370 | 0.261051 |
| Up | SERPINE2 | 2 | 2.62E-06 | 672 | 0.246784 |
| Up | TRAPPC5 | 2 | 6.26E-05 | 68134 | 0.249861 |
| Up | APCS | 2 | 0 | 0 | 0.295158 |
| Up | SERPINA6 | 2 | 3.33E-04 | 309232 | 0.296276 |
| Up | REG3A | 2 | 1.52E-05 | 52952 | 0.299908 |
| Up | BBC3 | 2 | 0 | 0 | 0.259778 |
| Up | RPRD1A | 2 | 3.92E-05 | 277120 | 0.309669 |
| Up | DACH1 | 2 | 0 | 0 | 0.309936 |
| Up | RGS4 | 2 | 5.44E-05 | 21664 | 0.272307 |
| Up | CASP10 | 2 | 3.66E-05 | 73078 | 0.264702 |
| Up | PPARA | 2 | 3.08E-05 | 28360 | 0.285403 |
| Up | DHCR24 | 2 | 2.06E-05 | 30004 | 0.275109 |
| Up | RAP1A | 2 | 3.50E-05 | 22882 | 0.262835 |
| Up | PCDH10 | 2 | 6.75E-06 | 11790 | 0.233911 |
| Up | F10 | 2 | 2.24E-06 | 2690 | 0.247787 |
| Up | PSMG1 | 2 | 0 | 0 | 0.29646 |
| Up | CFB | 2 | 0 | 0 | 0.297072 |
| Up | PPP1R12B | 2 | 1.19E-05 | 11496 | 0.250122 |
| Up | ABLIM1 | 2 | 4.21E-05 | 35910 | 0.250479 |
| Up | LTBP3 | 1 | 0 | 0 | 0.241806 |
| Up | NAT9 | 1 | 0 | 0 | 0.267189 |
| Up | GGA1 | 1 | 0 | 0 | 0.229025 |
| Up | PJA1 | 1 | 0 | 0 | 0.239559 |
| Up | SERPINA10 | 1 | 0 | 0 | 0.240834 |
| Up | RNF32 | 1 | 0 | 0 | 0.249376 |
| Up | HLA-DMB | 1 | 0 | 0 | 0.232768 |
| Up | SAA1 | 1 | 0 | 0 | 0.241757 |
| Up | C4BPA | 1 | 0 | 0 | 0.294264 |
| Up | GABBR1 | 1 | 0 | 0 | 0.259778 |
| Up | GET4 | 1 | 0 | 0 | 0.239455 |
| Up | MUC20 | 1 | 0 | 0 | 0.235388 |
| Up | NUDT16 | 1 | 0 | 0 | 0.304543 |
| Up | ITPK1 | 1 | 0 | 0 | 0.304543 |
| Up | FIGNL1 | 1 | 0 | 0 | 0.304543 |
| Up | SLC25A19 | 1 | 0 | 0 | 0.304543 |
| Up | RAP1GAP | 1 | 0 | 0 | 0.282411 |
| Up | RPS6KA6 | 1 | 0 | 0 | 0.247514 |
| Up | ARRDC4 | 1 | 0 | 0 | 0.263412 |
| Up | RAB27A | 1 | 0 | 0 | 0.263412 |
| Up | SPOCK3 | 1 | 0 | 0 | 0.236807 |
| Up | DENND2D | 1 | 0 | 0 | 0.25945 |
| Up | FAM20A | 1 | 0 | 0 | 0.245219 |
| Up | TXNDC11 | 1 | 0 | 0 | 0.248884 |
| Up | CCL20 | 1 | 0 | 0 | 0.219681 |
| Up | HLA-F | 1 | 0 | 0 | 0.231435 |
| Up | LRRC24 | 1 | 0 | 0 | 0.231435 |
| Up | BCS1L | 1 | 0 | 0 | 0.274867 |
| Up | MACROD1 | 1 | 0 | 0 | 0.274867 |
| Up | KDM1B | 1 | 0 | 0 | 0.237111 |
| Up | WFDC2 | 1 | 0 | 0 | 0.24227 |
| Up | SLC25A36 | 1 | 0 | 0 | 0.24227 |
| Up | PTPRH | 1 | 0 | 0 | 0.253159 |
| Up | ELL | 1 | 0 | 0 | 0.263094 |
| Up | XAB2 | 1 | 0 | 0 | 0.263094 |
| Up | MTMR12 | 1 | 0 | 0 | 0.229339 |
| Up | PPP1R1B | 1 | 0 | 0 | 0.27675 |
| Up | POLD3 | 1 | 0 | 0 | 0.231405 |
| Up | LSM6 | 1 | 0 | 0 | 0.231405 |
| Up | MUC13 | 1 | 0 | 0 | 0.258853 |
| Up | C1orf122 | 1 | 0 | 0 | 0.25146 |
| Up | SLC22A17 | 1 | 0 | 0 | 0.219916 |
| Up | NR1H4 | 1 | 0 | 0 | 0.28167 |
| Up | ITGB3BP | 1 | 0 | 0 | 0.28167 |
| Up | PTPRN2 | 1 | 0 | 0 | 0.245663 |
| Up | ZC3H12A | 1 | 0 | 0 | 0.271608 |
| Up | TMEM143 | 1 | 0 | 0 | 0.231853 |
| Up | FAM8A1 | 1 | 0 | 0 | 0.237785 |
| Up | QSOX1 | 1 | 0 | 0 | 0.242974 |
| Up | LDLR | 1 | 0 | 0 | 0.229574 |
| Up | LRG1 | 1 | 0 | 0 | 0.294264 |
| Up | ORM1 | 1 | 0 | 0 | 0.259778 |
| Up | CA2 | 1 | 0 | 0 | 0.304543 |
| Up | OIP5 | 1 | 0 | 0 | 0.304543 |
| Up | IMPA2 | 1 | 0 | 0 | 0.245671 |
| Up | SDC4 | 1 | 0 | 0 | 0.252041 |
| Up | GCNT3 | 1 | 0 | 0 | 0.252041 |
| Up | MT1A | 1 | 0 | 0 | 0.257113 |
| Up | FFAR1 | 1 | 0 | 0 | 0.257113 |
| Up | CEBPD | 1 | 0 | 0 | 0.28167 |
| Up | CA12 | 1 | 0 | 0 | 0.231853 |
| Up | ADORA2A | 1 | 0 | 0 | 0.22721 |
| Up | DOCK8 | 1 | 0 | 0 | 0.257888 |
| Up | STX19 | 1 | 0 | 0 | 0.230169 |
| Up | TRIOBP | 1 | 0 | 0 | 0.242237 |
| Up | CACNA1H | 1 | 0 | 0 | 0.220929 |
| Up | SLC39A1 | 1 | 0 | 0 | 0.239591 |
| Up | P4HA1 | 1 | 0 | 0 | 0.234338 |
| Down | FN1 | 800 | 0.121948 | 1.77E+08 | 0.416937 |
| Down | CDK1 | 321 | 0.052371 | 37130306 | 0.393533 |
| Down | TK1 | 183 | 0.028359 | 22061908 | 0.356407 |
| Down | A2M | 165 | 0.021657 | 31093610 | 0.33221 |
| Down | ACTA2 | 156 | 0.021508 | 16177182 | 0.356813 |
| Down | ANLN | 147 | 0.012288 | 32652030 | 0.31966 |
| Down | RPTOR | 138 | 0.015393 | 20848032 | 0.33123 |
| Down | FLNC | 130 | 0.020359 | 9885742 | 0.364849 |
| Down | MME | 124 | 0.01613 | 22827922 | 0.327014 |
| Down | CHEK1 | 123 | 0.0138 | 9428096 | 0.36459 |
| Down | MCM6 | 119 | 0.012797 | 10240308 | 0.34408 |
| Down | HEY1 | 115 | 0.007467 | 26394370 | 0.324521 |
| Down | TOP2A | 113 | 0.00822 | 7037480 | 0.366131 |
| Down | PTN | 109 | 0.01344 | 17516900 | 0.319717 |
| Down | RIPK3 | 109 | 0.005618 | 13238646 | 0.329395 |
| Down | KIT | 105 | 0.00936 | 8415988 | 0.318274 |
| Down | BUB1 | 102 | 0.012926 | 9378750 | 0.327669 |
| Down | IGF2BP3 | 101 | 0.006201 | 5000402 | 0.358021 |
| Down | CDH5 | 97 | 0.014275 | 14697350 | 0.31753 |
| Down | STIL | 94 | 0.010914 | 12929196 | 0.315912 |
| Down | JAK3 | 93 | 0.006847 | 6645686 | 0.330743 |
| Down | TUBB2B | 89 | 0.011127 | 6631350 | 0.358521 |
| Down | CXCR4 | 82 | 0.010003 | 6983362 | 0.336956 |
| Down | ALK | 79 | 0.006945 | 9288848 | 0.319235 |
| Down | COL1A1 | 78 | 0.007539 | 4482218 | 0.336184 |
| Down | KIF2C | 78 | 0.00852 | 6501484 | 0.336909 |
| Down | NOTCH3 | 78 | 0.010163 | 11236576 | 0.315068 |
| Down | CDC6 | 74 | 0.00694 | 5538892 | 0.323892 |
| Down | UBE2C | 69 | 0.00668 | 5381164 | 0.318429 |
| Down | ST8SIA4 | 68 | 0.011163 | 8424288 | 0.30563 |
| Down | EPPK1 | 68 | 0.004854 | 7870332 | 0.320301 |
| Down | NES | 63 | 0.005965 | 5426230 | 0.324301 |
| Down | GRIP1 | 61 | 0.006138 | 14613812 | 0.304595 |
| Down | LEF1 | 60 | 0.005343 | 4927416 | 0.310793 |
| Down | MMP9 | 60 | 0.006943 | 5669948 | 0.345071 |
| Down | THBS1 | 57 | 0.003128 | 2389380 | 0.326747 |
| Down | COL1A2 | 48 | 0.003029 | 1783500 | 0.32932 |
| Down | TPX2 | 46 | 0.003701 | 1798214 | 0.340352 |
| Down | KRT17 | 45 | 0.003227 | 3063576 | 0.325652 |
| Down | CENPE | 43 | 0.003357 | 2673372 | 0.310807 |
| Down | TRIM9 | 43 | 0.004332 | 4076190 | 0.298973 |
| Down | PLAT | 42 | 0.004548 | 2384326 | 0.317222 |
| Down | NEXN | 40 | 0.002111 | 1827254 | 0.320901 |
| Down | ARG1 | 39 | 0.003041 | 4412562 | 0.305942 |
| Down | PTGS2 | 39 | 0.002083 | 2565408 | 0.316649 |
| Down | PCDH7 | 38 | 0.004648 | 3503260 | 0.300108 |
| Down | TFF1 | 38 | 0.002965 | 1245772 | 0.334139 |
| Down | FGG | 37 | 0.002101 | 1367870 | 0.309656 |
| Down | HMGA2 | 36 | 9.38E-04 | 969146 | 0.320587 |
| Down | LCP1 | 35 | 0.002865 | 1240328 | 0.343735 |
| Down | COL5A1 | 34 | 0.001354 | 878512 | 0.318627 |
| Down | MMP3 | 34 | 0.002251 | 1415208 | 0.31027 |
| Down | ASB9 | 34 | 0.001201 | 3273876 | 0.29823 |
| Down | COL6A2 | 33 | 0.003726 | 2359320 | 0.308699 |
| Down | IL16 | 33 | 0.003152 | 1947338 | 0.313625 |
| Down | PRC1 | 32 | 0.002506 | 1394808 | 0.310378 |
| Down | SPARC | 32 | 0.00126 | 838072 | 0.31898 |
| Down | NETO2 | 32 | 0.003703 | 1704326 | 0.304298 |
| Down | LOX | 32 | 0.001666 | 910394 | 0.322152 |
| Down | CDKN3 | 32 | 0.003019 | 1564204 | 0.302991 |
| Down | COL4A1 | 31 | 0.001704 | 998644 | 0.318825 |
| Down | VCAN | 31 | 0.002243 | 3325576 | 0.281515 |
| Down | TNFAIP6 | 29 | 0.004894 | 3551464 | 0.309031 |
| Down | BMP4 | 29 | 0.00477 | 6323086 | 0.261868 |
| Down | DCAF4 | 29 | 0.002018 | 6457740 | 0.29138 |
| Down | DACT1 | 28 | 0.001674 | 1236348 | 0.321216 |
| Down | FBN2 | 27 | 0.002317 | 1481936 | 0.317068 |
| Down | FBN1 | 27 | 0.001379 | 1080758 | 0.307209 |
| Down | CPA4 | 27 | 0.002556 | 996084 | 0.289201 |
| Down | PXDN | 26 | 0.002559 | 2031162 | 0.300221 |
| Down | CNN1 | 24 | 0.002226 | 1637732 | 0.325932 |
| Down | GAP43 | 24 | 0.00294 | 1612258 | 0.325343 |
| Down | FAT3 | 23 | 0.002598 | 2919422 | 0.275457 |
| Down | KIF26B | 22 | 0.001863 | 792688 | 0.306372 |
| Down | CYBRD1 | 22 | 0.001123 | 1561118 | 0.291533 |
| Down | LRRC8E | 22 | 0.001826 | 2528860 | 0.287639 |
| Down | SIX2 | 22 | 0.001973 | 2753556 | 0.291415 |
| Down | POU2F2 | 21 | 0.001426 | 1334750 | 0.295316 |
| Down | FLI1 | 21 | 0.00112 | 2415044 | 0.285052 |
| Down | GPR161 | 20 | 0.002009 | 2695240 | 0.292303 |
| Down | PCDH12 | 19 | 0.001536 | 1167904 | 0.272989 |
| Down | ADAMTS2 | 19 | 0.002709 | 2167276 | 0.259404 |
| Down | ANKRD1 | 19 | 0.001457 | 2414200 | 0.271906 |
| Down | TEKT2 | 19 | 0.002071 | 3016046 | 0.282477 |
| Down | COLQ | 18 | 0.003044 | 3000882 | 0.306046 |
| Down | MSRB3 | 18 | 0.002113 | 3276728 | 0.289003 |
| Down | CSF2 | 17 | 1.81E-04 | 232386 | 0.284073 |
| Down | INHBE | 17 | 0.001905 | 3619024 | 0.291805 |
| Down | NTM | 17 | 0.001536 | 2018548 | 0.280112 |
| Down | SULF2 | 16 | 0.001763 | 1637484 | 0.282677 |
| Down | ADAMTS12 | 14 | 3.95E-04 | 228572 | 0.287916 |
| Down | CABLES1 | 14 | 4.83E-04 | 277378 | 0.300472 |
| Down | KANK4 | 14 | 0.002325 | 3632254 | 0.267836 |
| Down | EBF1 | 14 | 0.001351 | 2657292 | 0.252404 |
| Down | TGFB1 | 13 | 4.16E-04 | 410792 | 0.302748 |
| Down | CLIC5 | 13 | 6.32E-04 | 350440 | 0.308236 |
| Down | KLRK1 | 13 | 0.001636 | 1607184 | 0.253865 |
| Down | SLC16A10 | 13 | 0.002048 | 1684164 | 0.231182 |
| Down | KLHL1 | 12 | 8.40E-04 | 685122 | 0.283569 |
| Down | ULBP1 | 11 | 0.001627 | 1909902 | 0.252616 |
| Down | CENPL | 11 | 0.001015 | 1075848 | 0.281835 |
| Down | EFNA4 | 10 | 0.001001 | 1689268 | 0.262691 |
| Down | SGCD | 10 | 0.001345 | 549950 | 0.269018 |
| Down | TNFRSF11B | 8 | 8.96E-04 | 1069060 | 0.296057 |
| Down | ERG | 8 | 9.91E-04 | 1598524 | 0.326955 |
| Down | PDGFB | 8 | 2.27E-04 | 209072 | 0.266763 |
| Down | GPX8 | 8 | 9.35E-04 | 811058 | 0.233017 |
| Down | ADORA2B | 7 | 8.76E-04 | 1026210 | 0.25586 |
| Down | POSTN | 6 | 8.80E-04 | 1154784 | 0.237699 |
| Down | KIF6 | 6 | 8.38E-04 | 1734970 | 0.202506 |
| Down | HLA-DQB1 | 5 | 3.89E-04 | 258228 | 0.235782 |
| Down | NTNG2 | 5 | 1.04E-04 | 119956 | 0.2581 |
| Down | HHATL | 4 | 3.03E-04 | 173754 | 0.247812 |
| Down | SRPX2 | 4 | 1.18E-05 | 10068 | 0.23472 |
| Down | TIMP3 | 4 | 2.01E-04 | 158044 | 0.310097 |
| Down | HCST | 4 | 2.61E-04 | 63872 | 0.273894 |
| Down | TCF7L1 | 4 | 1.72E-05 | 21986 | 0.253936 |
| Down | ZBTB47 | 4 | 2.82E-04 | 354140 | 0.266063 |
| Down | SALL4 | 4 | 1.14E-05 | 20262 | 0.267368 |
| Down | CST6 | 3 | 5.80E-05 | 89820 | 0.250776 |
| Down | LAMA1 | 3 | 6.11E-05 | 12132 | 0.257601 |
| Down | NID1 | 3 | 4.30E-06 | 4204 | 0.258499 |
| Down | BGN | 3 | 8.53E-05 | 103066 | 0.254728 |
| Down | PDGFRA | 3 | 7.58E-05 | 77934 | 0.284916 |
| Down | COL3A1 | 3 | 1.64E-06 | 1006 | 0.253096 |
| Down | ADA | 3 | 3.21E-05 | 15284 | 0.235058 |
| Down | NELL2 | 2 | 2.10E-05 | 14054 | 0.246513 |
| Down | KDR | 2 | 2.89E-05 | 27088 | 0.254746 |
| Down | ECEL1 | 2 | 4.32E-05 | 24446 | 0.234537 |
| Down | STAMBPL1 | 2 | 4.96E-05 | 27352 | 0.254268 |
| Down | SERPINE1 | 2 | 2.85E-06 | 2312 | 0.251741 |
| Down | CD93 | 2 | 0 | 0 | 0.253811 |
| Down | ZNF16 | 2 | 0 | 0 | 0.253811 |
| Down | NID2 | 2 | 1.07E-06 | 1590 | 0.25443 |
| Down | IGDCC4 | 2 | 1.01E-04 | 72314 | 0.262566 |
| Down | ECM1 | 2 | 4.51E-06 | 13198 | 0.248764 |
| Down | F2R | 2 | 2.19E-05 | 6442 | 0.256902 |
| Down | ASPM | 2 | 1.17E-04 | 27730 | 0.255107 |
| Down | THBS2 | 2 | 0 | 0 | 0.264556 |
| Down | COL5A3 | 2 | 0 | 0 | 0.251187 |
| Down | GREM2 | 2 | 1.13E-06 | 1018 | 0.207542 |
| Down | COL14A1 | 2 | 3.69E-05 | 13532 | 0.278335 |
| Down | GLRA1 | 2 | 0 | 0 | 0 |
| Down | LDHAL6A | 2 | 8.46E-06 | 18176 | 0.280834 |
| Down | MFAP2 | 2 | 0 | 0 | 0.242474 |
| Down | MST1R | 2 | 2.48E-05 | 76094 | 0.293269 |
| Down | FOXQ1 | 2 | 2.41E-06 | 5328 | 0.256728 |
| Down | PRR11 | 2 | 3.99E-06 | 4210 | 0.259441 |
| Down | PLAU | 2 | 3.89E-06 | 3598 | 0.246159 |
| Down | RGS5 | 2 | 3.15E-05 | 39976 | 0.279535 |
| Down | HCLS1 | 2 | 1.38E-05 | 64778 | 0.279764 |
| Down | RASSF9 | 2 | 0 | 0 | 0.279275 |
| Down | CAPN3 | 2 | 1.29E-04 | 71456 | 0.26935 |
| Down | EMCN | 2 | 3.95E-06 | 4814 | 0.206713 |
| Down | PTPRE | 1 | 0 | 0 | 0.244935 |
| Down | GHR | 1 | 0 | 0 | 0.24974 |
| Down | KIF14 | 1 | 0 | 0 | 0.23932 |
| Down | FRMD5 | 1 | 0 | 0 | 0.23932 |
| Down | TBC1D2 | 1 | 0 | 0 | 0.246809 |
| Down | MMP11 | 1 | 0 | 0 | 0.246758 |
| Down | ENPEP | 1 | 0 | 0 | 0.234483 |
| Down | NGF | 1 | 0 | 0 | 0.249376 |
| Down | DDR2 | 1 | 0 | 0 | 0.251609 |
| Down | AMPD3 | 1 | 0 | 0 | 0.259778 |
| Down | KRT6B | 1 | 0 | 0 | 0.259778 |
| Down | PALMD | 1 | 0 | 0 | 0.304543 |
| Down | BACH2 | 1 | 0 | 0 | 0.304543 |
| Down | NOX5 | 1 | 0 | 0 | 0.304543 |
| Down | USP6 | 1 | 0 | 0 | 0.304543 |
| Down | MCAM | 1 | 0 | 0 | 0.263412 |
| Down | LUM | 1 | 0 | 0 | 0.247744 |
| Down | CTSK | 1 | 0 | 0 | 0.241847 |
| Down | CMTM3 | 1 | 0 | 0 | 0.245219 |
| Down | FAT4 | 1 | 0 | 0 | 0.248884 |
| Down | COL12A1 | 1 | 0 | 0 | 0.256554 |
| Down | ACSM3 | 1 | 0 | 0 | 0.274867 |
| Down | LAMA4 | 1 | 0 | 0 | 0.24227 |
| Down | PHLDA1 | 1 | 0 | 0 | 0.253159 |
| Down | ZNF74 | 1 | 0 | 0 | 0.263094 |
| Down | NLRP1 | 1 | 0 | 0 | 0.236161 |
| Down | IL18 | 1 | 0 | 0 | 0.236161 |
| Down | MFAP4 | 1 | 0 | 0 | 0.235019 |
| Down | APOBEC3B | 1 | 0 | 0 | 0.27675 |
| Down | SPOCD1 | 1 | 0 | 0 | 0.27675 |
| Down | GPRC5A | 1 | 0 | 0 | 0.258853 |
| Down | E2F7 | 1 | 0 | 0 | 0.244776 |
| Down | NRG3 | 1 | 0 | 0 | 0.205779 |
| Down | S1PR3 | 1 | 0 | 0 | 0.257113 |
| Down | TAF4B | 1 | 0 | 0 | 0.28167 |
| Down | NRBP2 | 1 | 0 | 0 | 0.238756 |
| Down | SYNPO2 | 1 | 0 | 0 | 0.267328 |
| Down | MMS22L | 1 | 0 | 0 | 0.256006 |
| Down | TIPIN | 1 | 0 | 0 | 0.256006 |
| Down | ACOX2 | 1 | 0 | 0 | 0.231323 |
| Down | CCNJL | 1 | 0 | 0 | 0.240079 |
| Down | ZNF564 | 1 | 0 | 0 | 0.242793 |
| Down | COLEC11 | 1 | 0 | 0 | 0.24466 |
| Down | RTEL1 | 1 | 0 | 0 | 0.237785 |
| Down | CHL1 | 1 | 0 | 0 | 0.304543 |
| Down | BCAS1 | 1 | 0 | 0 | 0.304543 |
| Down | DCX | 1 | 0 | 0 | 0.245219 |
| Down | COL6A3 | 1 | 0 | 0 | 0.241643 |
| Down | EGF | 1 | 0 | 0 | 0.253159 |
| Down | ZSCAN20 | 1 | 0 | 0 | 0.242605 |
| Down | IFITM1 | 1 | 0 | 0 | 0.256106 |
| Down | STK10 | 1 | 0 | 0 | 0.256106 |
| Down | HRH1 | 1 | 0 | 0 | 0.256106 |
| Down | PECAM1 | 1 | 0 | 0 | 0.252041 |
| Down | PCK1 | 1 | 0 | 0 | 0.257113 |
| Down | S1PR1 | 1 | 0 | 0 | 0.257113 |
| Down | MYL9 | 1 | 0 | 0 | 0.252714 |
| Down | SLC7A5 | 1 | 0 | 0 | 0.252714 |
| Down | GATA4 | 1 | 0 | 0 | 0.238613 |
| Down | RASSF2 | 1 | 0 | 0 | 0.236417 |
| Down | CDCA7L | 1 | 0 | 0 | 0.24466 |
| Down | ISLR | 1 | 0 | 0 | 0.24466 |
| Down | GPC3 | 1 | 0 | 0 | 0.201677 |
| Down | EFNB2 | 1 | 0 | 0 | 0.233486 |
| Down | HEY2 | 1 | 0 | 0 | 0.245018 |

### MiRNA-hub gene regulatory network construction

The miRNA-hub gene regulatory network was shown in Fig. 5, consisting of 2807 nodes and 21728 interactions. MiRNAs includes hsa-mir-1255b-5p, hsa-mir-499a-5p, hsa-mir-200a-3p, hsa-mir-449c-5p, hsa-mir-520a-5p, hsa-mir-410-3p, hsa-mir-3941, hsa-mir-181d-3p, hsa-mir-181c-5p and hsa-mir-708-5p were with the highest connectivity target genes, targeting 10 hub genes (up regulated: MRPL12, RAB5C, VCAM1, HIF1A and CLU; down regulated: CDK1, CHEK1, TK1, KIT and TOP2A) and are listed in Table 6. **TF-hub gene regulatory network construction**

**Fig. 5.**
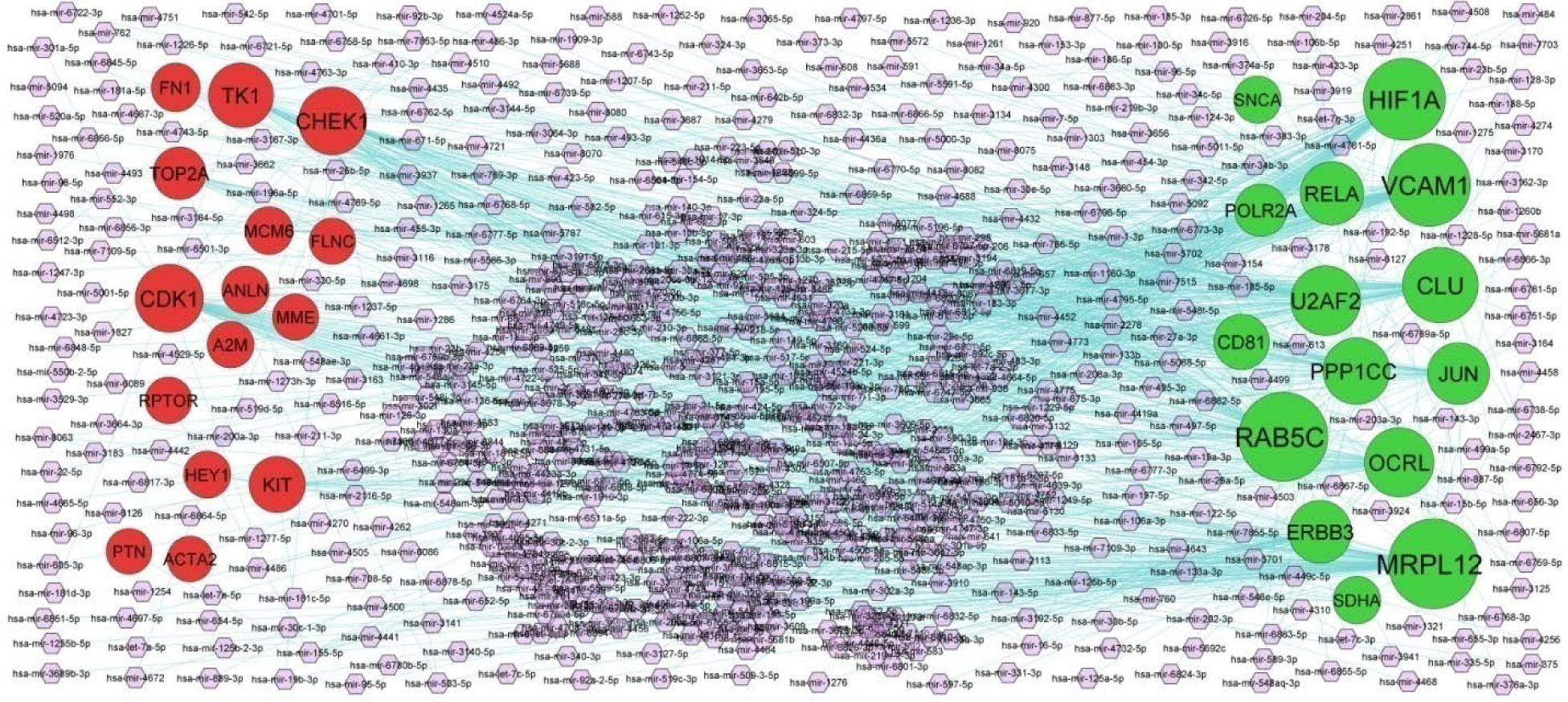
MiRNA - hub gene regulatory network. The purple color diamond nodes represent the key miRNAs; up regulated genes are marked in green; down regulated genes are marked in red.

**Table 6.** miRNA - hub gene and TF – hub gene interaction

| Regulation | Hub Genes | Degree | MicroRNA | Regulation | Hub Genes | Degree | TF |
| --- | --- | --- | --- | --- | --- | --- | --- |
| Up | MRPL12 | 102 | hsa-mir-1255b-5p | Up | JUN | 62 | VDR |
| Up | RAB5C | 100 | hsa-mir-499a-5p | Up | POLR2A | 45 | BACH1 |
| Up | VCAM1 | 86 | hsa-mir-200a-3p | Up | HIF1A | 39 | TFCP2L1 |
| Up | HIF1A | 84 | hsa-mir-449c-5p | Up | MRPL12 | 36 | HOXC9 |
| Up | CLU | 69 | hsa-mir-520a-5p | Up | U2AF2 | 34 | GATA2 |
| Up | U2AF2 | 56 | hsa-mir-3144-5p | Up | RELA | 34 | FOXO3 |
| Up | OCRL | 55 | hsa-mir-95-5p | Up | PPP1CC | 32 | SOX2 |
| Up | PPP1CC | 49 | hsa-mir-4643 | Up | ADRB2 | 27 | GATA1 |
| Up | RELA | 41 | hsa-mir-1303 | Up | ERBB3 | 25 | STAT1 |
| Up | ERBB3 | 40 | hsa-mir-4732-5p | Up | SNCA | 25 | RUNX2 |
| Up | JUN | 38 | hsa-mir-3178 | Up | CD81 | 22 | XRN2 |
| Up | CD81 | 23 | hsa-mir-552-3p | Up | SDHA | 22 | ESR1 |
| Up | POLR2A | 16 | hsa-mir-744-5p | Up | VCAM1 | 16 | EP300 |
| Up | SDHA | 5 | hsa-mir-760 | Up | ALB | 12 | STAT3 |
| Up | SNCA | 4 | hsa-mir-7-5p | Up | OCRL | 9 | TEAD4 |
| Down | CDK1 | 52 | hsa-mir-410-3p | Down | RPTOR | 52 | TBX3 |
| Down | CHEK1 | 52 | hsa-mir-3941 | Down | FN1 | 46 | ATF3 |
| Down | TK1 | 45 | hsa-mir-181d-3p | Down | ANLN | 42 | GATA4 |
| Down | KIT | 25 | hsa-mir-181c-5p | Down | MME | 31 | PAX6 |
| Down | TOP2A | 17 | hsa-mir-708-5p | Down | HEY1 | 28 | FOXP1 |
| Down | FN1 | 8 | hsa-mir-140-3p | Down | TK1 | 28 | SOX11 |
| Down | MCM6 | 6 | hsa-mir-34a-5p | Down | PTN | 27 | SOX17 |
| Down | ANLN | 5 | hsa-mir-497-5p | Down | CHEK1 | 27 | HOXB4 |
| Down | HEY1 | 4 | hsa-mir-30b-5p | Down | TOP2A | 25 | RUNX1 |
| Down | RPTOR | 3 | hsa-mir-27a-3p | Down | MCM6 | 24 | HNF4A |
| Down | A2M | 3 | hsa-mir-98-5p | Down | FLNC | 21 | RNF2 |
| Down | MME | 2 | hsa-mir-1-3p | Down | CDK1 | 19 | FOXM1 |
| Down | FLNC | 2 | hsa-mir-454-3p | Down | RIPK3 | 16 | NACC1 |
| Down | PTN | 1 | hsa-mir-155-5p | Down | A2M | 14 | FOXA2 |
| Down | ACTA2 | 1 | hsa-mir-484 | Down | ACTA2 | 14 | SMAD4 |

The TF-hub gene regulatory network was shown in Fig. 6, consisting of 625 nodes and 10244 interactions. TFs includes VDR, BACH1, TFCP2L1, HOXC9, GATA2, TBX3, ATF3, GATA4, PAX6 and FOXP1 were with the highest connectivity target genes, targeting 10 hub genes (up regulated: JUN, POLR2A, HIF1A, MRPL12 and U2AF2; down regulated: RPTOR, FN1, ANLN, MME and HEY1) and are listed in Table 6.

**Fig. 6.**
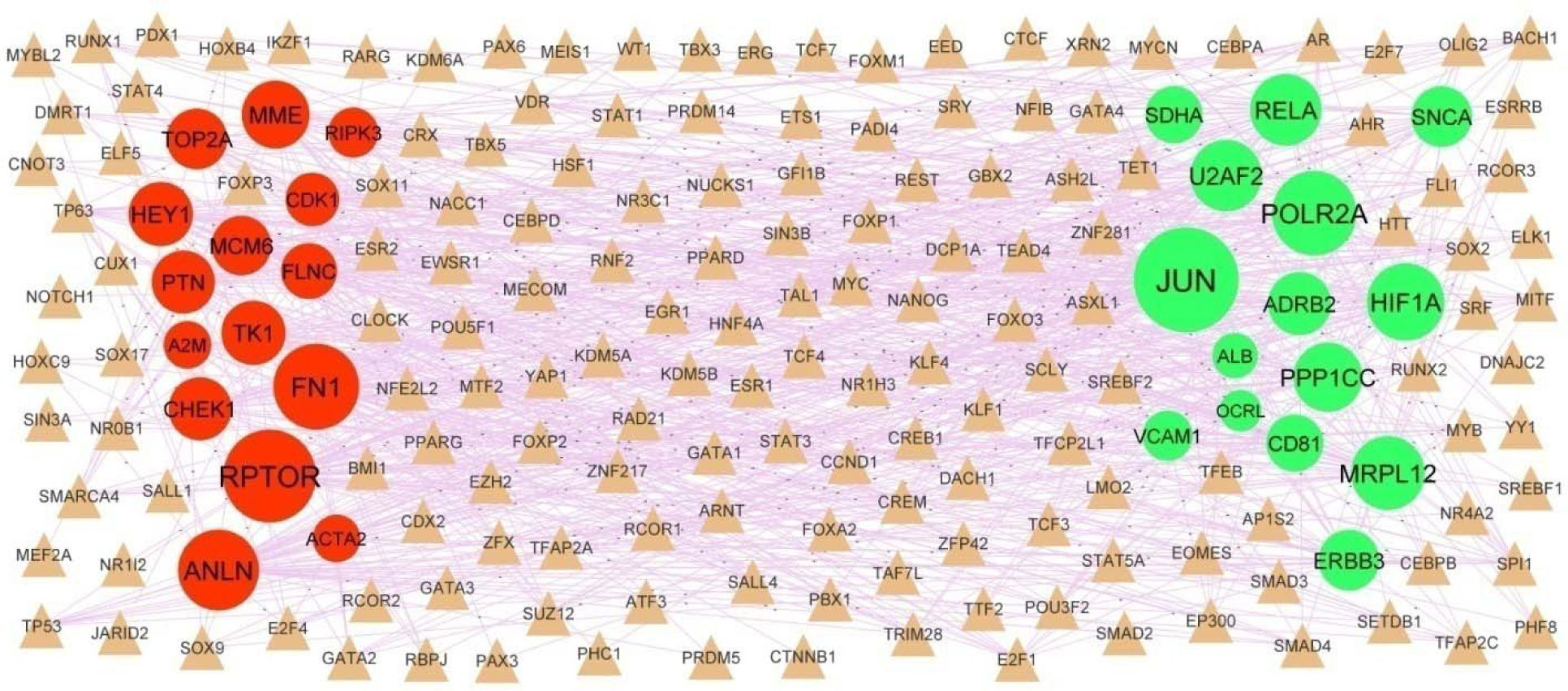
TF - hub gene regulatory network. The brown color triangle nodes represent the key TFs; up regulated genes are marked in green; down regulated genes are marked in red.

### Validation of hub genes by receiver operating characteristic curve (ROC) analysis

Validate the expression of these 10 genes in T2DM and performed ROC analysis in RStudio with pROC package. The analysis results were available in Fig.7. Ten genes (JUN, VCAM1, RELA, U2AF2, ADRB2, FN1, CDK1, TK1, A2M and ACTA2) with AUC more than 0.90 were considered as hub genes, indicating that they have the capability to diagnose T2DM patients with excellent specificity and sensitivity.

**Fig. 7.**
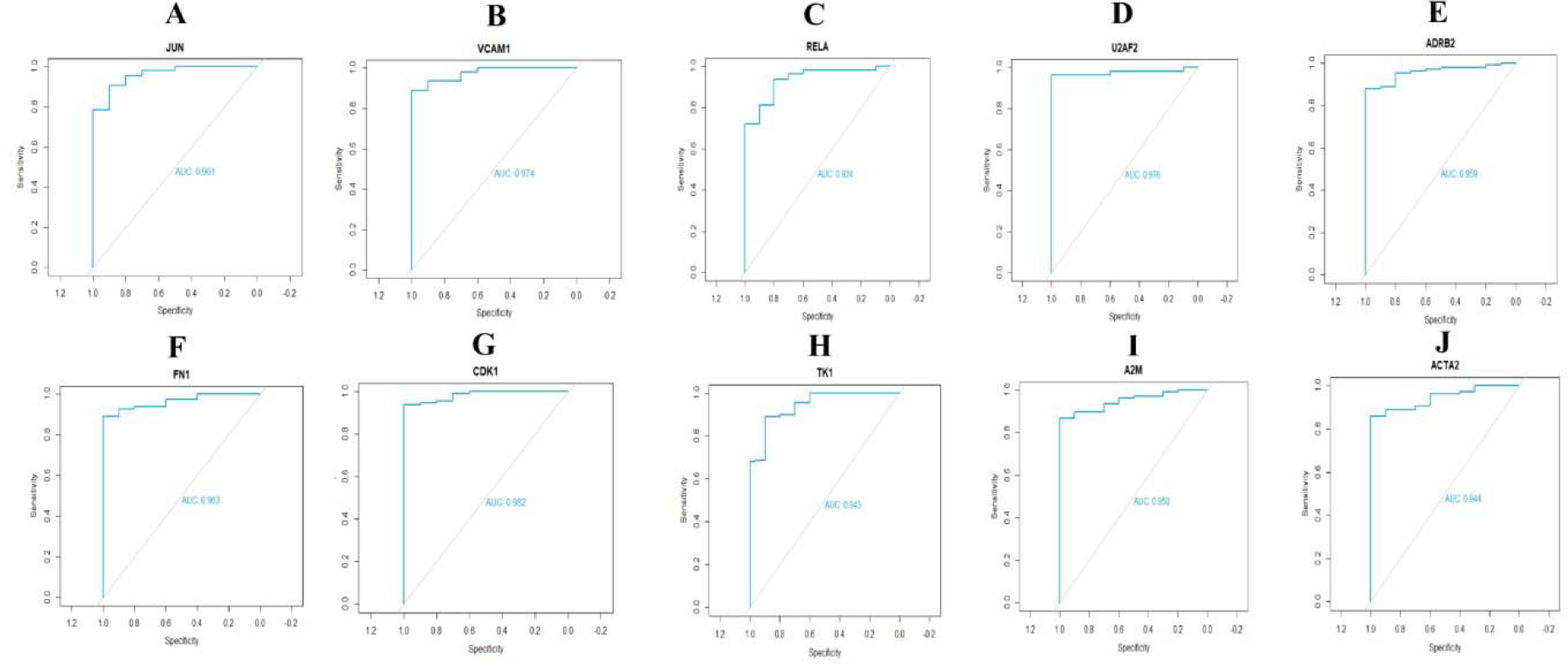
ROC curve validated the sensitivity, specificity of hub genes as a predictive biomarker for T2DM prognosis. A) JUN B) VCAM1C) RELA D) U2AF2 E) ADRB2 F) FN1 G) CDK1 H) TK1 I) A2M J) ACTA2

### Detection of the mRNA expression of the hub genes by RT-PCR

To further verify the expression level of hub genes in T2DM (pancreatic beta cell line MIN6 culture treated with streptozotocin) and normal (pancreatic beta cell line MIN6 culture), RT-PCR was performed to calculate the mRNA levels of the ten hub genes identified in the present study (JUN, VCAM1, RELA, U2AF2, ADRB2, FN1, CDK1, TK1, A2M and ACTA2) in T2DM. As illustrated in Fig. 8, the expression of JUN, VCAM1, RELA, U2AF2 and ADRB2 were significantly up regulated in T2DM compared with normal, while FN1, CDK1, TK1, A2M and ACTA2 were significantly up regulated in T2DM compared with normal. The current RT-PCR results were in line with the previous bioinformatics analysis, suggesting that these hub genes might be associated to the molecular mechanism underlying T2DM.

**Fig. 8.**
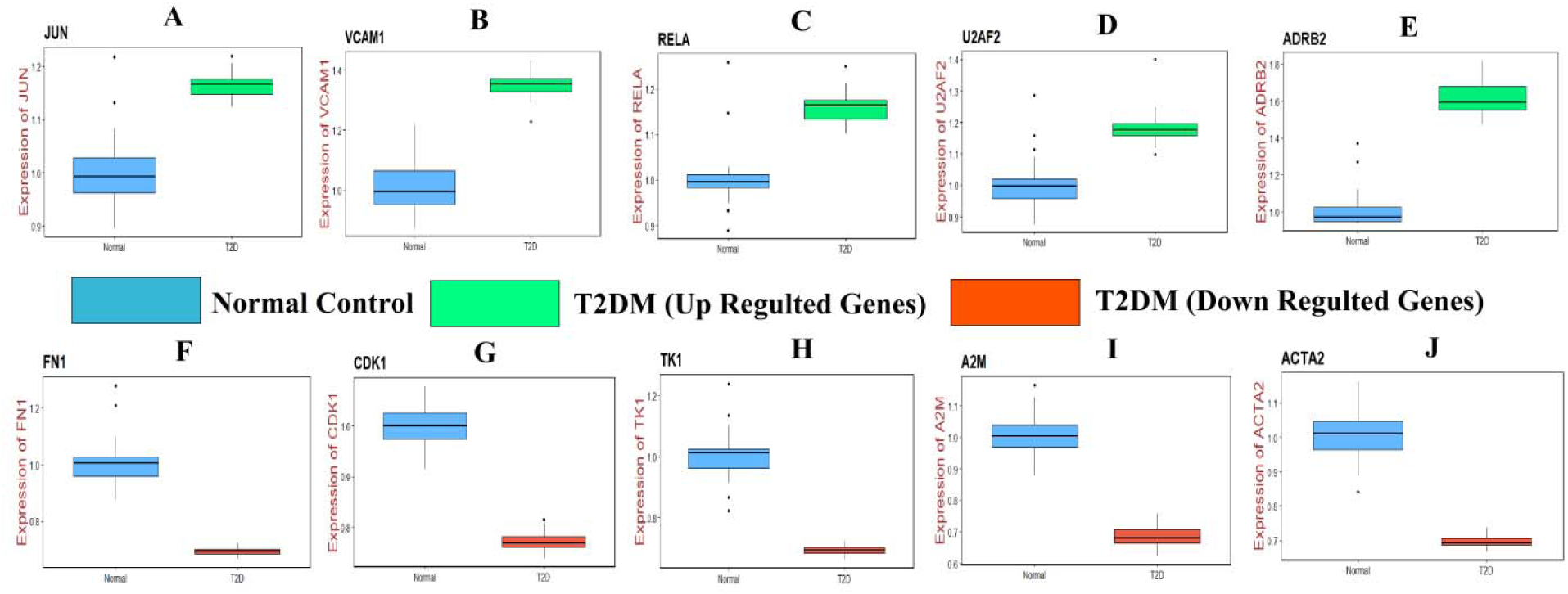
Validation of hub genes by RT-PCR. A) JUN B) VCAM1C) RELA D) U2AF2 E) ADRB2 F) FN1 G) CDK1 H) TK1 I) A2M J) ACTA2

## Discussion

T2DM is a complex metabolic disorder characterized by defective glucose uptake in muscle and fat, varied glucose-induced insulin secretion, and heightened hepatic glucose production. It is largely ascribed to insulin resistance and leads in a heavy health and economic burden. Therefore, it is important to study the molecular mechanism of T2DM. In this investigation, we tried to identify the important genes related to T2DM by comparing the differences in expression profiling by high throughput sequencing between the T2DM and normal controls. A total of 925 DEGs (447 up regulated genes and 478 down regulated genes) were screened between T2DM and control samples. Emmens et al [39] found that PENK (proenkephalin) expression was up regulated in cardiovascular diseases, but this gene might be responsible for progression of T2DM. Recent studies have proposed that the SAA1 [40], SUCNR1 [41], CRP (C-reactive protein) [42], APCS (amyloid P component, serum) [43], PCK1 [44], IGF2BP3 [45], POSTN (periostin) [46] and COL6A3 [47] are associated with progression of T2DM. CARTPT (CART prepropeptide) [48], SEMA3C [49] and TDO2 [50] were proved to participate in the development of obesity, but these genes might be linked with progression of T2DM. SOX11 is associated with progression of neurological disorders [51], but this gene might be liable for advancement of T2DM.

The functional enrichment analysis indicated that the DEGs were primarily involved in the defense response, secretory vesicle, identical protein binding, cell adhesion, extracellular matrix, structural molecule activity, neutrophil degranulation and extracellular matrix organization. Neutrophil degranulation [52], innate immune system [53], platelet degranulation [54], extracellular matrix organization [55], diseases of glycosylation [56], platelet activation, signaling and aggregation [57], hemostasis [58], secretion [59], secretory vesicle [60], transmembrane transporter activity [61], cell adhesion [62], localization of cell [63], extracellular matrix [55], intrinsic component of plasma membrane [64], structural molecule activity [65], signaling receptor binding [66], have been highly noted in T2DM. Reports indicate that HIF1A [67], HLA-DRB1 [68], CHI3L1 [69], ADORA2A [70], ADRB2 [71], CLU (clusterin) [72], AGT (angiotensinogen) [73], VCAM1 [74], PPARA (peroxisome proliferator activated receptor alpha) [75], APOL1 [76], ZFP36 [77], PPM1B [78], SOCS1 [79], SNCA (synuclein alpha) [80], CTSS (cathepsin S) [81], IL6R [82], CFB (complement factor B) [83], DEFB1 [84], VNN1 [85], RAB27A [86], DPP4 [87], RARRES2 [88], CASP1 [89], LCN2 [90], REG3A [91], CD74 [92], PCSK2 [93], CHGB (chromogranin B) [94], TTR (transthyretin) [95], LRG1 [96], ALB (albumin) [97], DPP7 [98], APOH (apolipoprotein H) [99], CTSD (cathepsin D) [100], GCG (glucagon) [101], KCNQ1 [102], NR4A1 [103], PLIN5 [104], ALDH2 [105], ANG (angiogenin) [106], CLDN7 [107], PRLR (prolactin receptor) [108], SOD2 [109], MLXIPL (MLX interacting protein like) [110], CTSD (cathepsin D) [111], PECAM1 [112], ADA (adenosine deaminase) [113], MFGE8 [114], COL1A1 [115], COL3A1 [116], NID2 [117], ARG1 [118], CD93 [119], IGF2 [120], IL18 [121], LAMA1 [122], HPSE (heparanase) [123], BMP4 [124], CXCR4 [125], KDR (kinase insert domain receptor) [126], ESAM (endothelial cell adhesion molecule) [127], THBS1 [128], CD34 [129], SERPINE1 [130], WNT5B [131], MGP (matrix Gla protein) [132], MMP3 [133], MMP9 [134], MMP11 [135], IL16 [136], TNFRSF11B [137], TIMP3 [138] and CAPN3 [139] were found in T2DM. [140], HLA-B [141], HLA-C [142], HLA-DPB1 [143], HLA-E [144], ERBB3 [145], MFAP4 [146] and JAK3 [147] have been reported significantly expressed in type 1 diabetes mellitus, but these genes might be involved in T2DM progression. HLA-DRA [148], SERPINA1 [149], ABHD12 [150], IMPA2 [151], ARSA (arylsulfatase A) [152], LRFN5 [153], PLXNA4 [154], CHL1 [155], ITPKB (inositol-trisphosphate 3-kinase B) [156], PTN (pleiotrophin) [157], LAMA2 [158], CDH6 [159] and A2M [160] have been shown to have an important role in neurological disorders, but these genes might be associated with progression of T2DM. HLA-F [161], HLA-H [162], FGA (fibrinogen alpha chain) [163], HSPA1B [164], MRC1 [165], DAB2IP [166], KCNJ8 [167], KLKB1 [168], CXCL2 [169], SERPINE2 [170], ADH1C [171], AMBP (alpha-1-microglobulin/bikunin precursor) [172], NR4A2 [173], TYMP (thymidine phosphorylase) [174], TFRC (transferrin receptor) [175], PLAU (plasminogen activator, urokinase) [176], COL6A2 [177], COL15A1 [178], ABI3BP [179], NEXN (nexilin F-actin binding protein) [180], S1PR1 [181], THY1 [182], COL4A1 [183], COL5A2 [184], ADAMTS2 [185], ECM1 [186] and LTBP2 [187] have been found to be differentially expressed in cardiovascular diseases, but these genes might be linked with progression of T2DM. CCL20 [188], CRH (corticotropin releasing hormone) [189], SPP1 [190], LDLR (low density lipoprotein receptor) [191], RORA (RAR related orphan receptor A) [192], LYZ (lysozyme) [193], PTPRN2 [194], DAPK2 [195], OIP5 [196], PON3 [197], NR4A3 [198], VCAN (versican) [199], CNTNAP2 [200], IL1RAP [201], GLI2 [202], CDH13 [203], AEBP1 [204], BGN (biglycan) [205], LOX (lysyl oxidase) [206], IL1RL1 [207] and LUM (lumican) [208] were found to be involved in advancement of obesity, but these genes might be key for development of T2DM. C4BPA [209], KYNU (kynureninase) [210], ORM1 [211], ARSA (arylsulfatase A) [212], CYB5R3 [213], MAT1A [214], SDC4 [215], ASL (argininosuccinate lyase) [216], SLC4A4 [217], EPHB6 [218], SPARCL1 [219], THBS2 [220], EFNB2 [221] and CD248 [222] were revealed to be associated with hypertension, but these genes might be involved in progression of T2DM. GSDMD (gasdermin D) [223], UNC5B [224] and PDGFB (platelet derived growth factor subunit B) [225] have been reported significantly expressed in diabetic nephropathy.

A PPI network and modules was constructed and analyzed with DEGs, and then hub genes were identified. Gregg et al [226] revealed that CDK1 involvement in of obesity, but this gene might be associated with advancement of T2DM. ACTA2 was revealed to be associated with development of T2DM [227]. JUN (jun proto-oncogene, AP-1 transcription factor subunit), RELA (RELA proto-oncogene, NF-kB subunit), U2AF2, TK1, RAB5C, OCRL (OCRL inositol polyphosphate-5-phosphatase), COL5A1, FN1, SPARC (secreted protein acidic and cysteine rich), TGFBI (transforming growth factor beta induced) and COL1A2 might be the novel biomarkers for the advancement of T2DM.

In this investigation, miRNA-hub gene regulatory network and TF-hub gene regulatory network were constructed and analyzed. Demirsoy et al [228], Hourigan et al [229], Safar et al [230], Zmuda et al [231], Boese et al [232] and Suhara et al [233] found that hsa-mir-200a-3p, hsa-mir-181c-5p, VDR (vitamin D receptor), ATF3, PAX6 and RPTOR (regulatory associated protein of MTOR complex 1) were participating in the occurrence and development of T2DM. Cabiati et al [234] and Patankar et al [235] proved that hsa-mir-410-3p and GATA4 mediate the development of obesity, but these genes might be associated with development of T2DM. Hsa-mir-3941 [236], BACH1 [237], GATA2 [238], FOXP1 [239] and HEY1 [240] were found to be involved in the process of cardiovascular diseases, but these genes might be associated with development of T2DM. Hsa-mir-708-5p was revealed to be correlated with disease outcome in patients with neurological disorders [241], but this gene might be linked with development of T2DM. A previous study found that MRPL12 [242] and MME (membrane metalloendopeptidase) [243] expression could enhance the diabetic nephropathy. A previous study showed that CHEK1 gene is involved in the hypertension [244], but this gene might be associated in development of T2DM. Hsa-mir-1255b-5p, hsa-mir-499a-5p, hsa-mir-449c-5p, hsa-mir-520a-5p, hsa-mir-181d-3p, TFCP2L1, HOXC9, TBX3, KIT (KIT proto-oncogene, receptor tyrosine kinase), TOP2A, POLR2A and ANLN (anillin actin binding protein) might be the novel biomarkers for the advancement of T2DM.

In conclusion, the current investigation exhibits the global profile of DEGs and relative signaling pathways that might participate in the initiation and advancement of T2DM. In the pathogenesis of T2DM, the possible hub genes are JUN, VCAM1, RELA, U2AF2, ADRB2, FN1, CDK1, TK1, A2M and ACTA2. These results could help elucidate the molecular mechanism underlying T2DM pathogenesis and provide potential targets for T2DM therapy. Therefore, more experiments are required to ascertain the clinical value of the identified hub genes as biomarkers and the underlying molecular mechanism.

## Acknowledgement

I thank Jonathan Schug, University of Pennsylvania, School of Medicine, Next-Generation Sequencing Core, Philadelphia, Pennsylvania, USA, very much, the author who deposited their profiling by high throughput sequencing dataset GSE154126, into the public GEO database.

## Conflict of interest

The authors declare that they have no conflict of interest.

## Ethical approval

This article does not contain any studies with human participants or animals performed by any of the authors.

## Informed consent

No informed consent because this study does not contain human or animals participants.

## Availability of data and materials

The datasets supporting the conclusions of this article are available in the GEO (Gene Expression Omnibus) (https://www.ncbi.nlm.nih.gov/geo/) repository. [(GSE154126)

(https://www.ncbi.nlm.nih.gov/geo/query/acc.cgi?acc=GSE154126)]

## Consent for publication

Not applicable.

## Competing interests

The authors declare that they have no competing interests.

## Author Contributions

B. V. - Writing original draft, and review and editing

C. V. - Software and investigation

## Notes

### Competing Interest Statement

The authors have declared no competing interest.

### Summary of Updates

Improved results of RT-PCR included and some typographical mistakes are rectified

## References

1. Jing X, Chen J, Dong Y, Han D, Zhao H, Wang X, Gao F, Li C, Cui Z, Liu Y, et al. Related factors of quality of life of type 2 diabetes patients: a systematic review and meta-analysis. Health Qual Life Outcomes. 2018;16(1):189. doi:10.1186/s12955-018-1021-9

2. Moran C, Beare R, Wang W, Callisaya M, Srikanth V; Alzheimer’s Disease Neuroimaging Initiative (ADNI). Type 2 diabetes mellitus, brain atrophy, and cognitive decline. Neurology. 2019;92(8):e823-e830. doi:10.1212/WNL.0000000000006955

3. Cannon CP, Perkovic V, Agarwal R, Baldassarre J, Bakris G, Charytan DM, de Zeeuw D, Edwards R, Greene T, Heerspink HJL, et al Evaluating the Effects of Canagliflozin on Cardiovascular and Renal Events in Patients With Type 2 Diabetes Mellitus and Chronic Kidney Disease According to Baseline HbA1c, Including Those With HbA1c <7%: Results From the CREDENCE Trial. Circulation. 2020;141(5):407-410. doi:10.1161/CIRCULATIONAHA.119.044359

4. Choi EY, Park SE, Lee SC, Koh HJ, Kim SS, Byeon SH, Kim M. et al. Association Between Clinical Biomarkers and Optical Coherence Tomography Angiography Parameters in Type 2 Diabetes Mellitus. Invest Ophthalmol Vis Sci. 2020;61(3):4. doi:10.1167/iovs.61.3.4

5. Kenny HC, Abel ED. Heart Failure in Type 2 Diabetes Mellitus. Circ Res. 2019;124(1):121–141. doi:10.1161/CIRCRESAHA.118.311371

6. García-Chapa EG, Leal-Ugarte E, Peralta-Leal V, Durán-González J, Meza-Espinoza JP. Genetic Epidemiology of Type 2 Diabetes in Mexican Mestizos. Biomed Res Int. 2017;2017:3937893. doi:10.1155/2017/3937893

7. Dendup T, Feng X, Clingan S, Astell-Burt T. Environmental Risk Factors for Developing Type 2 Diabetes Mellitus: A Systematic Review. Int J Environ Res Public Health. 2018;15(1):78. doi:10.3390/ijerph15010078

8. Sun Y, Gao HY, Fan ZY, He Y, Yan YX. Metabolomics Signatures in Type 2 Diabetes: A Systematic Review and Integrative Analysis. J Clin Endocrinol Metab. 2020;105(4):dgz240. doi:10.1210/clinem/dgz240

9. Kahn SE, Hull RL, Utzschneider KM. Mechanisms linking obesity to insulin resistance and type 2 diabetes. Nature. 2006;444(7121):840–846. doi:10.1038/nature05482

10. Climie RE, van Sloten TT, Bruno RM, Taddei S, Empana JP, Stehouwer CDA, Sharman JE, Boutouyrie P, Laurent S. Macrovasculature and Microvasculature at the Crossroads Between Type 2 Diabetes Mellitus and Hypertension. Hypertension. 2019;73(6):1138–1149. doi:10.1161/HYPERTENSIONAHA.118.11769

11. Mofo Mato EP, Guewo-Fokeng M, Essop MF, Owira PMO. Genetic polymorphisms of organic cation transporter 1 (OCT1) and responses to metformin therapy in individuals with type 2 diabetes: A systematic review. Medicine (Baltimore). 2018;97(27):e11349. doi:10.1097/MD.0000000000011349

12. Guo X, Li C, Wu J, Mei Q, Liu C, Sun W, Xu L, Fu S. The association of TNF-α -308G/A and -238G/A polymorphisms with type 2 diabetes mellitus: doi:10.1042/BSR20191301

13. Ding W, Xu L, Zhang L, Han Z, Jiang Q, Wang Z, Jin S. Meta-analysis of association between TCF7L2 polymorphism rs7903146 and type 2 diabetes mellitus. BMC Med Genet. 2018;19(1):38. doi:10.1186/s12881-018-0553-5

14. Rizvi S, Raza ST, Mahdi F, Singh SP, Rajput M, Rahman Q. Genetic polymorphisms in KCNJ11 (E23K, rs5219) and SDF-1β (G801A, rs1801157) genes are associated with the risk of type 2 diabetes mellitus. Br J Biomed Sci. 2018;75(3):139–144. doi:10.1080/09674845.2018.1473939

15. Bregar D, Cilenšek I, Mankoč S, Reschner A, Petrovič D, Globočnik Petrovič M. The joint effect of the endothelin receptor B gene (EDNRB) polymorphism rs10507875 and nitric oxide synthase 3 gene (NOS3) polymorphism rs869109213 in Slovenian patients with type 2 diabetes mellitus and diabetic retinopathy. Bosn J Basic Med Sci. 2018;18(1):80–86. doi:10.17305/bjbms.2017.2244

16. Huang X, Liu G, Guo J, Su Z. The PI3K/AKT pathway in obesity and type 2 diabetes. Int J Biol Sci. 2018;14(11):1483–1496. doi:10.7150/ijbs.27173

17. Akbari M, Hassan-Zadeh V. IL-6 signalling pathways and the development of type 2 diabetes. Inflammopharmacology. 2018;26(3):685–698. doi:10.1007/s10787-018-0458-0

18. Jin T, Liu L. The Wnt signaling pathway effector TCF7L2 and type 2 diabetes mellitus. Mol Endocrinol. 2008;22(11):2383–2392. doi:10.1210/me.2008-0135

19. Karolina DS, Armugam A, Tavintharan S, et al. MicroRNA 144 impairs insulin signaling by inhibiting the expression of insulin receptor substrate 1 in type 2 diabetes mellitus. PLoS One. 2011;6(8):e22839. doi:10.1371/journal.pone.0022839

20. Zuo Y. The role of adiponectin gene mediated by NF-κB signaling pathway in the pathogenesis of type 2 diabetes. Eur Rev Med Pharmacol Sci. 2018;22(4):1106–1112. doi:10.26355/eurrev_201802_14398

21. Yang F, Chen Y, Xue Z, Lv Y, Shen L, Li K, Zheng P, Pan P, Feng T, Jin L, et al. High-Throughput Sequencing and Exploration of the lncRNA-circRNA-miRNA-mRNA Network in Type 2 Diabetes Mellitus. Biomed Res Int. 2020;2020:8162524. doi:10.1155/2020/8162524

22. Avrahami D, Wang YJ, Schug J, Feleke E, Gao L, Liu C; HPAP Consortium, Naji A, Glaser B, Kaestner KH. Single-cell transcriptomics of human islet ontogeny defines the molecular basis of β-cell dedifferentiation in T2D. Mol Metab. 2020;42:101057. doi:10.1016/j.molmet.2020.101057

23. Clough E, Barrett T. The Gene Expression Omnibus Database. Methods Mol Biol. 2016;1418:93–110. doi:10.1007/978-1-4939-3578-9_5

24. Ritchie ME, Phipson B, Wu D, Hu Y, Law CW, Shi W, Smyth GK. limma powers differential expression analyses for RNA-sequencing and microarray studies. Nucleic Acids Res. 2015;43(7):e47. doi:10.1093/nar/gkv007

25. Thomas PD. The Gene Ontology and the Meaning of Biological Function. Methods Mol Biol. 2017;1446:15–24. doi:10.1007/978-1-4939-3743-1_2

26. Fabregat A, Jupe S, Matthews L, Sidiropoulos K, Gillespie M, Garapati P, Haw R, Jassal B, Korninger F, May B et al The Reactome Pathway Knowledgebase. Nucleic Acids Res. 2018;46(D1):D649–D655. doi:10.1093/nar/gkx1132

27. Chen J, Bardes EE, Aronow BJ, Jegga AG. ToppGene Suite for gene list enrichment analysis and candidate gene prioritization. Nucleic Acids Res. 2009;37(Web Server issue):W305–W311. doi:10.1093/nar/gkp427

28. Orchard S, Ammari M, Aranda B, Breuza L, Briganti L, Broackes-Carter F, Campbell NH, Chavali G, Chen C, del-Toro N, et al. The MIntAct project--IntAct as a common curation platform for 11 molecular interaction databases. Nucleic Acids Res. 2014;42(Database issue):D358-D363. doi:10.1093/nar/gkt1115

29. Shannon P, Markiel A, Ozier O, Baliga NS, Wang JT, Ramage D, Amin N, Schwikowski B, Ideker T Cytoscape: a software environment for integrated models of biomolecular interaction networks. Genome Res 2003;13(11):2498–2504. doi:10.1101/gr.1239303

30. Przulj N, Wigle DA, Jurisica I. Functional topology in a network of protein interactions. Bioinformatics. 2004;20(3):340–348. doi:10.1093/bioinformatics/btg415

31. Nguyen TP, Liu WC, Jordán F. Inferring pleiotropy by network analysis: linked diseases in the human PPI network. BMC Syst Biol. 2011;5:179. doi:10.1186/1752-0509-5-179

32. Shi Z, Zhang B. Fast network centrality analysis using GPUs. BMC Bioinformatics. 2011;12:149. doi:10.1186/1471-2105-12-149

33. Fadhal E, Gamieldien J, Mwambene EC. Protein interaction networks as metric spaces: a novel perspective on distribution of hubs. BMC Syst Biol. 2014;8:6. doi:10.1186/1752-0509-8-6

34. Zaki N, Efimov D, Berengueres J. Protein complex detection using interaction reliability assessment and weighted clustering coefficient. BMC Bioinformatics. 2013;14:163. doi:10.1186/1471-2105-14

35. Fan Y, Xia J (2018) miRNet-Functional Analysis and Visual Exploration of miRNA-Target Interactions in a Network Context. Methods Mol Biol 1819:215–233. doi:10.1007/978-1-4939-8618-7_10

36. Zhou G, Soufan O, Ewald J, Hancock REW, Basu N, Xia J (2019) NetworkAnalyst 3.0: a visual analytics platform for comprehensive gene expression profiling and meta-analysis. Nucleic Acids Res 47:W234–W241. doi:10.1093/nar/gkz240

37. Robin X, Turck N, Hainard A, Tiberti N, Lisacek F, Sanchez JC, Müller M. pROC: an open-source package for R and S+ to analyze and compare ROC curves. BMC Bioinformatics 2011;12:77. doi:10.1186/1471-2105-12-77

38. Livak KJ, Schmittgen TD Analysis of relative gene expression data using real-time quantitative PCR and the 2(-Delta Delta C(T)) Method. Methods 2001;25:402–408. doi:10.1006/meth.2001.1262

39. Emmens JE, Ter Maaten JM, Damman K, van Veldhuisen DJ, de Boer RA, Struck J, Bergmann A, Sama IE, Streng KW, Anker SD, et al. Proenkephalin, an Opioid System Surrogate, as a Novel Comprehensive Renal Marker in Heart Failure. Circ Heart Fail. 2019;12(5):e005544. doi:10.1161/CIRCHEARTFAILURE.118.005544

40. Wang Y, Cao F, Wang Y, Yu G, Jia BL. Silencing of SAA1 inhibits palmitate-or high-fat diet induced insulin resistance through suppression of the NF-κB pathway. Mol Med. 2019;25(1):17. doi:10.1186/s10020-019-0075-4

41. Du B, Jia X, Tian W, Yan X, Wang N, Cai D, Li X, Zhang H, Jin M, Wu N, et al. Associations of SUCNR1, GRK4, CAMK1D gene polymorphisms and the susceptibility of type 2 diabetes mellitus and essential hypertension in a northern Chinese Han population. J Diabetes Complications. 2021;35(1):107752. doi:10.1016/j.jdiacomp.2020.107752

42. Yeo ES, Hwang JY, Park JE, Choi YJ, Huh KB, Kim WY. Tumor necrosis factor (TNF-alpha) and C-reactive protein (CRP) are positively associated with the risk of chronic kidney disease in patients with type 2 diabetes. Yonsei Med J. 2010;51(4):519–525. doi:10.3349/ymj.2010.51.4.519

43. Scarale MG, Copetti M, Garofolo M, Fontana A, Salvemini L, De Cosmo S, Lamacchia O, Penno G, Trischitta V, Menzaghi C.The Synergic Association of hs-CRP and Serum Amyloid P Component in Predicting All-Cause Mortality in Patients With Type 2 Diabetes. Diabetes Care. 2020;43(5):1025–1032. doi:10.2337/dc19-2489

44. Rees SD, Britten AC, Bellary S, O’Hare JP, Kumar S, Barnett AH, Kelly MA. The promoter polymorphism -232C/G of the PCK1 gene is associated with type 2 diabetes in a UK-resident South Asian population. BMC Med Genet. 2009;10:83. doi:10.1186/1471-2350-10-83

45. Rodriguez S, Eiriksdottir G, Gaunt TR, Harris TB, Launer LJ, Gudnason V, Day IN. IGF2BP1, IGF2BP2 and IGF2BP3 genotype, haplotype and genetic model studies in metabolic syndrome traits and diabetes. Growth Horm IGF Res. 2010;20(4):310–318. doi:10.1016/j.ghir.2010.04.002

46. Luo Y, Qu H, Wang H, Wei H, Wu J, Duan Y, Liu D, Deng H. Plasma Periostin Levels Are Increased in Chinese Subjects with Obesity and Type 2 Diabetes and Are Positively Correlated with Glucose and Lipid Parameters. Mediators Inflamm. 2016;2016:6423637. doi:10.1155/2016/6423637

47. Dankel SN, Svärd J, Matthä S, Claussnitzer M, Klöting N, Glunk V, Fandalyuk Z, Grytten E, Solsvik MH, Nielsen HJ, et al. COL6A3 expression in adipocytes associates with insulin resistance and depends on PPARγ and adipocyte size. Obesity (Silver Spring). 2014;22(8):1807–1813. doi:10.1002/oby.20758

48. Lisa Y, Sook- HF, Yee- HS. Association of the Cocaine-and Amphetamine-Regulated Transcript Prepropeptide Gene (CARTPT) rs2239670 Variant with Obesity among Kampar Health Clinic Patrons, Malaysia. Malays J Med Sci. 2012;19(1):43–51.

49. Nam JS, Ahn CW, Park HJ, Kim YS. Semaphorin 3 C is a Novel Adipokine Representing Exercise-Induced Improvements Metabolism in Metabolically Healthy Obese Young Males. Sci Rep. 2020;10(1):10005. doi:10.1038/s41598-020-67004-7

50. Poulain-Godefroy O, Eury E, Leloire A, Hennart B, Guillemin GJ, Allorge D, Froguel P.Induction of TDO2 and IDO2 in Liver by High-Fat Feeding in Mice: Discrepancies with Human Obesity. Int J Tryptophan Res. 2013;6(Suppl 1):29–37. doi:10.4137/IJTR.S11717

51. Sun CP, Sun D, Luan ZL, Dai X, Bie X, Ming WH, Sun XW, Huo XX, Lu TL, Zhang D. Association of SOX11 Polymorphisms in distal 3’UTR with Susceptibility for Schizophrenia. J Clin Lab Anal. 2020;34(8):e23306. doi:10.1002/jcla.23306

52. Gorudko IV, Kostevich VA, Sokolov AV, Shamova EV, Buko IV, Konstantinova EE, Vasiliev VB, Cherenkevich SN, Panasenko OM. Functional activity of neutrophils in diabetes mellitus and coronary heart disease: role of myeloperoxidase in the development of oxidative stress. Bull Exp Biol Med. 2012;154(1):23–26. doi:10.1007/s10517-012-1865-7

53. Creely SJ, McTernan PG, Kusminski CM, Fisher fM, Da Silva NF, Khanolkar M, Evans M, Harte AL, Kumar S. Lipopolysaccharide activates an innate immune system response in human adipose tissue in obesity and type 2 diabetes. Am J Physiol Endocrinol Metab. 2007;292(3):E740–E747. doi:10.1152/ajpendo.00302.2006

54. Fateh-Moghadam S, Li Z, Ersel S, Reuter T, Htun P, Plöckinger U, Bocksch W, Dietz R, Gawaz M. Platelet degranulation is associated with progression of intima-media thickness of the common carotid artery in patients with diabetes mellitus type 2. Arterioscler Thromb Vasc Biol. 2005;25(6):1299–1303. doi:10.1161/01.ATV.0000165699.41301.c5

55. Liu JH, Chen Y, Zhen Z, Ho LM, Tsang A, Yuen M, Lam K, Tse HF, Yiu KH. Relationship of biomarkers of extracellular matrix with myocardial function in Type 2 diabetes mellitus. Biomark Med. 2017;11(7):569–578. doi:10.2217/bmm-2017-0044

56. Kundu D, Roy A, Mandal T, Bandyopadhyay U, Ghosh E, Ray D. Relation of microalbuminuria to glycosylated hemoglobin and duration of type 2 diabetes. Niger J Clin Pract. 2013;16(2):216–220. doi:10.4103/1119-3077.110159

57. Suslova TE, Sitozhevskii AV, Ogurkova ON, Kravchenko ES, Kologrivova IV, Anfinogenova Y, Karpov RS. Platelet hemostasis in patients with metabolic syndrome and type 2 diabetes mellitus: cGMP-and NO-dependent mechanisms in the insulin-mediated platelet aggregation. Front Physiol. 2015;5:501. doi:10.3389/fphys.2014.00501

58. Palella E, Cimino R, Pullano SA, Fiorillo AS, Gulletta E, Brunetti A, Foti DP, Greco M. Laboratory Parameters of Hemostasis, Adhesion Molecules, and Inflammation in Type 2 Diabetes Mellitus: Correlation with Glycemic Control. Int J Environ Res Public Health. 2020;17(1):300. doi:10.3390/ijerph17010300

59. Cortez-Navarrete M, Martínez-Abundis E, Pérez-Rubio KG, González-Ortiz M, Méndez-Del Villar M. Momordica charantia Administration Improves Insulin Secretion in Type 2 Diabetes Mellitus. J Med Food. 2018;21(7):672–677. doi:10.1089/jmf.2017.0114

60. Xiao Y, Zheng L, Zou X, Wang J, Zhong J, Zhong T. Extracellular vesicles in type 2 diabetes mellitus: key roles in pathogenesis, complications, and therapy. J Extracell Vesicles. 2019;8(1):1625677. doi:10.1080/20013078.2019.1625677

61. Saha S. Association between the membrane transporter proteins and type 2 diabetes mellitus. Expert Rev Clin Pharmacol. 2020;13(3):287–297. doi:10.1080/17512433.2020.1729125

62. Hegazy GA, Awan Z, Hashem E, Al-Ama N, Abunaji AB. Levels of soluble cell adhesion molecules in type 2 diabetes mellitus patients with macrovascular complications. J Int Med Res. 2020;48(4):300060519893858. doi:10.1177/0300060519893858

63. Tang X, Hu X, Yang X, Fan Y, Li Y, Hu W, Liao Y, Zheng MC, Peng W, Gao L. Predicting diabetes mellitus genes via protein-protein interaction and protein subcellular localization information. BMC Genomics. 2016;17 Suppl 4(Suppl 4):433. doi:10.1186/s12864-016-2795-y

64. Weijers RN. Lipid composition of cell membranes and its relevance in type 2 diabetes mellitus. Curr Diabetes Rev. 2012;8(5):390–400. doi:10.2174/157339912802083531

65. Nicolle E, Souard F, Faure P, Boumendjel A. Flavonoids as promising lead compounds in type 2 diabetes mellitus: molecules of interest and structure-activity relationship. Curr Med Chem. 2011;18(17):2661–2672. doi:10.2174/092986711795933777

66. Alghamdi F, Guo M, Abdulkhalek S, Crawford N, Amith SR, Szewczuk MR. A novel insulin receptor-signaling platform and its link to insulin resistance and type 2 diabetes. Cell Signal. 2014;26(6):1355–1368. doi:10.1016/j.cellsig.2014.02.015

67. Huang Y, Jin L, Yu H, Jiang G, Tam CHT, Jiang S, Zheng C, Jiang F, Zhang R, Zhang H, et al. SNPs in PRKCA-HIF1A-GLUT1 are associated with diabetic kidney disease in a Chinese Han population with type 2 diabetes. Eur J Clin Invest. 2020;50(9):e13264. doi:10.1111/eci.13264

68. Al-Daghri NM, Al-Attas O, Alokail MS, Alkharfy KM, Draz HM, Agliardi C, Mohammed AK, Guerini FR, Clerici M. Vitamin D receptor gene polymorphisms and HLA DRB1*04 cosegregation in Saudi type 2 diabetes patients. J Immunol. 2012;188(3):1325–1332. doi:10.4049/jimmunol.1101954

69. Zhang S, Sousa A, Lin M, Iwano A, Jain R, Ma B, Lee CM, Park JW, Kamle S, Carlson R, et al. Role of Chitinase 3-Like 1 Protein in the Pathogenesis of Hepatic Insulin Resistance in Nonalcoholic Fatty Liver Disease. Cells. 2021;10(2):201. doi:10.3390/cells10020201

70. Yamamoto M, Guo DH, Hernandez CM, Stranahan AM. Endothelial Adora2a Activation Promotes Blood-Brain Barrier Breakdown and Cognitive Impairment in Mice with Diet-Induced Insulin Resistance. J Neurosci. 2019;39(21):4179–4192. doi:10.1523/JNEUROSCI.2506-18.2019

71. Connor A, Baumgartner RN, Kerber RA, O’Brien E, Rai SN, Wolff RK, Slattery ML, Giuliano AR, Risendal BC, Byers TE, et al. ADRB2 G-G haplotype associated with breast cancer risk among Hispanic and non-Hispanic white women: interaction with type 2 diabetes and obesity. Cancer Causes Control. 2012;23(10):1653–1663. doi:10.1007/s10552-012-0043-6

72. Cai R, Han J, Sun J, Huang R, Tian S, Shen Y, Dong X, Xia W, Wang S. Plasma Clusterin and the CLU Gene rs11136000 Variant Are Associated with Mild Cognitive Impairment in Type 2 Diabetic Patients. Front Aging Neurosci. 2016;8:179. doi:10.3389/fnagi.2016.00179

73. Woods TC, Satou R, Miyata K, Katsurada A, Dugas CM, Klingenberg NC, Fonseca VA, Navar LG. Canagliflozin Prevents Intrarenal Angiotensinogen Augmentation and Mitigates Kidney Injury and Hypertension in Mouse Model of Type 2 Diabetes Mellitus. Am J Nephrol. 2019;49(4):331–342. doi:10.1159/000499597

74. Nakamura K, Yamagishi S, Adachi H, Matsui T, Kurita-Nakamura Y, Takeuchi M, Inoue H, Imaizumi T. Serum levels of soluble form of receptor for advanced glycation end products (sRAGE) are positively associated with circulating AGEs and soluble form of VCAM-1 in patients with type 2 diabetes. Microvasc Res. 2008;76(1):52–56. doi:10.1016/j.mvr.2007.09.004

75. Bajaj M, Suraamornkul S, Hardies LJ, Glass L, Musi N, DeFronzo RA. Effects of peroxisome proliferator-activated receptor (PPAR)-alpha and PPAR-gamma agonists on glucose and lipid metabolism in patients with type 2 diabetes mellitus. Diabetologia. 2007;50(8):1723–1731. doi:10.1007/s00125-007-0698-9

76. Chan GC, Divers J, Russell GB, Langefeld CD, Wagenknecht LE, Hsu FC, Xu J, Smith SC, Palmer ND, Hicks PJ, et al. FGF23 Concentration and APOL1 Genotype Are Novel Predictors of Mortality in African Americans With Type 2 Diabetes. Diabetes Care. 2018;41(1):178–186. doi:10.2337/dc17-0820

77. Caracciolo V, Young J, Gonzales D, Ni Y, Flowers SJ, Summer R, Waldman SA, Kim JK, Jung DY, Noh HL, et al. Myeloid-specific deletion of Zfp36 protects against insulin resistance and fatty liver in diet-induced obese mice. Am J Physiol Endocrinol Metab. 2018;315(4):E676–E693. doi:10.1152/ajpendo.00224.2017

78. Wu YS, Li ZM, Chen YT, Dai SJ, Zhou XJ, Yang YX, Lou JS, Ji LT, Bao YT, Xuan L, et al. Berberine Improves Inflammatory Responses of Diabetes Mellitus in Zucker Diabetic Fatty Rats and Insulin-Resistant HepG2 Cells through the PPM1B Pathway. J Immunol Res. 2020;2020:2141508. doi:10.1155/2020/2141508

79. Opazo-Ríos L, Sanchez Matus Y, Rodrigues-Díez RR, Carpio D, Droguett A, Egido J, Gomez-Guerrero C, Mezzano S.Anti-inflammatory, antioxidant and renoprotective effects of SOCS1 mimetic peptide in the BTBR ob/ob mouse model of type 2 diabetes. BMJ Open Diabetes Res Care. 2020;8(1):e001242. doi:10.1136/bmjdrc-2020-001242

80. Hong CT, Chen KY, Wang W, Chiu JY, Wu D, Chao TY, Hu CJ, Chau KD, Bamodu OA. Insulin Resistance Promotes Parkinson’s Disease through Aberrant Expression of α-synuclein, Mitochondrial Dysfunction, and Deregulation of the Polo-Like Kinase 2 Signaling. Cells. 2020;9(3):740. doi:10.3390/cells9030740

81. Wen W, Sun H, Yang Y, Jia Y, Fang F, Qin Y, Zhang M, Wei Y. Usefulness of Cathepsin S to Predict Risk for Obstructive Sleep Apnea among Patients with Type 2 Diabetes. Dis Markers. 2020;2020:8819134. doi:10.1155/2020/8819134

82. Wu X, Yu T, Ji N, Huang Y, Gao L, Shi W, Yan Y, Li H, Ma L, Wu K, et al. IL6R inhibits viability and apoptosis of pancreatic beta-cells in type 2 diabetes mellitus via regulation by miR-22 of the JAK/STAT signaling pathway. Diabetes Metab Syndr Obes. 2019;12:1645–1657. doi:10.2147/DMSO.S211700

83. Wang J, Yang MM, Li YB, Liu GD, Teng Y, Liu XM. Association of CFH and CFB gene polymorphisms with retinopathy in type 2 diabetic patients. Mediators Inflamm. 2013;2013:748435. doi:10.1155/2013/748435

84. Soraia Aguiar de Melo Dias R, Carlos Mourão Pinho R, Almeida FR, Alfredo Farias Bandeira F, Celerino da Silva R, Crovella S, de Carvalho Farias Vajgel B, Cimões R. Evaluation of DEFB1 polymorphisms in individuals with chronic periodontitis and diabetes mellitus type 2 in a population of northeastern Brazil. Spec Care Dentist. 2018;38(4):227–233. doi:10.1111/scd.12296

85. van Diepen JA, Jansen PA, Ballak DB, Hijmans A, Rutjes FP, Tack CJ, Netea MG, Schalkwijk J, Stienstra R. Genetic and pharmacological inhibition of vanin-1 activity in animal models of type 2 diabetes. Sci Rep. 2016;6:21906. Published 2016 Mar 2. doi:10.1038/srep21906

86. Jiang S, Shen D, Jia WJ, Han X, Shen N, Tao W, Gao X, Xue B, Li CJ. GGPPS-mediated Rab27A geranylgeranylation regulates β cell dysfunction during type 2 diabetes development by affecting insulin granule docked pool formation. J Pathol. 2016;238(1):109–119. doi:10.1002/path.4652

87. Lee SA, Kim YR, Yang EJ, Kwon EJ, Kim SH, Kang SH, Park DB, Oh BC, Kim J, Heo ST, et al. CD26/DPP4 levels in peripheral blood and T cells in patients with type 2 diabetes mellitus. J Clin Endocrinol Metab. 2013;98(6):2553–2561. doi:10.1210/jc.2012-4288

88. Zhao K, Ding W, Zhang Y, Ma K, Wang D, Hu C, Liu J, Zhang X. Variants in the RARRES2 gene are associated with serum chemerin and increase the risk of diabetic kidney disease in type 2 diabetes. Int J Biol Macromol. 2020;165(Pt A):1574–1580. doi:10.1016/j.ijbiomac.2020.10.030

89. Vincent JA, Mohr S. Inhibition of caspase-1/interleukin-1beta signaling prevents degeneration of retinal capillaries in diabetes and galactosemia. Diabetes. 2007;56(1):224–230. doi:10.2337/db06-0427

90. Wang W, Ye S, Qian L, Xing Y, Ren A, Chen C, Li S, Xu J, Liu Q, Dong L, et al. Elevated serum lipocalin 2 levels are associated with indexes of both glucose and bone metabolism in type 2 diabetes mellitus. Endokrynol Pol. 2018;69(3):276–282. doi:10.5603/EP.a2018.0030

91. Wu Y, Quan Y, Liu Y, Liu K, Li H, Jiang Z, Zhang T, Lei H, Radek KA, Li D, et al. Hyperglycaemia inhibits REG3A expression to exacerbate TLR3-mediated skin inflammation in diabetes. Nat Commun. 2016;7:13393. doi:10.1038/ncomms13393

92. Chan PC, Wu TN, Chen YC, Lu CH, Wabitsch M, Tian YF, Hsieh PS. Targetted inhibition of CD74 attenuates adipose COX-2-MIF-mediated M1 macrophage polarization and retards obesity-related adipose tissue inflammation and insulin resistance. Clin Sci (Lond). 2018;132(14):1581–1596. doi:10.1042/CS20180041

93. Chang TJ, Chiu YF, Sheu WH, Shih KC, Hwu CM, Quertermous T, Jou YS, Kuo SS, et al. Genetic polymorphisms of PCSK2 are associated with glucose homeostasis and progression to type 2 diabetes in a Chinese population. Sci Rep. 2015;5:14380. doi:10.1038/srep14380

94. Herold Z, Herold M, Rosta K, Doleschall M, Somogyi A. Lower serum chromogranin B level is associated with type 1 diabetes and with type 2 diabetes patients with intensive conservative insulin treatment. Diabetol Metab Syndr. 2020;12:61. doi:10.1186/s13098-020-00569-5

95. Kwanbunjan K, Panprathip P, Phosat C, Chumpathat N, Wechjakwen N, Puduang S, Auyyuenyong R, Henkel I, Schweigert FJ. Association of retinol binding protein 4 and transthyretin with triglyceride levels and insulin resistance in rural thais with high type 2 diabetes risk. BMC Endocr Disord. 2018;18(1):26. doi:10.1186/s12902-018-0254-2

96. Liu JJ, Pek SLT, Wang J, Liu S, Ang K, Shao YM, Tang JI, Gurung RL, Tavintharan S, Tang WE, et al. Association of Plasma Leucine-Rich α Glycoprotein 1, a Modulator of Transforming Growth Factor-β Signaling Pathway, With Incident Heart Failure in Individuals With Type 2 Diabetes. Diabetes Care. 2021;44(2):571-577. doi:10.2337/dc20-2065

97. Hu F, Lou Y, Shi J, Cao L, Wang C, Ma J, Peng X, Xu S, Chen H, Zhao D et al. Baseline serum albumin and its dynamic change is associated with type 2 diabetes risk: A large cohort study in China. Diabetes Metab Res Rev. 2020;36(5):e3296. doi:10.1002/dmrr.3296

98. Makrilakis K. The Role of DPP-4 Inhibitors in the Treatment Algorithm of Type 2 Diabetes Mellitus: When to Select, What to Expect. Int J Environ Res Public Health. 2019;16(15):2720. doi:10.3390/ijerph16152720

99. Castro A, Lázaro I, Selva DM, Céspedes E, Girona J, NúriaPlana, Guardiola M, Cabré A, Simó R, Masana L. APOH is increased in the plasma and liver of type 2 diabetic patients with metabolic syndrome. Atherosclerosis. 2010;209(1):201–205. doi:10.1016/j.atherosclerosis.2009.09.072

100. Ding L, Houben T, Oligschlaeger Y, Bitorina AV, Verwer BJ, Tushuizen ME, Shiri-Sverdlov R. Plasma Cathepsin D Activity Rather Than Levels Correlates With Metabolic Parameters of Type 2 Diabetes in Male Individuals. Front Endocrinol (Lausanne). 2020;11:575070. doi:10.3389/fendo.2020.575070

101. Li L, Gao K, Zhao J, Feng T, Yin L, Wang J, Wang C, Li C, Wang Y, Wang Q, et al. Glucagon gene polymorphism modifies the effects of smoking and physical activity on risk of type 2 diabetes mellitus in Han Chinese. Gene. 2014;534(2):352–355. doi:10.1016/j.gene.2013.09.121

102. Travers ME, Mackay DJ, Dekker Nitert M, Morris AP, Lindgren CM, Berry A, Johnson PR, Hanley N, Groop LC, McCarthy MI, et al. Insights into the molecular mechanism for type 2 diabetes susceptibility at the KCNQ1 locus from temporal changes in imprinting status in human islets. Diabetes. 2013;62(3):987–992. doi:10.2337/db12-0819

103. Ming Y, Yin Y, Sun Z. Interaction of Nuclear Receptor Subfamily 4 Group A Member 1 (Nr4a1) and Liver Linase B1 (LKB1) Mitigates Type 2 Diabetes Mellitus by Activating Monophosphate-Activated Protein Kinase (AMPK)/Sirtuin 1 (SIRT1) Axis and Inhibiting Nuclear Factor-kappa B (NF-κB) Activation. Med Sci Monit. 2020;26:e920278.. doi:10.12659/MSM.920278

104. Gemmink A, Daemen S, Brouwers B, Huntjens PR, Schaart G, Moonen-Kornips E, Jörgensen J, Hoeks J, Schrauwen P, Hesselink MKC. Dissociation of intramyocellular lipid storage and insulin resistance in trained athletes and type 2 diabetes patients; involvement of perilipin 5?. J Physiol. 2018;596(5):857–868. doi:10.1113/JP275182

105. Li GY, Li ZB, Li F, Dong LP, Tang L, Xiang J, Li JM, Bao MH. Meta-Analysis on the Association of ALDH2 Polymorphisms and Type 2 Diabetic Mellitus, Diabetic Retinopathy. Int J Environ Res Public Health. 2017;14(2):165. doi:10.3390/ijerph14020165

106. Siebert J, Reiwer-Gostomska M, Mysliwska J, Marek N, Raczynska K, Glasner L. Glycemic control influences serum angiogenin concentrations in patients with type 2 diabetes. Diabetes Care. 2010;33(8):1829–1830. doi:10.2337/dc10-0130

107. Belalcazar LM, Papandonatos GD, McCaffery JM, Peter I, Pajewski NM, Erar B, Allred ND, Balasubramanyam A, Bowden DW, Brautbar A, et al. A common variant in the CLDN7/ELP5 locus predicts adiponectin change with lifestyle intervention and improved fitness in obese individuals with diabetes. Physiol Genomics. 2015;47(6):215–224. doi:10.1152/physiolgenomics.00109.2014

108. Erdenebaatar C, Yamaguchi M, Monsur M, Saito F, Honda R, Tashiro H, Ohba T, Iyama KI, Katabuchi H. Serum Prolactin Contributes to Enhancing Prolactin Receptor and pJAK2 in Type I Endometrial Cancer Cells in Young Women Without Insulin Resistance. Int J Gynecol Pathol. 2019;38(4):318–325. doi:10.1097/PGP.0000000000000527

109. Işikli A, Kubat-Üzüm A, Satman İ, Matur Z, Öge AE, Küçükali Cİ, Tüzün E, Erden S, Özkök E. A SOD2 Polymorphism is Associated with Abnormal Quantitative Sensory Testing in Type 2 Diabetic Patients. Noro Psikiyatr Ars. 2018;55(3):276–279. doi:10.29399/npa.23027

110. Mtiraoui N, Turki A, Nemr R, Echtay A, Izzidi I, Al-Zaben GS, Irani-Hakime N, Keleshian SH, Mahjoub T, Almawi WY. Contribution of common variants of ENPP1, IGF2BP2, KCNJ11, MLXIPL, PPARγ SLC30A8 and TCF7L2 to the risk of type 2 diabetes in Lebanese and Tunisian Arabs. Diabetes Metab. 2012;38(5):444–449. doi:10.1016/j.diabet.2012.05.002

111. Ding L, Houben T, Oligschlaeger Y, Bitorina AV, Verwer BJ, Tushuizen ME, Shiri-Sverdlov R. Plasma Cathepsin D. Plasma Cathepsin D Activity Rather Than Levels Correlates With Metabolic Parameters of Type 2 Diabetes in Male Individuals. Front Endocrinol (Lausanne). 2020;11:575070. doi:10.3389/fendo.2020.575070

112. Završnik M, Kariž S, Makuc J, Šeruga M, Cilenšek I, Petrovič D. PECAM-1 Leu125Val (rs688) Polymorphism and Diabetic Nephropathy in Caucasians with Type 2 Diabetes Mellitus. Anal Cell Pathol (Amst). 2016;2016:3152967. doi:10.1155/2016/3152967

113. Niraula A, Thapa S, Kunwar S, Lamsal M, Baral N, Maskey R. Adenosine deaminase activity in type 2 diabetes mellitus: does it have any role?. BMC Endocr Disord. 2018;18(1):58. doi:10.1186/s12902-018-0284-9

114. Sun G, Liu J, Xia G, Zhang L, Li Y, Zhou Z, Lv Y, Wei S, Liu J, Quan J. Reduced serum milk fat globule-epidermal growth factor 8 (MFG-E8) concentrations are associated with an increased risk of microvascular complications in patients with type 2 diabetes. Clin Chim Acta. 2017;466:201–206. doi:10.1016/j.cca.2017.01.013

115. Lin G, Wan X, Liu D, Wen Y, Yang C, Zhao C. COL1A1 as a potential new biomarker and therapeutic target for type 2 diabetes. Pharmacol Res. 2021;105436. doi:10.1016/j.phrs.2021.105436

116. Gaikwad AB, Gupta J, Tikoo K. Epigenetic changes and alteration of Fbn1 and Col3A1 gene expression under hyperglycaemic and hyperinsulinaemic conditions. Biochem J. 2010;432(2):333–341. doi:10.1042/BJ20100414

117. Sasso FC, Marfella R, Pagano A, Porta G, Signoriello G, Lascar N, Minutolo R, Carbonara O, Persico M, Piscione F, et al. Lack of effect of aspirin in primary CV prevention in type 2 diabetic patients with nephropathy: results from 8 years follow-up of NID-2 study. Acta Diabetol. 2015;52(2):239–247. doi:10.1007/s00592-014-0623-x

118. Fawad Ali Shah S, Iqbal T, Naveed N, Akram S, Arshad Rafiq M, Hussain S. ARG1 single nucleotide polymorphisms rs2781666 and rs2781665 confer risk of Type 2 diabetes mellitus. EXCLI J. 2018;17:847–855. doi:10.17179/excli2018-1178

119. Lee M, Park HS, Choi MY, Kim HZ, Moon SJ, Ha JY, Choi A, Park YW, Park JS, Shin EC, et al. Significance of Soluble CD93 in Type 2 Diabetes as a Biomarker for Diabetic Nephropathy: Integrated Results from Human and Rodent Studies. J Clin Med. 2020;9(5):1394. doi:10.3390/jcm9051394

120. Mercader JM, Liao RG, Bell AD, Dymek Z, Estrada K, Tukiainen T, Huerta-Chagoya A, Moreno-Macías H, Jablonski KA, Hanson RL, et al. A Loss-of-Function Splice Acceptor Variant in IGF2 Is Protective for Type 2 Diabetes. Diabetes. 2017;66(11):2903–2914. doi:10.2337/db17-018

121. Hivert MF, Sun Q, Shrader P, Mantzoros CS, Meigs JB, Hu FB. Circulating IL-18 and the risk of type 2 diabetes in women. Diabetologia. 2009;52(10):2101–2108. doi:10.1007/s00125-009-1455-z

122. Perry JR, Voight BF, Yengo L, Amin N, Dupuis J, Ganser M, Grallert H, Navarro P, Li M, Qi L, et al. Yengo L, et al. Stratifying type 2 diabetes cases by BMI identifies genetic risk variants in LAMA1 and enrichment for risk variants in lean compared to obese cases. PLoS Genet. 2012;8(5):e1002741. doi:10.1371/journal.pgen.1002741

123. Simeonovic CJ, Ziolkowski AF, Wu Z, Choong FJ, Freeman C, Parish CR. Heparanase and autoimmune diabetes. Front Immunol. 2013;4:471. doi:10.3389/fimmu.2013.00471

124. Youn JY, Zhou J, Cai H. Bone Morphogenic Protein 4 Mediates NOX1-Dependent eNOS Uncoupling, Endothelial Dysfunction, and COX2 Induction in Type 2 Diabetes Mellitus. Mol Endocrinol. 2015;29(8):1123-1133. doi:10.1210/ME.2014-1313

125. Vidaković M, Grdović N, Dinić S, Mihailović M, Uskoković A, Arambašić Jovanović J. The Importance of the CXCL12/CXCR4 Axis in Therapeutic Approaches to Diabetes Mellitus Attenuation. Front Immunol. 2015;6:403. doi:10.3389/fimmu.2015.00403

126. Merlo S, Starčević JN, Mankoč S, Šantl Letonja M, Cokan Vujkovac A, Zorc M, Petrovič D. Vascular Endothelial Growth Factor Gene Polymorphism (rs2010963) and and Its Receptor, Kinase Insert Domain-Containing Receptor Gene Polymorphism (rs2071559), and Markers of Carotid Atherosclerosis in Patients with Type 2 Diabetes Mellitus. J Diabetes Res. 2016;2016:1482194. doi:10.1155/2016/1482194

127. Kacso IM, Potra AR, Rusu A, Moldovan D, Rusu CC, Kacso G, Hancu ND, Muresan A, Bondor CI. su A, et al. Relationship of endothelial cell selective adhesion molecule to markers of oxidative stress in type 2 diabetes. Scand J Clin Lab Invest. 2014;74(2):170-176. doi:10.3109/00365513.2013.869700

128. Bai J, Xia M, Xue Y, Ma F, Cui A, Sun Y, Han Y, Xu X, Zhang F, Hu Z, et al. Thrombospondin 1 improves hepatic steatosis in diet-induced insulin-resistant mice and is associated with hepatic fat content in humans. EBioMedicine. 2020;57:102849. doi:10.1016/j.ebiom.2020.102849

129. Yamashita A, Nishihira K, Matsuura Y, Ito T, Kawahara K, Hatakeyama K, Hashiguchi T, Maruyama I, Yagi H, Matsumoto M, et al. Paucity of CD34-positive cells and increased expression of high-mobility group box 1 in coronary thrombus with type 2 diabetes mellitus. Atherosclerosis. 2012;224(2):511–514. doi:10.1016/j.atherosclerosis.2012.07.027

130. Fan Q, Li H, Qin Y, Li L, Chen L, Zhang L, Lv Y, Liang D, Liang Y, Long T, et al. Association of SERPINE1 rs6092 with type 2 diabetes and related metabolic traits in a Chinese population. Gene. 2018;661:176–181. doi:10.1016/j.gene.2018.04.011

131. Salpea KD, Gable DR, Cooper JA, Stephens JW, Hurel SJ, Ireland HA, Feher MD, Godsland IF, Humphries SE. The effect of WNT5B IVS3C>G on the susceptibility to type 2 diabetes in UK Caucasian subjects. Nutr Metab Cardiovasc Dis. 2009;19(2):140–145. doi:10.1016/j.numecd.2008.02.009

132. Sardana M, Vasim I, Varakantam S, Kewan U, Tariq A, Koppula MR, Syed AA, Beraun M, Drummen NE, Vermeer C, et al. Inactive Matrix Gla-Protein and Arterial Stiffness in Type 2 Diabetes Mellitus. Am J Hypertens. 2017;30(2):196–201. doi:10.1093/ajh/hpw146

133. Lee JE, Choi YK, Seo HA, Jeon JH, Jeong JY, Moon SS, Kim JG, Kim BW, Kim SW, Min Yoo, et al. Impact of ENPP1 and MMP3 gene polymorphisms on aortic calcification in patients with type 2 diabetes in a Korean population. Diabetes Res Clin Pract. 2010;88(1):87–96. doi:10.1016/j.diabres.2010.01.002

134. Kaminari A, Tsilibary EC, Tzinia A. A New Perspective in Utilizing MMP-9 as a Therapeutic Target for Alzheimer’s Disease and Type 2 Diabetes Mellitus. J Alzheimers Dis. 2018;64(1):1–16. doi:10.3233/JAD-180035

135. Arcidiacono B, Chiefari E, Laria AE, Messineo S, Bilotta FL, Britti D, Foti DP, Foryst-Ludwig A, Kintscher U, Brunetti A. Expression of matrix metalloproteinase-11 is increased under conditions of insulin resistance. World J Diabetes. 2017;8(9):422–428. doi:10.4239/wjd.v8.i9.422

136. Cheng F, Liu L, Zhang H, Zhu Y, Li X, Li H. Association of IL-16 gene polymorphisms with the risk of developing type 2 diabetes mellitus in the Chinese Han population. Biosci Rep. 2019;39(8):BSR20190821. doi:10.1042/BSR20190821

137. Biscetti F, Porreca CF, Bertucci F, Straface G, Santoliquido A, Tondi P, Angelini F, Pitocco D, Santoro L, Gasbarrini A, et al. TNFRSF11B gene polymorphisms increased risk of peripheral arterial occlusive disease and critical limb ischemia in patients with type 2 diabetes. Acta Diabetol. 2014;51(6):1025–1032. doi:10.1007/s00592-014-0664-1

138. Jiang C, Xia W, Wu T, Pan C, Shan H, Wang F, Zhou Z, Yu X. Inhibition of microRNA-222 up-regulates TIMP3 to promotes osteogenic differentiation of MSCs from fracture rats with type 2 diabetes mellitus. J Cell Mol Med. 2020;24(1):686–694. doi:10.1111/jcmm.14777

139. Walder K, McMillan J, Lapsys N, Kriketos A, Trevaskis J, Civitarese A, Southon A, Zimmet P, Collier G. Calpain 3 gene expression in skeletal muscle is associated with body fat content and measures of insulin resistance. Int J Obes Relat Metab Disord. 2002;26(4):442–449. doi:10.1038/sj.ijo.0801946

140. Howson JM, Walker NM, Clayton D, Todd JA; Type 1 Diabetes Genetics Consortium. Confirmation of HLA class II independent type 1 diabetes associations in the major histocompatibility complex including HLA-B and HLA-A. Diabetes Obes Metab. 2009;11 Suppl 1(Suppl 1):31-45. doi:10.1111/j.1463-1326.2008.01001.x

141. Schloss J, Ali R, Racine JJ, Chapman HD, Serreze DV, DiLorenzo TP. HLA-B*39:06 Efficiently Mediates Type 1 Diabetes in a Mouse Model Incorporating Reduced Thymic Insulin Expression. J Immunol. 2018;200(10):3353–3363. doi:10.4049/jimmunol.1701652

142. Zhi D, Sun C, Sedimbi SK, Luo F, Shen S, Sanjeevi CB. Killer cell immunoglobulin-like receptor along with HLA-C ligand genes are associated with type 1 diabetes in Chinese Han population. Diabetes Metab Res Rev. 2011;27(8):872–877. doi:10.1002/dmrr.1264

143. Varney MD, Valdes AM, Carlson JA, Noble JA, Tait BD, Bonella P, Lavant E, Fear AL, Louey A, Moonsamy P, et al. HLA DPA1, DPB1 alleles and haplotypes contribute to the risk associated with type 1 diabetes: analysis of the type 1 diabetes genetics consortium families. Diabetes. 2010;59(8):2055-2062. doi:10.2337/db09-0680

144. Jiang H, Canfield SM, Gallagher MP, Jiang HH, Jiang Y, Zheng Z, Chess L. HLA-E-restricted regulatory CD8(+) T cells are involved in development and control of human autoimmune type 1 diabetes. J Clin Invest. 2010;120(10):3641–3650. doi:10.1172/JCI4352

145. Wang D, Pan G. The Association between rs2292239 Polymorphism in ERBB3 Gene and Type 1 Diabetes: A Meta-Analysis. Biomed Res Int. 2019;2019:7689642. doi:10.1155/2019/7689642

146. Blindbæk SL, Schlosser A, Green A, Holmskov U, Sorensen GL, Grauslund J. Association between microfibrillar-associated protein 4 (MFAP4) and micro-and macrovascular complications in long-term type 1 diabetes mellitus. Acta Diabetol. 2017;54(4):367–372. doi:10.1007/s00592-016-0953-y

147. Cetkovic-Cvrlje M, Dragt AL, Vassilev A, Liu XP, Uckun FM. Targeting JAK3 with JANEX-1 for prevention of autoimmune type 1 diabetes in NOD mice. Clin Immunol. 2003;106(3):213–225. doi:10.1016/s1521-6616(02)00049-9

148. Li J, Jian L, Huang J, Xiong N, Wang T. The rs3129882/rs4248166 in HLA-DRA and rs34372695 in SYT11 are not associated with sporadic Parkinson’s disease in Central Chinese population. Int J Neurosci. 2020;1–7. doi:10.1080/00207454.2020.1753728

149. Halbgebauer S, Nagl M, Klafki H, Haußmann U, Steinacker P, Oeckl P, Kassubek J, Pinkhardt E, Ludolph AC, Soininen H, et al. Modified serpinA1 as risk marker for Parkinson’s disease dementia: Analysis of baseline data. Sci Rep. 2016;6:26145. doi:10.1038/srep26145

150. Ichu TA, Reed A, Ogasawara D, Ulanovskaya O, Roberts A, Aguirre CA, Bar-Peled L, Gao J, Germain J, Barbas S, et al. ABHD12 and LPCAT3 Interplay Regulates a Lyso-phosphatidylserine-C20:4 Phosphatidylserine Lipid Network Implicated in Neurological Disease. Biochemistry. 2020;59(19):1793–1799. doi:10.1021/acs.biochem.0c00292

151. Bloch PJ, Weller AE, Doyle GA, Ferraro TN, Berrettini WH, Hodge R, Lohoff FW. Association analysis between polymorphisms in the myo-inositol monophosphatase 2 (IMPA2) gene and bipolar disorder. Prog Neuropsychopharmacol Biol Psychiatry. 2010;34(8):1515–1519. doi:10.1016/j.pnpbp.2010.08.015

152. Yoo HS, Lee JS, Chung SJ, Ye BS, Sohn YH, Lee SJ, Lee PH. Changes in plasma arylsulfatase A level as a compensatory biomarker of early Parkinson’s disease. Sci Rep. 2020;10(1):5567. doi:10.1038/s41598-020-62536-4

153. Cappuccio G, Attanasio S, Alagia M, Mutarelli M, Borzone R, Karali M, Genesio R, Mormile A, Nitsch L, Imperati F, et al. Microdeletion of pseudogene chr14.232.a affects LRFN5 expression in cells of a patient with autism spectrum disorder. Eur J Hum Genet. 2019;27(9):1475–1480. doi:10.1038/s41431-019-0430-5

154. Han Q, Sun YA, Zong Y, Chen C, Wang HF, Tan L. Common Variants in PLXNA4 and Correlation to CSF-related Phenotypes in Alzheimer’s Disease. Front Neurosci. 2018;12:946. doi:10.3389/fnins.2018.00946

155. Yang CR, Ning L, Zhou FH, Sun Q, Meng HP, Han Z, Liu Y, Huang W, Liu S, Li XH, et al. Downregulation of Adhesion Molecule CHL1 in B Cells but Not T Cells of Patients with Major Depression and in the Brain of Mice with Chronic Stress. Neurotox Res. 2020;38(4):914–928. doi:10.1007/s12640-020-00234-9

156. Stygelbout V, Leroy K, Pouillon V, Ando K, D’Amico E, Jia Y, Luo HR, Duyckaerts C, Erneux C, Schurmans S, et al. on V, et al. Inositol trisphosphate 3-kinase B is increased in human Alzheimer brain and exacerbates mouse Alzheimer pathology. Brain. 2014;137(Pt 2):537–552. doi:10.1093/brain/awt344

157. Alguacil LF, Herradón G. Midkine and Pleiotrophin in the Treatment of Neurodegenerative Diseases and Drug Addiction. Recent Pat CNS Drug Discov. 2015;10(1):28–33. doi:10.2174/1574889810666150326103916

158. Ahmad S, Milan MDC, Hansson O, Demirkan A, Agustin R, Sáez ME, Giagtzoglou N, Cabrera-Socorro A, Bakker MHM, Ramirez A, et al. CDH6 and HAGH protein levels in plasma associate with Alzheimer’s disease in APOE ε4 carriers. Sci Rep. 2020;10(1):8233. doi:10.1038/s41598-020-65038-5

159. Arreguin AJ, Colognato H. Brain Dysfunction in LAMA2-Related Congenital Muscular Dystrophy: Lessons From Human Case Reports and Mouse Models. Front Mol Neurosci. 2020;13:118. doi:10.3389/fnmol.2020.00118

160. Bruno E, Quattrocchi G, Nicoletti A, Le Pira F, Maci T, Mostile G, Andreoli V, Quattrone A, Zappia M. Lack of interaction between LRP1 and A2M polymorphisms for the risk of Alzheimer disease. Neurosci Lett. 2010;482(2):112–116. doi:10.1016/j.neulet.2010.07.012

161. Zidi I, Kharrat N, Abdelhedi R, Hassine AB, Laaribi AB, Yahia HB, Abdelmoula NB, Abid L, Rebai A, Rizzo R. Nonclassical human leukocyte antigen (HLA-G, HLA-E, and HLA-F) in coronary artery disease. Hum Immunol. 2016;77(4):325–329. doi:10.1016/j.humimm.2016.01.008

162. Annichino-Bizzacchi JM, Saad ST, Arruda VR, Ramires JA, Siqueira LH, Chiaparini LC, Mansur AP. C282Y mutation in the HLA-H gene is not a risk factor for patients with myocardial infarction. J Cardiovasc Risk. 2000;7(1):37–40. doi:10.1177/204748730000700107

163. Jacquemin B, Antoniades C, Nyberg F, Plana E, Müller M, Greven S, Salomaa V, Sunyer J, Bellander T, Chalamandaris AG, et al. Common genetic polymorphisms and haplotypes of fibrinogen alpha, beta, and gamma chains affect fibrinogen levels and the response to proinflammatory stimulation in myocardial infarction survivors: the AIRGENE study. J Am Coll Cardiol. 2008;52(11):941–952. doi:10.1016/j.jacc.2008.06.016

164. Yang Z, Zhang Q, Yu H, Du H, Li L, He Y, Zhu S, Li C, Zhang S, Luo B, et al. Genetic association study of a novel indel polymorphism in HSPA1B with the risk of sudden cardiac death in the Chinese populations. Forensic Sci Int. 2021;318:110637. doi:10.1016/j.forsciint.2020.110637

165. Yarnazari A, Hassanpour P, Hosseini-Fard SR, Amirfarhangi A, Najafi M. The sdLDL Reduces MRC1 Expression Level and Secretion of Histamin e in Differentiated M2-macrophages from Patients with Coronary Artery Stenosis. Cardiovasc Hematol Disord Drug Targets. 2017;17(1):28–32. doi:10.2174/1871529X17666170106095554

166. Harrison SC, Cooper JA, Li K, Talmud PJ, Sofat R, Stephens JW, Hamsten A; HIFMECH Consortium, Sanders J, Montgomery H, Neil A; Simon Broome Research Consortium, Humphries SE. Association of a sequence variant in DAB2IP with coronary heart disease. Eur Heart J. 2012;33(7):881–888. doi:10.1093/eurheartj/ehr075

167. Delaney JT, Muhammad R, Blair MA, Kor K, Fish FA, Roden DM, Darbar D. A KCNJ8 mutation associated with early repolarization and atrial fibrillation. Europace. 2012;14(10):1428–1432. doi:10.1093/europace/eus150

168. Gittleman HR, Merkulova A, Alhalabi O, Stavrou EX, Veigl ML, Barnholtz-Sloan JS, Schmaier AH. A Cross-sectional Study of KLKB1 and PRCP Polymorphisms in Patient Samples with Cardiovascular Disease. Front Med (Lausanne). 2016;3:17. doi:10.3389/fmed.2016.00017

169. Guo LY, Yang F, Peng LJ, Li YB, Wang AP. CXCL2, a new critical factor and therapeutic target for cardiovascular diseases. Clin Exp Hypertens. 2020;42(5):428–437. doi:10.1080/10641963.2019.1693585

170. Li X, Zhao D, Guo Z, Li T, Qili M, Xu B, Qian M, Liang H, E X, Chege Gitau S, Wang L, et al. Overexpression of SerpinE2/protease nexin-1 Contribute to Pathological Cardiac Fibrosis via increasing Collagen Deposition. Sci Rep. 2016;6:37635. doi:10.1038/srep37635

171. Latella MC, Di Castelnuovo A, de Lorgeril M, Arnout J, Cappuccio FP, Krogh V, Siani A, van Dongen M, Donati MB, de Gaetano G, et al. Genetic variation of alcohol dehydrogenase type 1C (ADH1C), alcohol consumption, and metabolic cardiovascular risk factors: results from the IMMIDIET study. Atherosclerosis. 2009;207(1):284–290. doi:10.1016/j.atherosclerosis.2009.04.022

172. Liu H, Luo D, Qiu Y, Huang Y, Chen C, Song X, Gao L, Zhou Y. The Effect of AMBP SNPs, Their Haplotypes, and Gene-Environment Interactions on the Risk of Atherothrombotic Stroke Among the Chinese Population. Genet Test Mol Biomarkers. 2019;23(7):487–494. doi:10.1089/gtmb.2018.0248

173. Kardys I, van Tiel CM, de Vries CJ, Pannekoek H, Uitterlinden AG, Hofman A, Witteman JC, de Maat MP. Haplotypes of the NR4A2/NURR1 gene and cardiovascular disease: the Rotterdam Study. Hum Mutat. 2009;30(3):417–423. doi:10.1002/humu.20902

174. Li W, Yue H. Thymidine phosphorylase: A potential new target for treating cardiovascular disease. Trends Cardiovasc Med. 2018;28(3):157–171. doi:10.1016/j.tcm.2017.10.003

175. Xu W, Barrientos T, Mao L, Rockman HA, Sauve AA, Andrews NC. Lethal Cardiomyopathy in Mice Lacking Transferrin Receptor in the Heart. Cell Rep. 2015;13(3):533–545. doi:10.1016/j.celrep.2015.09.023

176. Persson M, Engström G, Björkbacka H, Hedblad B. Soluble urokinase plasminogen activator receptor in plasma is associated with incidence of CVD. Results from the Malmö Diet and Cancer Study. Atherosclerosis. 2012;220(2):502–505. doi:10.1016/j.atherosclerosis.2011.10.039

177. Grossman TR, Gamliel A, Wessells RJ, Taghli-Lamallem O, Jepsen K, Ocorr K, Korenberg JR, Peterson KL, Rosenfeld MG, Bodmer R, et al. Over-expression of DSCAM and COL6A2 cooperatively generates congenital heart defects. PLoS Genet. 2011;7(11):e1002344. doi:10.1371/journal.pgen.1002344

178. Connelly JJ, Cherepanova OA, Doss JF, Karaoli T, Lillard TS, Markunas CA, Nelson S, Wang T, Ellis PD, Langford CF, et al. Epigenetic regulation of COL15A1 in smooth muscle cell replicative aging and atherosclerosis. Hum Mol Genet. 2013;22(25):5107–5120. doi:10.1093/hmg/ddt365

179. Delfín DA, DeAguero JL, McKown EN. The Extracellular Matrix Protein ABI3BP in Cardiovascular Health and Disease. Front Cardiovasc Med. 2019;6:23. doi:10.3389/fcvm.2019.00023

180. Hu YW, Guo FX, Xu YJ, Li P, Lu ZF, McVey DG, Zheng L, Wang Q, Ye JH, Kang CM, et al. Long noncoding RNA NEXN-AS1 mitigates atherosclerosis by regulating the actin-binding protein NEXN. J Clin Invest. 2019;129(3):1115–1128. doi:10.1172/JCI98230

181. Deng S, Zhou X, Ge Z, Song Y, Wang H, Liu X, Zhang D. Exosomes from adipose-derived mesenchymal stem cells ameliorate cardiac damage after myocardial infarction by activating S1P/SK1/S1PR1 signaling and promoting macrophage M2 polarization. Int J Biochem Cell Biol. 2019;114:105564. doi:10.1016/j.biocel.2019.105564

182. Li Y, Song D, Mao L, Abraham DM, Bursac N. Lack of Thy1 defines a pathogenic fraction of cardiac fibroblasts in heart failure. Biomaterials. 2020;236:119824. doi:10.1016/j.biomaterials.2020.119824

183. Raza ST, Abbas S, Eba A, Karim F, Wani IA, Rizvi S, Zaidi A, Mahdi F. Association of COL4A1 (rs605143, rs565470) and CD14 (rs2569190) genes polymorphism with coronary artery disease. Mol Cell Biochem. 2018;445(1-2):117–122. doi:10.1007/s11010-017-3257-9

184. Azuaje F, Zhang L, Jeanty C, Puhl SL, Rodius S, Wagner DR. Analysis of a gene co-expression network establishes robust association between Col5a2 and ischemic heart disease. BMC Med Genomics. 2013;6:13. doi:10.1186/1755-8794-6-13

185. Wang X, Chen W, Zhang J, Khan A, Li L, Huang F, Qiu Z, Wang L, Chen X. Critical Role of ADAMTS2 (A Disintegrin and Metalloproteinase With Thrombospondin Motifs 2) in Cardiac Hypertrophy Induced by Pressure Overload. Hypertension. 2017;69(6):1060–1069. doi:10.1161/HYPERTENSIONAHA.116.08581

186. Hardy SA, Mabotuwana NS, Murtha LA, Coulter B, Sanchez-Bezanilla S, Al-Omary MS, Senanayake T, Loering S, Starkey M, Lee RJ, et al. Novel role of extracellular matrix protein 1 (ECM1) in cardiac aging and myocardial infarction. PLoS One. 2019;14(2):e0212230. doi:10.1371/journal.pone.0212230

187. Chen HX, Yang ZY, Hou HT, Wang J, Wang XL, Yang Q, Liu L, He GW. Novel mutations of TCTN3/LTBP2 with cellular function changes in congenital heart disease associated with polydactyly. J Cell Mol Med. 2020;24(23):13751–13762. doi:10.1111/jcmm.15950

188. El Shamieh S, Stathopoulou MG, Bonnefond A, Ndiaye NC, Lecoeur C, Meyre D, Dadé S, Chedid P, Salami A, Shahabi P, et al. Obesity status modifies the association between rs7556897T>C in the intergenic region SLC19A3-CCL20 and blood pressure in French children. Clin Chem Lab Med. 2020;58(11):1819–1827. doi:10.1515/cclm-2019-0292

189. Mastorakos G, Zapanti E. The hypothalamic-pituitary-adrenal axis in the neuroendocrine regulation of food intake and obesity: the role of corticotropin releasing hormone. Nutr Neurosci. 2004;7(5-6):271–280. doi:10.1080/10284150400020516

190. Yang H, Graham LC, Reagan AM, Grabowska WA, Schott WH, Howell GR. Transcriptome profiling of brain myeloid cells revealed activation of Itgal, Trem1, and Spp1 in western diet-induced obesity. J Neuroinflammation. 2019;16(1):169. doi:10.1186/s12974-019-1527-z

191. Du Y, Li S, Cui CJ, Zhang Y, Yang SH, Li JJ. Leptin decreases the expression of low-density lipoprotein receptor via PCSK9 pathway: linking dyslipidemia with obesity. J Transl Med. 2016;14(1):276. doi:10.1186/s12967-016-1032-4

192. Hams E, Roberts J, Bermingham R, Hogan AE, O’Shea D, O’Neill L, Fallon PG. Role for Retinoic Acid-Related Orphan Receptor Alpha (RORα Expressing Macrophages in Diet-Induced Obesity. Front Immunol. 2020;11:1966. doi:10.3389/fimmu.2020.01966

193. Moreno-Navarrete JM, Latorre J, Lluch A, Ortega FJ, Comas F, Arnoriaga-Rodríguez M, Ricart W, Fernández-Real JM. Lysozyme is a component of the innate immune system linked to obesity associated-chronic low-grade inflammation and altered glucose tolerance. Clin Nutr. 2021;40(3):1420–1429. doi:10.1016/j.clnu.2020.08.036

194. Lee S. The association of genetically controlled CpG methylation (cg158269415) of protein tyrosine phosphatase, receptor type N2 (PTPRN2) with childhood obesity. Sci Rep. 2019;9(1):4855. doi:10.1038/s41598-019-40486-w

195. Soussi H, Reggio S, Alili R, Prado C, Mutel S, Pini M, Rouault C, Clément K, Dugail I. DAPK2 Downregulation Associates With Attenuated Adipocyte Autophagic Clearance in Human Obesity. Diabetes. 2015;64(10):3452–3463. doi:10.2337/db14-1933

196. Inoue K, Maeda N, Mori T, Sekimoto R, Tsushima Y, Matsuda K, Yamaoka M, Suganami T, Nishizawa H, Ogawa Y et al. Possible involvement of Opa-interacting protein 5 in adipose proliferation and obesity. PLoS One. 2014;9(2):e87661. doi:10.1371/journal.pone.0087661

197. Salas-Pérez F, Cuevas-Sierra A, Cuervo M, Goni L, Milagro FI, Martínez JA, Riezu-Boj JI. Differentially methylated regions (DMRs) in PON3 gene between responders and non-responders to a weight loss dietary intervention: a new tool for precision management of obesity. Epigenetics. 2021;1–12. doi:10.1080/15592294.2021.1873629

198. Lee JH, Zhang D, Kwak SE, Shin HE, Song W. Effects of Exercise and a High-Fat, High-Sucrose Restriction Diet on Metabolic Indicators, Nr4a3, and Mitochondria-Associated Protein Expression in the Gastrocnemius Muscles of Mice with Diet-Induced Obesity. J Obes Metab Syndr. 2021;10.7570/jomes20043. doi:10.7570/jomes20043

199. Han CY, Kang I, Harten IA, Gebe JA, Chan CK, Omer M, Alonge KM, den Hartigh LJ, Gomes Kjerulf D, Goodspeed L, et al. Adipocyte-Derived Versican and Macrophage-Derived Biglycan Control Adipose Tissue Inflammation in Obesity. Cell Rep. 2020;31(13):107818. doi:10.1016/j.celrep.2020.107818

200. Buchner DA, Geisinger JM, Glazebrook PA, Morgan MG, Spiezio SH, Kaiyala KJ, Schwartz MW, Sakurai T, Furley AJ, Kunze DL, et al. The juxtaparanodal proteins CNTNAP2 and TAG1 regulate diet-induced obesity. Mamm Genome. 2012;23(7-8):431–442. doi:10.1007/s00335-012-9400-8

201. Yu GI, Song DK, Shin DH. Associations of IL1RAP and IL1RL1 gene polymorphisms with obesity and inflammation mediators. Inflamm Res. 2020;69(2):191–202. doi:10.1007/s00011-019-01307-y

202. Shi Y, Long F. Hedgehog signaling via Gli2 prevents obesity induced by high-fat diet in adult mice. Elife. 2017;6:e31649. doi:10.7554/eLife.31649

203. Jo J, Sull JW, Park EJ, Jee SH. Effects of smoking and obesity on the association between CDH13 (rs3865188) and adiponectin among Korean men: the KARE study. Obesity (Silver Spring). 2012;20(8):1683–1687. doi:10.1038/oby.2011.128

204. Zhang L, Reidy SP, Nicholson TE, Lee HJ, Majdalawieh A, Webber C, Stewart BR, Dolphin P, Ro HS. The role of AEBP1 in sex-specific diet-induced obesity. Mol Med. 2005;11(1-12):39–47. doi:10.2119/2005-00021.Ro

205. Han CY, Kang I, Harten IA, Gebe JA, Chan CK, Omer M, Alonge KM, den Hartigh LJ, Gomes Kjerulf D, Goodspeed L, et al. Adipocyte-Derived Versican and Macrophage-Derived Biglycan Control Adipose Tissue Inflammation in Obesity. Cell Rep. 2020;31(13):107818. doi:10.1016/j.celrep.2020.107818

206. Martínez-Martínez E, Rodríguez C, Galán M, Miana M, Jurado-López R, Bartolomé MV, Luaces M, Islas F, Martínez-González J, López-Andrés N, et al. The lysyl oxidase inhibitor (β-aminopropionitrile) reduces leptin profibrotic effects and ameliorates cardiovascular remodeling in diet-induced obesity in rats. J Mol Cell Cardiol. 2016;92:96–104. doi:10.1016/j.yjmcc.2016.01.012

207. Yu GI, Song DK, Shin DH. Associations of IL1RAP and IL1RL1 gene polymorphisms with obesity and inflammation mediators. Inflamm Res. 2020;69(2):191–202. doi:10.1007/s00011-019-01307-y

208. Wolff G, Taranko AE, Meln I, Weinmann J, Sijmonsma T, Lerch S, Heide D, Billeter AT, Tews D, Krunic D, et al. Diet-dependent function of the extracellular matrix proteoglycan Lumican in obesity and glucose homeostasis. Mol Metab. 2019;19:97–106. doi:10.1016/j.molmet.2018.10.007

209. Liu X, Jiang C, Yang P. Association of single nucleotide polymorphisms in the 5’ upstream region of the C4BPA gene with essential hypertension in a northeastern Han Chinese population. Mol Med Rep. 2017;16(2):1289–1297. doi:10.3892/mmr.2017.6736

210. Zhang Y, Shen J, He X, Zhang K, Wu S, Xiao B, Zhou X, Phillips RS, Gao P, Jeunemaitre X, et al. A rare variant at the KYNU gene is associated with kynureninase activity and essential hypertension in the Han Chinese population. Circ Cardiovasc Genet. 2011;4(6):687–694. doi:10.1161/CIRCGENETICS.110.959064

211. Pei Q, Yang L, Tan HY, Liu SK, Liu Y, Huang L, Li RH, Wan Q, Huang J, Guo CX, et al. Effects of genetic variants in UGT1A1, SLCO1B3, ABCB1, ABCC2, ABCG2, ORM1 on PK/PD of telmisartan in Chinese patients with mild to moderate essential hypertension. Int J Clin Pharmacol Ther. 2017;55(8):659–665. doi:10.5414/CP202744

212. Tang D, Fakiola M, Syn G, Anderson D, Cordell HJ, Scaman ESH, Davis E, Miles SJ, McLeay T, Jamieson SE, et al. Arylsulphatase A Pseudodeficiency (ARSA-PD), hypertension and chronic renal disease in Aboriginal Australians. Sci Rep. 2018;8(1):10912. doi:10.1038/s41598-018-29279-9

213. Durgin BG, Hahn SA, Schmidt HM, Miller MP, Hafeez N, Mathar I, Freitag D, Sandner P, Straub AC. Loss of smooth muscle CYB5R3 amplifies angiotensin II-induced hypertension by increasing sGC heme oxidation. JCI Insight. 2019;4(19):e129183. doi:10.1172/jci.insight.129183

214. Lai CQ, Parnell LD, Troen AM, Shen J, Caouette H, Warodomwichit D, Lee YC, Crott JW, Qiu WQ, Rosenberg IH, et al. MAT1A variants are associated with hypertension, stroke, and markers of DNA damage and are modulated by plasma vitamin B-6 and folate. Am J Clin Nutr. 2010;91(5):1377–1386. doi:10.3945/ajcn.2009.28923

215. Lipphardt M, Dihazi H, Maas JH, Schäfer AK, Amlaz SI, Ratliff BB, Koziolek MJ, Wallbach M. Syndecan-4 as a Marker of Endothelial Dysfunction in Patients with Resistant Hypertension. J Clin Med. 2020;9(9):3051. doi:10.3390/jcm9093051

216. Kho J, Tian X, Wong WT, Bertin T, Jiang MM, Chen S, Jin Z, Shchelochkov OA, Burrage LC, Reddy AK, et al. Argininosuccinate Lyase Deficiency Causes an Endothelial-Dependent Form of Hypertension. Am J Hum Genet. 2018;103(2):276–287. doi:10.1016/j.ajhg.2018.07.008

217. Yang HC, Liang YJ, Chen JW, Chiang KM, Chung CM, Ho HY, Ting CT, Lin TH, Sheu SH, Tsai WC, et al. Identification of IGF1, SLC4A4, WWOX, and SFMBT1 as hypertension susceptibility genes in Han Chinese with a genome-wide gene-based association study. PLoS One. 2012;7(3):e32907. doi:10.1371/journal.pone.0032907

218. Wu T, Zhang BQ, Raelson J, Yao YM, Wu HD, Xu ZX, Marois-Blanchet FC, Tahir MR, Wang Y, Bradley WE, et al. Analysis of the association of EPHB6, EFNB1 and EFNB3 variants with hypertension risks in males with hypogonadism. Sci Rep. 2018;8(1):14497. doi:10.1038/s41598-018-32836-x

219. Keranov S, Dörr O, Jafari L, Liebetrau C, Keller T, Troidl C, Kriechbaum S, Voss S, Richter M, Tello K, et al. SPARCL1 as a biomarker of maladaptive right ventricular remodelling in pulmonary hypertension. Biomarkers. 2020;25(3):290–295. doi:10.1080/1354750X.2020.1745889

220. Oguri M, Kato K, Yokoi K, Watanabe S, Metoki N, Yoshida H, Satoh K, Aoyagi Y, Nishigaki Y, Yoshida H, et al. Association of polymorphisms of THBS2 and HSPA8 with hypertension in Japanese individuals with chronic kidney disease. Mol Med Rep. 2009;2(2):205–211. doi:10.3892/mmr_00000085

221. Wang Y, Hamet P, Thorin E, Tremblay J, Raelson J, Wu Z, Luo H, Jin W, Lavoie JL, Peng J, et al. Reduced blood pressure after smooth muscle EFNB2 deletion and the potential association of EFNB2 mutation with human hypertension risk. Eur J Hum Genet. 2016;24(12):1817–1825. doi:10.1038/ejhg.2016.105

222. Xu T, Shao L, Wang A, Liang R, Lin Y, Wang G, Zhao Y, Hu J, Liu S. CD248 as a novel therapeutic target in pulmonary arterial hypertension. Clin Transl Med. 2020;10(5):e175. doi:10.1002/ctm2.175

223. Cheng Q, Pan J, Zhou ZL, Yin F, Xie HY, Chen PP, Li JY, Zheng PQ, Zhou L, Zhang W, et al. Caspase-11/4 and gasdermin D-mediated pyroptosis contributes to podocyte injury in mouse diabetic nephropathy. Acta Pharmacol Sin. 2020;10.1038/s41401-020-00525-z. doi:10.1038/s41401-020-00525-z

224. Jiao X, Zhang D, Hong Q, Yan L, Han Q, Shao F, Cai G, Chen X, Zhu H. Netrin-1 works with UNC5B to regulate angiogenesis in diabetic kidney disease. Front Med. 2020;14(3):293–304. doi:10.1007/s11684-019-0715-7

225. Uehara G, Suzuki D, Toyoda M, Umezono T, Sakai H. Glomerular expression of platelet-derived growth factor (PDGF)-A, -B chain and PDGF receptor-alpha, -beta in human diabetic nephropathy. Clin Exp Nephrol. 2004;8(1):36–42. doi:10.1007/s10157-003-0265-8

226. Gregg T, Sdao SM, Dhillon RS, Rensvold JW, Lewandowski SL, Pagliarini DJ, Denu JM, Merrins MJ. Obesity-dependent CDK1 signaling stimulates mitochondrial respiration at complex I in pancreatic β-cells. J Biol Chem. 2019;294(12):4656–4666. doi:10.1074/jbc.RA118.006085

227. Fang H, Luo X, Wang Y, Liu N, Fu C, Wang H, Fang Y, Shi W, Zhang Y, Zeng C, et al. Correlation between single nucleotide polymorphisms of the ACTA2 gene and coronary artery stenosis in patients with type 2 diabetes mellitus. Exp Ther Med. 2014;7(4):970–976. doi:10.3892/etm.2014.1510

228. Demirsoy İH, Ertural DY, Balci Ş, Çınkır Ü, Sezer K, Tamer L, Aras N. Profiles of Circulating MiRNAs Following Metformin Treatment in Patients with Type 2 Diabetes. J Med Biochem. 2018;37(4):499–506. doi:10.2478/jomb-2018-0009

229. Hourigan ST, Solly EL, Nankivell VA, Ridiandries A, Weimann BM, Henriquez R, Tepper ER, Zhang JQJ, Tsatralis T, Clayton ZE, et al. The regulation of miRNAs by reconstituted high-density lipoproteins in diabetes-impaired angiogenesis. Sci Rep. 2018;8(1):13596. doi:10.1038/s41598-018-32016-x

230. Safar HA, Chehadeh SEH, Abdel-Wareth L, Haq A, Jelinek HF, ElGhazali G, Anouti FA. Vitamin D receptor gene polymorphisms among Emirati patients with type 2 diabetes mellitus. J Steroid Biochem Mol Biol. 2018;175:119–124. doi:10.1016/j.jsbmb.2017.03.012

231. Zmuda EJ, Qi L, Zhu MX, Mirmira RG, Montminy MR, Hai T. The roles of ATF3, an adaptive-response gene, in high-fat-diet-induced diabetes and pancreatic beta-cell dysfunction. Mol Endocrinol. 2010;24(7):1423–1433. doi:10.1210/me.2009-0463

232. Boese EA, Tollefson MR, Schnieders MJ, Darbro BW, Alward WLM, Fingert JH. Novel Intragenic PAX6 Deletion in a Pedigree with Aniridia, Morbid Obesity, and Diabetes. Curr Eye Res. 2020;45(1):91–96. doi:10.1080/02713683.2019.1649704

233. Suhara T, Baba Y, Shimada BK, Higa JK, Matsui T. The mTOR Signaling Pathway in Myocardial Dysfunction in Type 2 Diabetes Mellitus. Curr Diab Rep. 2017;17(6):38. doi:10.1007/s11892-017-0865-4

234. Cabiati M, Randazzo E, Salvadori C, Peroni D, Federico G, Del Ry S. Circulating microRNAs associated with C-type natriuretic peptide in childhood obesity. Peptides. 2020;133:170387. doi:10.1016/j.peptides.2020.170387

235. Patankar JV, Chandak PG, Obrowsky S, Pfeifer T, Diwoky C, Uellen A, Sattler W, Stollberger R, Hoefler G, Heinemann A, et al. Loss of intestinal GATA4 prevents diet-induced obesity and promotes insulin sensitivity in mice. Am J Physiol Endocrinol Metab. 2011;300(3):E478–E488. doi:10.1152/ajpendo.00457.2010

236. Yan Y, Shi R, Yu X, Sun C, Zang W, Tian H. Identification of atrial fibrillation-associated microRNAs in left and right atria of rheumatic mitral valve disease patients [published correction appears in Genes Genet Syst. 2019;94(4):181]. Genes Genet Syst. 2019;94(1):23–34. doi:10.1266/ggs.17-00043

237. Ozono R. New biotechnological methods to reduce oxidative stress in the cardiovascular system: focusing on the Bach1/heme oxygenase-1 pathway. Curr Pharm Biotechnol. 2006;7(2):87–93. doi:10.2174/138920106776597630

238. Qiu C, Wang Y, Zhao H, Qin L, Shi Y, Zhu X, Song L, Zhou X, Chen J, Zhou H, et al. The critical role of SENP1-mediated GATA2 deSUMOylation in promoting endothelial activation in graft arteriosclerosis. Nat Commun. 2017;8:15426. doi:10.1038/ncomms15426

239. Liu XM, Du SL, Miao R, Wang LF, Zhong JC. Targeting the forkhead box protein P1 pathway as a novel therapeutic approach for cardiovascular diseases. Heart Fail Rev. 2020;10.1007/s10741-020-09992-2. doi:10.1007/s10741-020-09992-2

240. Fischer A, Steidl C, Wagner TU, Lang E, Jakob PM, Friedl P, Knobeloch KP, Gessler M. Combined loss of Hey1 and HeyL causes congenital heart defects because of impaired epithelial to mesenchymal transition. Circ Res. 2007;100(6):856–863. doi:10.1161/01.RES.0000260913.95642.3b

241. Hecker M, Thamilarasan M, Koczan D, Schröder I, Flechtner K, Freiesleben S, Füllen G, Thiesen HJ, Zettl UK. MicroRNA expression changes during interferon-beta treatment in the peripheral blood of multiple sclerosis patients. Int J Mol Sci. 2013;14(8):16087–16110.. doi:10.3390/ijms140816087

242. Gu X, Liu Y, Wang N, Zhen J, Zhang B, Hou S, Cui Z, Wan Q, Feng H. Transcription of MRPL12 regulated by Nrf2 contributes to the mitochondrial dysfunction in diabetic kidney disease. Free Radic Biol Med. 2021;164:329–340. doi:10.1016/j.freeradbiomed.2021.01.004

243. Zhang D, Gu T, Forsberg E, Efendic S, Brismar K, Gu HF. Genetic and functional effects of membrane metalloendopeptidase on diabetic nephropathy development. Am J Nephrol. 2011;34(5):483–490. doi:10.1159/000333006

244. Bourgeois A, Bonnet S, Breuils-Bonnet S, Habbout K, Paradis R, Tremblay E, Lampron MC, Orcholski ME, Potus F, Bertero T, et al. Inhibition of CHK 1 (Checkpoint Kinase 1) Elicits Therapeutic Effects in Pulmonary Arterial Hypertension. Arterioscler Thromb Vasc Biol. 2019;39(8):1667–1681. doi:10.1161/ATVBAHA.119.312537

